# Loss of PIMREG impairs double-strand break signaling and repair

**DOI:** 10.1101/2025.10.15.682590

**Authors:** Maria Vitória de Rizzo Gasparini, Laís do Carmo, Giovanna Gonçalves de Almeida, Mariana Siqueira Lacerda Mamede, Isabela David Cardoso, Lígia Arantes Sardinha, Isabela Spido Dias, Jennifer Ann Black, Vanessa Cristina Arfelli, Yun-Chien Chang, Elza Tiemi Sakamoto Hojo, Stephanie Heinzlmeir, Bernhard Kuster, Philipp A. Greif, Leticia Fröhlich Archangelo

## Abstract

Glioblastoma (GBM) is the most common primary brain tumor in adults. It is an aggressive type of tumor with an average overall survival of 12-15 months post-diagnosis. Standard treatment involves surgical resection, followed by radiotherapy and chemotherapy with temozolomide (TMZ). However, resistance to treatment is frequently observed. PIMREG is a marker of proliferation highly expressed across different tumor types, with particularly elevated levels in GBM. Our previous study revealed that PIMREG silencing sensitizes GBM cells to TMZ treatment and affects ATM activation, a key apical kinase involved in DNA damage signaling. Given the emerging significance of PIMREG in the DNA damage response, we sought to explore PIMREG’s role in DNA damage repair and signaling in GBM cells exposed to genotoxic agents. Here we assessed the functionally enriched pathways associated with high *PIMREG* expression in GBM patient samples, analyzed the interactome of PIMREG upon treatment of GBM cells with TMZ and evaluated the impact of PIMREG loss on DNA damage signaling, DNA break formation and the repair of double-strand breaks. Our findings reveal that PIMREG functions as a direct substrate for ATM/ATR kinases, interacts with key DNA repair proteins upon TMZ treatment and actively facilitates the cellular response to chemotherapy– and radiotherapy-induced DNA damage. Taken together, PIMREG may function in the DNA damage response, particularly in the signaling and repair of double strand breaks via homologous recombination in GBM cells. In all, our data suggested high PIMREG expression may affect resistance of GBM cells to treatment.

## 1. Introduction

Diffuse gliomas are the most common primary malignant brain tumors [1]. Among gliomas, GBM grade 4 is the most aggressive subtype, characterized by rapid progression and poor prognosis [2,3]. The median survival time for patients with GBM is approximately 12 to 15 months following diagnosis, with a 5-years survival rate of 9.8 % [4]. The standard treatment consists of surgical resection followed by radiotherapy with ionizing irradiation (IR) and chemotherapy with Temozolomide (TMZ) [5].

Treatment with IR and TMZ is based on inducing DNA damage in tumor cells, which promotes cytotoxicity and cell death. Whereas IR mostly induces double-strand breaks (DSBs) in the DNA through radiolysis and formation of reactive oxygen species (ROS) [6], TMZ primarily alkylates DNA bases. O6-methylguanine (O6-MeG), most cytotoxic lesion is repaired by the direct repair mechanism mediated by the O6-methylguanine-DNA methyltransferase (MGMT) [7]. Thus, hypermethylation of the promoter region of the *MGMT* gene predicts response to TMZ treatment [8]. In this scenario, inefficient repair of O6-MeG leads to futile cycles of mismatch repair (MMR), replication fork collapse and ultimately DSB formation [9].

DSBs are highly cytotoxic DNA lesions that threaten genome stability if unresolved. In response, eukaryotic organisms possess a multitude of pathways orchestrated through a network of proteins to tackle these severe lesions. DSBs however, are primarily resolved through the activation of two main pathways: homologous recombination (HR) [10] and non-homologous end joining (NHEJ) [11]. Overcoming DSB lesions through DNA repair is a critical mechanism for acquiring resistance to therapy. In fact, elevated activities of HR and NHEJ are observed in patient-derived GBM samples with acquired TMZ resistance, suggesting that these pathways play a central role in the development of treatment resistance [12].

PICALM-interacting mitotic regulator (*PIMREG*) plays a well-established role in cell cycle progression, specifically in proliferation and chromosome segregation during cell division [13–15]. Additional functions have been proposed for PIMREG in the developing central nervous system (CNS) [13], cell differentiation [15–17], fetal cardiomyocyte cell cycle progression [18,19], and in signaling pathways such as JAK-STAT3 [17], NF-κB [20], and βCcatenin [21,22]. PIMREG expression levels strongly correlate with cellular proliferation in both normal and malignant cells [13].

The contribution of PIMREG in promoting malignant phenotypes has been primarily addressed in breast cancers, where multiple functional studies have linked PIMREG to cell migration, metastasis, epithelial-mesenchymal transition, enhanced stemness features, and aggressiveness of breast cancer cells [20,22,23]. In another hormone-sensitive cancer, such as prostate cancer (PCa), PIMREG was described as an oncogenic factor whose expression became activated through direct binding of the androgen receptor (AR) to the *PIMREG* promoter, leading to proliferation, colony formation, migration, invasion, and cell cycle progression of PCa cells [24].

The oncogenic role of PIMREG in cancer development has been shown *in vivo* by three elegant studies. Jiang and colleagues demonstrated that overexpression of PIMREG promotes breast cancer aggressiveness via constitutive activation of NF-κB signaling pathway. Mechanistically, PIMREG overexpression sustains constitutive activation of NF-κB signaling by competing with IkB for interaction, thereby disrupting the NF-κB/IκBα negative feedback loop. In cells expressing high levels of PIMREG, such as breast cancer cells, PIMREG interacts with the p65 subunit of NF-κB within the nucleus, thereby stabilizing p65, and enhancing its binding to NF-κB response elements, promoting activation of its target genes and consequently hyperactivation of NF-κB signaling that correlates with cancer development and progression [20].

The PIMREG-mediated activation of the NF-κB pathway is also implicated in promoting cisplatin resistance in triple-negative breast cancer (TNBC) both *in vivo* and *in vitro* [25]. For instance, SOD1-high fibroblasts produce exosomal miR-3960 that targets the kinase BRSK2 in TNBC cells. BRSK2 is responsible for phosphorylating PIMREG at Ser16 leading to PIMREG ubiquitination and subsequent degradation. Due to miR-3960-mediated downregulation of BRSK2, PIMREG Ser16 phosphorylation becomes reduced thereby increasing protein stability. Consequently, NF-κB signaling is activated correlating with cisplatin resistance in TNBC cells. Conversely, decreased PIMREG levels due its phosphorylation of Ser16, enhances cisplatin sensitivity in TNBC cells [25]. In a third study, Xu and colleagues revealed PIMREG’s critical role in inflammation-associated tumorigenesis. Mice lacking *Pimreg* exhibited impaired differentiation of Th17 lymphocytes, leading to reduced proinflammatory activity typically associated with these cells. PIMREG is essential for IL-6–mediated activation of STAT3 and regulates Th17 cell differentiation by modulating STAT3 transcriptional activity. As a result, PIMREG plays a key role in the development of Th17-driven inflammatory diseases, including autoimmune encephalomyelitis, colitis and colitis-associated carcinogenesis in animal models [17].

Consistent with its role as a proliferation marker, *PIMREG* is highly expressed in tumor tissues compared to normal tissue samples. Additionally, high expression of *PIMREG* in patient samples correlated with worse prognosis and reduced overall survival across various types of cancer [26,27]; including breast cancer [20,22,23,28]; pancreatic cancer [29]; lung cancer [30]; prostate cancer [24]; head and neck squamous cell carcinoma [31]; clear cell renal cell carcinoma [32]; gastric cancer [33]; melanoma [34] and gliomas [35]. We have previously shown that GBM exhibits the highest level of *PIMREG* expression amongst 25 types of cancer. As well as showing that PIMREG silencing reduced malignant characteristics of GBM cells, such as cell cycle progression, proliferation and migration capacity, we demonstrated that treating these cells with TMZ increased the levels of PIMREG protein. In addition, PIMREG silencing rendered GBM cells more sensitive to TMZ exposure and interfered with ATM activation, a key apical kinase that orchestrates the cellular response to DSBs. We hypothesized that PIMREG plays a role in DNA damage response (DDR) and therefore could contribute to TMZ resistance [35].

The mechanism by which PIMREG participates in the DDR in GBM cells is unclear. To begin addressing these knowledge deficits and gain further insight into PIMREG function and relevance in GBM, we tested for functionally enriched pathways related to elevated *PIMREG* expression in GBM patient samples and analyzed the correlation between *PIMREG* expression and those of DDR-related genes. We investigated the interactome of PIMREG upon treatment of GBM cells with TMZ and evaluated the impact of PIMREG loss on DNA damage signaling, DNA break formation and the repair of DSBs. Our data reveal that PIMREG interacts with proteins involved mainly in DDR and RNA metabolism following TMZ treatment in GBM cells. Additionally, PIMREG serves as a substrate for ATM/ATR and plays a role in DNA damage signaling and the repair of DSBs. These findings further suggest the importance of elevated expression of PIMREG in GBM maintenance and its possible contribution to the treatment resistance in GBM patients.

## 2. Material and methods

### 2.1. Public data analysis

GBM RNAseq-based mRNA expression (RSEM) data were downloaded from TCGA-PanCancer Atlas_2018 (592 samples) using cBioPortal for Cancer Genomics [36]. Gene set enrichment analysis was performed using GSEA version 4.3.2 [37]. Results were regarded as significant with a *p*-value < 0.05 and 25 % false discovery rate (FDR) threshold (*q*-value <0.25).

Genes coding for DDR proteins, whose expression covariates with *PIMREG* in GBM patient samples were plotted in a correlation curve using GraphPad Prism 9. *PIMREG* mRNA expression data, along with *MGMT* promoter methylation status, were also obtained from the merged cohorts of Brain Lower Grade Glioma (LGG) (TCGA, PanCancer Atlas, 514 samples) and GBM (TCGA, PanCancer Atlas, 592 samples). Among the 1106 available samples, 635 contained information regarding the methylation status of the *MGMT* promoter.

### 2.2. Cell lines

The GBM human cell line T98G was obtained from the American Type Culture Collection (ATCC, Manassas, USA) and U87MG from the Banco de Células do Rio de Janeiro (BCRJ, Rio de Janeiro, Brazil). The reporter cell lines HeLa EJ5-GFP and U2OS DR-GFP were kindly provided by Dr. Valeria Valente (School of Pharmaceutical Science, UNESP, Brazil). The cells were cultured in Dulbecco’s Modified Eagle’s Medium (DMEM, Gibco) supplemented with 10 % of fetal bovine serum (FBS, Gibco) and 1 % of the antibiotic penicillin/streptomycin (Gibco), at 37 °C and 5 % of CO_2_. All cell lines were tested for mycoplasma with MycoAlert PLUS Mycoplasma Detection Kit (Lonza). T98G were authenticated by STR matching analysis using the GenePrint 10 system (Promega, Madison, WI, USA) and 3730 DNA Analyzer – Applied Biosystems (Applied Biosystems, Foster City, CA, USA).

### 2.3. Cell treatment with genotoxic agents

Temozolomide (TMZ, Sigma-Aldrich # T2577 and Selleckchem S1237) were diluted in DMSO at a stock concentration of 50 mM and 200 mM, respectively. Stocks were stored in 500 µL aliquots at – 80 °C.

GBM cells were treated with 300 µM TMZ or vehicle (DMSO) for 24 h, for Western blot analysis, cell cycle, apoptosis, immunoprecipitation and immunofluorescence experiments.

For IR treatment, GBM cells were irradiated using the PRIMUS linear accelerator (Siemens Healthineers, Brazil) with a 6 MV photon beam at the Hospital das Clínicas – Ribeirão Preto Medical School. The administered dose was 4 Gy. Cells were collected 0.5 h and 1 h post-irradiation for Western blot analysis and immunofluorescence experiments. The concentration and time used for the TMZ and IR treatments were defined as such to induce DNA damage signaling but not cell cycle arrest or apoptosis. Cell cycle progression and viability were monitored in all GBM cell samples treated with genotoxic agents (Supplementary Fig. S1C, S1D and Fig. S2D).

### 2.4. Plasmid construction

For genome editing, single guide RNAs (sgRNAs) targeting exon 2 of *PIMREG* were designed using the Benchling CRISPR sgRNA design tool (https://www.benchling.com/crispr). The 20 bp oligos contained the restriction sites CACC and AAAC at the 5’end of the sense and antisense oligos, respectively, as described in [38]. Annealed oligos were cloned into the PX458 vector (Addgene plasmid #48138) digested with the *BpiI* restriction enzyme (Thermo Fisher Scientific) generating the PX458-PIMREG-KO vectors (KO-1, KO-2, KO-3 and KO-4). The target sequences are as follows: sgRNA-1 – sense: GAGAGATCTCCGGCGCACGG and antisense: CCGTGCGCCGGAGATCTCTC; sgRNA-2 – sense: CTTCTCGGTGGCAGAACATG and antisense: CATGTTCTGCCACCGAGAAG; sgRNA-3 – sense: CCATCAGGAGACCTCTGTAG and antisense: CTACAGAGGTCTCCTGATGG; sgRNA-4 – sense: TCAACCTCAACCTCCGCGCA and antisense: TGCGCGGAGGTTGAGGTTGA.

For ectopic expression of PIMREG, the pMIT-PIMREG-HA construct was generated. Briefly, the PIMREG hemagglutinin (HA)-epitope tagged cDNA was subcloned into the *XhoI* and *NotI* restriction sites of the pMIG-AEfl-TOM vector backbone, upstream of the internal ribosomal entry site (IRES) and the tdTomato reporter gene.

### 2.5. Generation of CRISPR/Cas9 edited monoclonal cell lines

The GBM T98G cells and the reporter cell lines HeLa EJ5-GFP and U2OS DR-GFP were edited to generate PIMREG knockout cell lines (PIMREG KO-1, KO-2, KO-3, KO-4). 2 – 3 x 10^5^ cells were seeded at 6-well plates and transfected with 2.5 µg of PX458-PIMREG-KO vectors and Lipofectamine 2000 according to manufacturer’s instructions (ThermoFisher). Efficiently transfected cells were single sorted by Fluorescence-Activated Cell Sorting (FACS) based on GFP expression. The individual cells were cultured in 96 well plates for clonal expansion and generation of the monoclonal cell lines. PIMREG KOs were confirmed by Western blot analysis and genomic sequencing of the edited *PIMREG* locus.

The presence of phenotypic variability among the different PIMREG KO clones, is a common limitation inherent to studies relying on single-cell CRISPR/Cas9-generated KOs. Particularly intriguing, was the unexpected re-expression of PIMREG protein after extended culture of the GBM T98G PIMREG KO-2 cell line. This finding prompted us to routinely verify PIMREG absence by immunofluorescence in all KO cell lines throughout the course of the experiments. Importantly, the PIMREG KO-2 cell line that recovered PIMREG protein expression (KO-2 recPIMREG) was subsequently used as a control in experiments analyzing γH2AX fluorescence intensity in PIMREG KO and WT cells following IR exposure.

### 2.6. Cell cycle analysis

1 x 10^6^ cells were fixed in 70 % ethanol for 30 min on ice, washed with PBS, treated with RNAse A solution (100 µg/ml) and stained with 100 µg/ml propidium iodide (PI). Cell cycle analysis was performed using FACS Symphony A1 (BD Biosciences) and FlowJo software version 10. Statistical analyses were performed with Two-way ANOVA, using GraphPad Prism 9.

### 2.7. Cell viability analysis

Cell viability was assessed with either Annexin V or Zombie NIR staining. For the Annexin V staining, cells were washed with the Annexin V Binding Buffer (BD Pharmingen™) and incubated with 5 µl Annexin V reagent (BD Pharmingen™) for 15 min at RT followed by 2.5 µL of PI solution (Miltenyi Biotec) (100 µg/mL). For the Zombie NIR staining, cells were incubated with Zombie NIR reagent (BioLegend) (1:400) for 15 min, washed with PBS, fixed with 2 % paraformaldehyde (PFA) for 15 min and resuspended in 1 % glycine/PBS. Cell viability was analyzed using FACS Symphony A1 (BD Biosciences) and FlowJo software version 10.

### 2.8. Immunoblotting and Immunoprecipitation

Cells were lysed in lysis buffer with protease inhibitors (10 mM EDTA, 100 mM Tris-HCl pH 7.5; 1 % Triton X-100, 10 mM Sodium Pyrophosphate (Na_2_P_2_O_7_), 100 mM Sodium Fluoride (NaF), 10 mM Sodium Orthovanadate (Na_3_VO_4_); 2 mM PMSF). Protein extracts were separated by electrophoresis on 12 % or on a gradient 2.5-12.5 % SDS polyacrylamide (SDS-PAGE) gel and transferred to a nitrocellulose membrane (GE Healthcare Life Science). Membranes were blocked with 1 % BSA or 5 % low fat milk and probed with a specific antibody, followed by incubation with a secondary antibody conjugated to horseradish peroxidase (HRP). Protein signals were detected by chemiluminescence and the images were captured by ChemiDoc™ Imaging System (Bio-Rad). Primary antibodies were: anti-PIMREG (anti-CATS 2C4) (1:10) [13], anti-β-Actin (ThermoFisher Scientific, # AM4302 and Santa Cruz Biotechnology sc-1616) (1:10000 and 1:2000, respectively), anti-α-tubulin (Santa Cruz Biotechnology sc-5286) (1:1000), anti-phospho-H2AX (S139) (Cell Signaling Technology, #2577 and #80312) (1:500 and 1:1000, respectively), anti-phospho-ATM (S1981) (Sigma-Aldrich, clone 10H11.E12) (1:1000), anti-ATM (Sigma-Aldrich, clone MAT3-4G10/8) (1:1000), anti-phospho-DNA-PKcs (S2056) (Cell Signaling Technology, #68716) (1:1000), anti-DNA-PKcs (Cell Signaling Technology, #38168) (1:1000), anti-phospho-CHK1 (S345) (Cell Signaling Technology, #2348) (1:1000), anti-CHK1 (Cell Signaling Technology, #2360) (1:1000), anti-phospho-KAP1 (S824) (Abcam, ab243870) (1:1000), anti-KAP1 (Abcam, ab109545) (1:1000) and anti-phospho-ATM/ATR substrate (Phospho-(Ser/Thr) ATM/ATR Substrate; Cell Signaling Technology, #2851) (1:1000). The secondary antibodies were anti-mouse (Santa Cruz Biotechnology, sc-2064) (1:3000), anti-rat (KPL, 52200364) (1:3000), anti-goat (Santa Cruz Biotechnology, sc-2354) (1:3000) and anti-rabbit (Abcam, ab6721) (1:3000). The blots were stripped with Restore Western Blot Stripping Buffer (Thermo Scientific, YE366939) before reprobing. Quantitative analyses of the optical intensities of the protein bands were determined with UN-SCAN-IT gel 6.1 software (Silk Scientific Inc., UT) and normalized by control protein.

For immunoprecipitation, 170 – 290 µg of cellular extracts were incubated overnight with 50 µl of monoclonal rat anti-PIMREG (anti-CATS 2C4), 1:20 (v/v) rabbit anti-phospho-ATM/ATR substrate antibodies and respective isotype controls (1.6 μg of rat IgG Santa Cruz Biotechnology, sc-2026 and 21 µg of rabbit IgG, Jackson ImmunoResearch Laboratories, Inc., 011-000-003) at 4 °C, under rotation. The next day, 40 μl of Protein G-Agarose (Roche) was added and incubated further for 3 h. Beads were extensively washed with lysis buffer (50 mM Tris-HCl pH 8, 150 mM NaCl, 0.1 % SDS, 1 % IGEPAL, and 0.5 % sodium deoxycholate), resuspended in SDS sample buffer, boiled, and separated on a 12 % SDS-PAGE gel. Subsequent Western blot analysis was carried out with the antibodies anti-PIMREG (anti-CATS 2C4) and anti-phospho-ATM/ATR substrate.

### 2.9. Immunofluorescence

Cells grown on coverslips were treated with TMZ and IR, fixed and permeabilized by sequential incubation in 1 % PFA/ 0.5 % Triton X-100 followed by 1 % PFA/ 0.3 % Triton X-100 and 0.5 % methanol for 20 min and blocked for 1 h with blocking solution (0.5 % Triton X-100, 1 mM EDTA, 1 mg/mL BSA, and 0.03 % FBS) at RT. Coverslips were incubated for 1 h with the primary antibodies diluted in blocking solution: anti-PIMREG (Anti-CATS-2C4) (1:10), anti-phospho-H2AX (S139) (Cell Signaling Technology, #80312) (1:100), anti-RAD51 (D4B10) (Cell Signaling Technology, #8875) (1:50), YY1 Recombinant Superclonal™ Antibody (Thermo Fisher Scientific, #712089) (1:400) and PRMT5 Antibody (A-11) (Santa Cruz, sc-376937) (1:100). Secondary Alexa Fluor-conjugated antibodies Alexa Fluor anti-rat 488 (Invitrogen, A-11006) (1:500), Alexa Fluor anti-mouse 488 (Invitrogen, A-11006) (1:500), Alexa Fluor anti-mouse 555 (Invitrogen, A21325) (1:600), Alexa Fluor anti-rat 555 (Invitrogen, A21434) (1:500), Alexa Fluor anti-rabbit 594 (Invitrogen, A-11012) (1:500), and Alexa Fluor anti-rabbit 594 (Thermo Fisher Scientific, A32754) (1:600) were used for detection of the primary antibodies. Nuclei were counterstained with 300 nM DAPI solution and coverslips were mounted on slides using Fluoromount-G™ Mounting Medium (Invitrogen).

Images were acquired using a Leica SP5 confocal microscope and a Leica DMI6000B epifluorescence microscope equipped with LAS AF software, employing 100x magnification in z-stack mode. γH2AX fluorescence intensity was quantified using ImageJ Fiji software [39]. Corrected Total Cell Fluorescence (CTCF) was calculated for each nucleus (CTCF = IntDen – (mean background × area)). A minimum of 75 cells were analysed per condition per experiment.

RAD51 images were captured using a Leica DMI6000B epifluorescence microscope, with identical acquisition settings maintained within each experiment. Approximately 10-16 random fields per condition were captured using a 100x oil immersion objective. Analysis of RAD51 foci was performed using Cell Profiler software [40]. Data from WT and each KO clone represent one or two technical replicates per condition in each experiment, across a total of three independent biological experiments. A minimum of 60 cells were analysed per experimental condition. Graphs and statistical analyses were performed using GraphPad Prism v.10.0. Images were processed using ImageJ Fiji, with brightness and contrast equally enhanced across each image to improve visualisation.

For colocalization, images were captured using a Leica Stellaris 5 confocal microscope. Line-scan analysis was performed using the “plot profile” tool in ImageJ Fiji. Fluorescence intensity values (gray values) across distance (µm) for each channel were plotted as line graphs in GraphPad Prism 9. Colocalization analysis was performed using Leica Application Suite X (LAS X, Leica Microsystems). Fluorescence images, acquired under identical settings were analyzed using the colocalization module. Regions of interest (ROIs) were defined, and background signal was minimized by applying intensity thresholds. The degree of colocalization between channels was quantified using Pearson’s correlation coefficient (*r*). All analyses were performed on raw images without further manipulation unless otherwise stated.

### 2.10. Comet assay

DNA breaks were assessed using the alkaline comet assay (single-cell gel electrophoresis). T98G WT and PIMREG KO cells were seeded in 24-well plates at a density of 8 x 10C cells per well and treated with TMZ or DMSO. Methyl methanesulfonate (MMS) was used as a positive control for DNA damage (300 µM for 1h). After treatment, cells were harvested, resuspended in 0.5 % low-melting-point agarose at 37 °C, and spread onto microscope slides pre-coated with 1.5 % agarose. Slides were incubated at 4 °C for 5 min to allow agarose solidification and then immersed in lysis buffer (2.5 M NaCl, 100 mM EDTA, 10 mM Tris, 200 mM NaOH, 1 % Triton X-100, pH 10) for at least 12 h at 4 °C. Slides were washed in PBS for 5 min at 4 °C and incubated in alkaline electrophoresis buffer (300 mM NaOH, 1 mM EDTA, pH 13) for 20 min at 4 °C to allow DNA unwinding. Electrophoresis was performed at 25 V and 300 mA for 20 min, followed by neutralization in 0.4 M Tris (pH 7,5) for 15 min at RT. Slides were air-dried, fixed in absolute ethanol for 3 min, stained with SybrGreen I (Sigma Aldrich) and analyzed using Comet Assay IV software (Instem, USA). All steps were performed in the dark.

### 2.11. DNA repair assay

The frequency of HR and NHEJ were estimated using U2OS DR-GFP and HeLa EJ5-GFP reporter cells [40,41]. 3 x 10^5^ HeLa EJ5-GFP and 1 x 10^5^ U2OS DR-GFP cells WT and PIMREG KOs (KO-1, –2, –3 and –4) were seeded at 6-well plates and transfected with 2.5 µL of the *I-SceI* endonuclease expressing plasmid, which creates the DSBs (pCBASceI; Addgene Plasmid #26477). Cells were collected 48 h after transfection and the percentage of GFP-positive cells was determined by flow cytometry using FACS Symphony A1 or Accuri C6 (BD Biosciences) and FlowJo software. Dead cells were excluded from the analysis using Zombie NIR (BioLegend) staining at proportion 1:400. Fifty thousand events were acquired. The percentage of GFP-positive cells of each sample was normalized by the mean value of WT GFP-positive cells used as internal control for each experiment.

For rescue experiments, the pMIT-PIMREG-HA construct (or the corresponding backbone plasmid as a control) was co-transfected with the *I-SceI* endonuclease-expressing plasmid. The percentage of GFP-positive cells was then quantified specifically within the tdTomato-positive population (GFP^+^/tdTomato^+^ double-positive cells). Fluorescence signals were acquired using FITC (GFP), APC-Cy7 (tdTomato), and PE-CF594 (Zombie viability dye) detection channels.

### 2.12. Mass spectrometry analysis

PIMREG was immunoprecipitated from cellular extracts of U87MG (900 µg) and T98G (500 µg) cells treated with TMZ and DMSO using the anti-PIMREG (anti-CAST2C4) antibody and protein G magnetics beads (PureProteome™ Protein G Magnetic Beads, Merck). Normal rat IgG antibody (Santa Cruz Biotechnology) was used as isotype control. The experiments were performed in triplicates (Supplementary Fig. S1).

Immunoprecipitated proteins were reduced with 10 mM dithiothreitol (DTT) at 37 °C for 45 min, then alkylated with 55 mM chloroacetamide at RT for 30 min in the dark. Samples were mixed with NuPAGE LDS sample buffer (Thermo Fisher Scientific), incubated at 70 °C for 10 min, and loaded onto a NuPAGE Bis-Tris gel by electrophoresis at 200 V for 5 min, allowing proteins to migrate approximately 1Ccm into the gel. Gels were fixed in 2 % acetic acid and 40 % methanol and stained with Coomassie blue. Gel bands were excised, transferred to a 96-well plate, and destained with 50 mM triethylammonium bicarbonate (TEAB) in 50 % ethanol for 2 h at RT followed by 1 h at 55 °C. Gel pieces were dehydrated with 100 % ethanol for 10Cmin, rehydrated in 5 mM TEAB for 20 min, and dehydrated again with 100 % ethanol for 10Cmin.

For digestion, 20 μl of trypsin solution (10 ng/μl in 5 mM TEAB) was added to each gel piece and incubated at 4 °C for 15 min. Excess of trypsin solution was removed and replaced with 20 μl of 5 mM TEAB, and digestion was performed overnight at 37 °C. Peptides were extracted by sequential incubations: two rounds with 20 μl of 1 % formic acid (FA) for 30 min each, followed by extraction with 20 μl of 60 % acetonitrile (ACN) containing 0.1 % FA for 30 min. Extracted peptides were dried and stored at – 20C°C until LC-MS analysis.

Peptides were analyzed using an Orbitrap Fusion Lumos mass spectrometer (Thermo Fisher Scientific) coupled to a Dionex UltiMate 3000 RSLCnano system (Thermo Fisher Scientific). Peptide separation was performed at a flow rate of 300CnL/min on a 50-min linear gradient from 4% to 32% solvent B (0.1 % FA and 5 % DMSO in ACN) in solvent A (0.1 % FA and 5 % DMSO in water). The MS was operated in positive ion mode using data-dependent acquisition (Top20 method). Full MS scans (MS1) were acquired in the Orbitrap from 360 to 1300Cm/z at a resolution of 60,000 in profile mode, with an automatic gain control (AGC) target of 4e5 and a maximum injection time (maxIT) of 50Cms. For MS/MS (MS2) acquisition, precursor ions were isolated with a 1.7CTh window, fragmented by higher-energy collisional dissociation (HCD) at a normalized collision energy (NCE) of 30 %, and analyzed in the Orbitrap at a resolution of 30,000 in centroid mode (AGC target of 2e5 and maxIT of 50 ms). Dynamic exclusion was set to 20Cs.

MS raw files were processed using MaxQuant (version 1.6.17.0) and searched against the UniProtKB human reference proteome (downloaded on 2019-07-15) with default parameters. Carbamidomethylation of cysteine was set as a fixed modification, and oxidation of methionine as well as N-terminal protein acetylation were defined as variable modifications. Protein groups labeled as “reverse” or “potential contaminant” were excluded. Protein abundance was estimated based on iBAQ values, and missing values were imputed with a constant value corresponding to a log2 iBAQ intensity of 10 to enable downstream statistical analysis. Differential expression analysis was performed using a two-sample t-test, and p-values were adjusted for multiple testing using the Benjamini-Hochberg procedure.

The comparison between PIMREG immunoprecipitates vs. IgG under vehicle (DMSO) conditions (PIMREG IP vs. IgG) identified the PIMREG-interacting partners (i.e. significantly upregulated proteins), considered as the *bona fide* PIMREG interactome. The comparison between PIMREG immunoprecipitates from TMZ vs. vehicle (TMZ vs. DMSO) revealed TMZ-induced alterations in PIMREG interactions and gave rise to the TMZ-regulated interactome. The identified proteins were filtered by significance based on log_2_ fold change cutoff > 1 (PIMREG IP vs. IgG) and log_2_ fold change cutoff > 1 and < –1 (TMZ vs. DMSO) and adjusted *p*-value < 0.05. Data was analyzed using R software (version 4.3.2) and Excel (Microsoft Office).

Upregulated proteins in the TMZ-regulated interactome are proteins that interact with PIMREG only under TMZ treatment or show increased association under treatment whereas the downregulated proteins are those with reduced or lost binding upon TMZ treatment.

### 2.13. Gene Ontology and network analyses

Gene Ontology (GO) analysis was carried out using the EnrichR platform (https://maayanlab.cloud/Enrichr/). Terms with adjusted *p*-value < 0.05 were considered statistically significant.

PIMREG and its interacting proteins were searched for interaction networks using the STRING database (https://string-db.org). Interactions determined experimentally, registered in curated databases, extracted from text mining or coexpression data were considered, using a medium confidence (0.400) interaction score. Network was clustered using the MCL clustering algorithm, with an inflation parameter of 3.

## 3. Results

### 3.1. GSEA analysis implicates *PIMREG* in DNA repair and splicing

*PIMREG* expression correlates with genes associated with cell cycle, DNA replication and DNA repair in GBM [35]. To dissect the relationship between high *PIMREG* expression and GBM further, we performed a GSEA analysis testing for functionally enriched pathways that correlate with high expression of *PIMREG* in GBM patient samples (TCGA, PanCancer Atlas, 592 samples). Our analysis revealed 392 enriched gene sets related to high *PIMREG* expression using the HALLMARK (17 gene sets), KEGG (65 gene sets) and REACTOME (310 data sets) databases (Supplementary Table 1). In support of previous results [35], our analysis uncovered correlations between high *PIMREG* expression and the gene sets from several DNA repair pathways (33 out of 392 gene sets), including base excision repair (BER), nucleotide excision repair (NER), mismatch repair (MMR), double-strand break repair (DSB repair), homology directed repair (HDR), Fanconi anemia (FA), and UV-response (Fig. 1A, Supplementary Table 2), indicating putative functions for PIMREG across multiple DNA repair pathways in GBM.

**Fig. 1:**
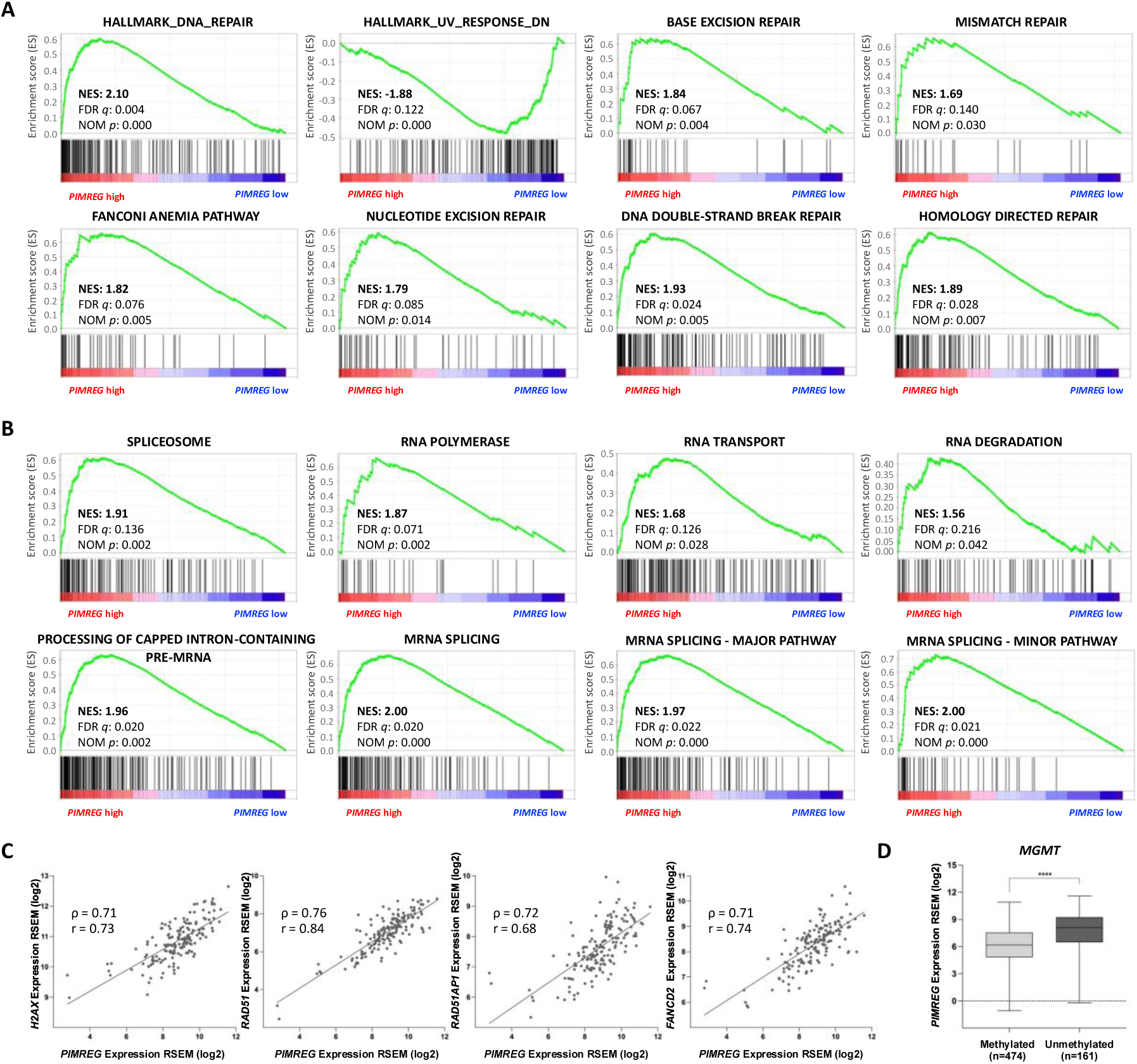
GBM samples with *PIMREG* high expression are enriched for DDR and RNA metabolism. (**A**) GSEA analysis reveals enriched gene sets related to DDR and (**B**) RNA metabolism. (**C**) Correlation plot between the transcript levels of *PIMREG* and the DDR coding genes, *H2AX*, *RAD51*, *RAD51AP1* and *FANCD2* in GBM patient samples. (ρ) Spearman’s correlation coefficient and (r) Pearson’s correlation coefficient. (**D**) *PIMREG* expression in glioma patients classified according to methylation status of the *MGMT* promoter region. (****) *p*-value < 0.0001, Mann-Whitney two-tailed test. RSEM: RNAseq by Expectation-Maximization.

Moreover, gene sets pertaining to RNA metabolism, such as spliceosome, RNA polymerase, RNA transport, RNA degradation, processing of capped intron-containing pre-mRNA and mRNA splicing were also abundant, accounting for 12.5 % of all gene sets (49 out of 392), suggesting an unforeseen function for *PIMREG* in RNA splicing and processing (Fig. 1B, Supplementary Table 3).

In line with putative functions for PIMREG in DDR, we performed a correlation analysis using data from the GBM patient samples uncovering a strong positive correlation between *PIMREG* transcript abundance and the abundance of transcripts from essential genes required for DSBs recognition and repair, including *H2AX*, *RAD51*, *RAD51AP1*, and *FANCD2* (Fig. 1C).

Since the methylation status of the *MGMT* promoter is a well-established predictive biomarker for response to TMZ treatment in gliomas [42], *PIMREG* expression was additionally assessed in glioma patient samples stratified by the methylation status of the *MGMT* promoter. The analysis revealed that samples with unmethylated *MGMT* promoters, (MGMT proficient) possessed significantly higher levels of *PIMREG* expression compared to samples with methylated *MGMT* promoters (Fig. 1D), suggesting that PIMREG is more abundant in the group of patients with lower response to TMZ. Altogether, these findings support our previous analyses, suggesting wide roles for PIMREG during the DDR but also in RNA metabolism in GBM.

### 3.2. PIMREG interactome in response to TMZ treatment

We have previously shown that treating GBM cells with TMZ triggers an increase in PIMREG protein levels, as cells accumulate DNA lesions. We hypothesized that PIMREG accumulation is required to mitigate DNA lesions induced by TMZ treatment [35].

To gain insight into the functional implications of PIMREG accumulation in response to TMZ treatment, PIMREG was immunoprecipitated from GBM cells treated with TMZ and its interactome identified by mass spectrometry (Supplementary Fig. S1).

To ensure reproducibility, only the proteins with peptides detected in all three replicates and filtered by significance based on log_2_ fold change cutoff > 1 (PIMREG IP vs. IgG) or log_2_ fold change cutoff > 1 and < –1 (TMZ vs. DMSO) and adjusted *p*-value < 0.05, were considered and identified as PIMREG interactors.

In U87MG cells, a total of 36 proteins were identified as PIMREG interacting partners (Fig. 2A and Supplementary Table 4), whereas 29 proteins were identified as TMZ-regulated PIMREG interactors (Fig. 2B and Supplementary Table 5). In T98G cells, 8 proteins were identified as PIMREG interacting partners (Fig. 2C and Supplementary Table 6) while 4 were identified as TMZ-induced interacting proteins (Fig. 2D and Supplementary Table 7). To better understand the molecular functions of PIMREG, we analyzed its protein interaction network using the STRING database. Both upregulated and downregulated interacting proteins were used for this analysis. This approach strongly implicated PIMREG in splicing and ribosome biology, as protein clusters related to these functions were identified in both PIMREG interactome and TMZ-regulated interactome of U87MG cells, albeit with distinct sets of interactors. Additionally, a cluster of proteins related to NADH dehydrogenase activity and one unidentified cluster of PIMREG and TRIP13 were observed in the TMZ-regulated interactome only, whereas a cluster related to immunoglobulin complex was observed in the PIMREG interactome only (Fig. 2E and 2F). For the T98G cells, a cluster of proteins related to the Polycomb complex was observed within the TMZ-regulated PIMREG interactors (Fig. 2G), whilst no cluster was formed among the proteins identified in the PIMREG interactome.

**Fig. 2:**
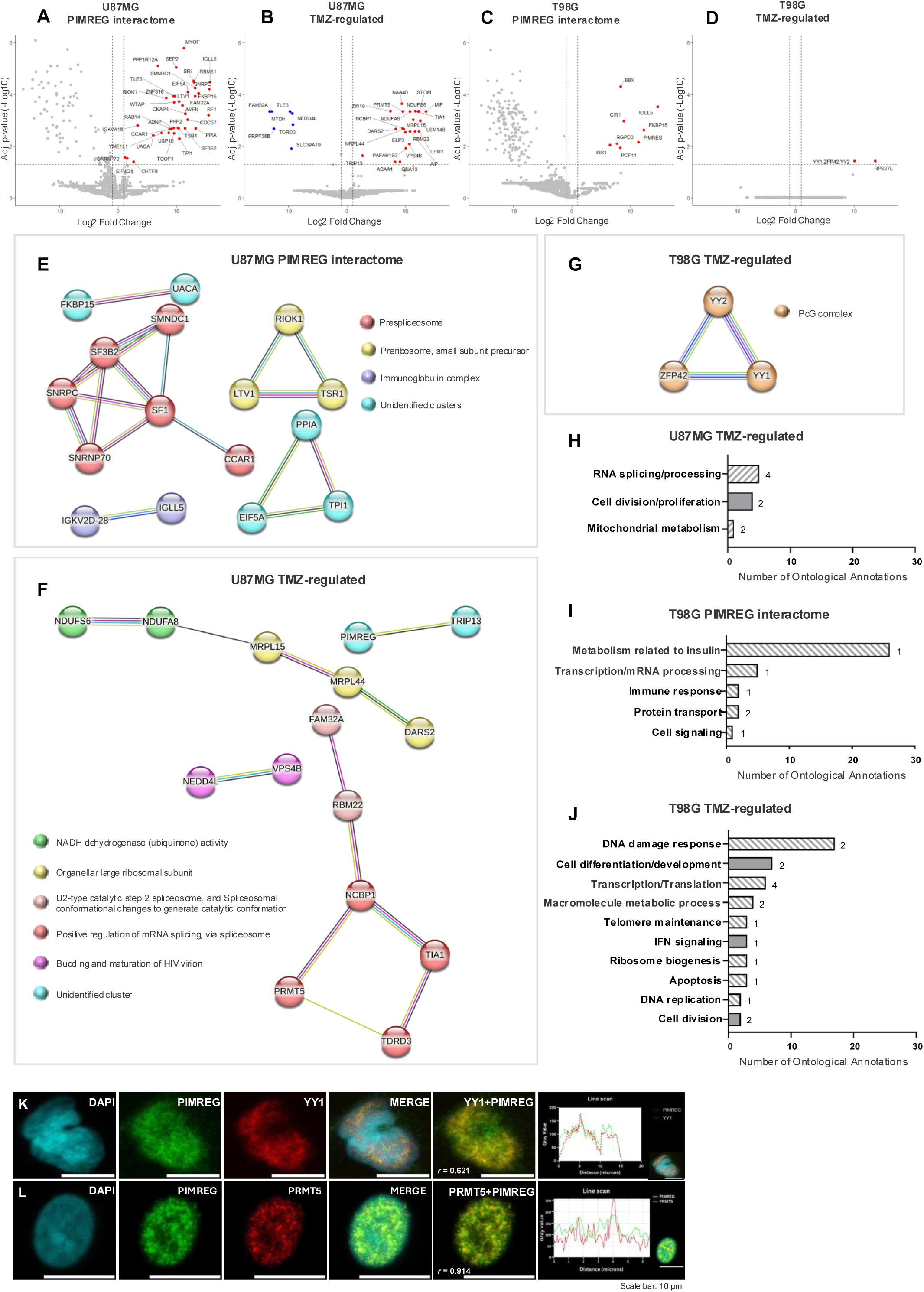
PIMREG interactome of GBM cells treated with TMZ and vehicle (DMSO). (**A-D**) Volcano plot graphs of PIMREG interacting proteins in U87MG (A, B) and T98G (C, D) cells. (**B, D**) TMZ-regulated PIMREG interactors (TMZ vs. DMSO). Upregulated proteins (red dots) interact with PIMREG only under TMZ treatment or show increased association under treatment. Downregulated proteins (blue dots) exhibit reduced or lost binding upon TMZ treatment. (**A, C**) PIMREG interacting proteins under vehicle (DMSO) conditions (PIMREG IP vs. IgG). Proteins with adjusted *p*-value < 0.05 and log_2_ fold change > 1 (PIMREG IP vs. IgG) or log_2_ fold change < –1 and > 1 (TMZ vs. DMSO) were considered significant (highlighted as blue or red dots). (**E**) Interaction network of proteins identified as PIMREG interactors in U87MG cells (PIMREG IP vs. IgG). (**F**) Interaction network of the proteins identified as TMZ-regulated PIMREG interactors in U87MG and (**G**) in T98G cells. Each node represents one protein. The proteins interacting with each other are represented in clusters. Node colors indicate proteins that have the same function within the clusters. Lines connecting the nodes represent known interactions (cyan: from curated databases; magenta: experimentally determined), predicted interactions (green: gene neighborhood; red: gene fusions; blue: gene co-occurrence) and other types of interactions (black: co-expression; yellow: text mining; light blue: protein homology). (**H, I, J**) Biological processes (BP) in each group of PIMREG interacting proteins. Number of ontological terms grouped as a BP (graph bars) is depicted in the x axis. Number of genes involved in each BP is shown next to each bar. The bars in gray represent previously described functions of PIMREG. The hatched bars represent the putative new functions of PIMREG. (**K, L)** PIMREG co-localization with YY1 and PRMT5 in the nuclei of TMZ-treated GBM cells. (K) Representative confocal images of T98G cells treated with TMZ. PIMREG is shown in green (Alexa Fluor 488), YY1 in red (Alexa Fluor 594) and nucleus (DAPI) in cyan. The right panel shows line-scan intensity profiles across the nucleus of the cell indicated (bottom right), illustrating the spatial distribution of PIMREG and YY1 signals. The overlap in signal peaks is supported by a strong positive correlation between PIMREG and YY1 (Pearson’s *r* = 0.621). (L) Representative confocal images of U87MG cells treated with TMZ. Fluorescence signals were pseudo-colored in ImageJ: the Alexa Fluor 555 channel (PIMREG) is displayed in green, while the Alexa Fluor 488 channel (PRMT5) is displayed in red. Nuclei (DAPI) are shown in cyan. The line-scan intensity profiles across the nucleus (right panel) demonstrates a strong overlap between PIMREG and PRMT5 signals, with high co-localization coefficient (Pearson’s *r* = 0.914). Scale bar: 10 μm.

GO analysis of the proteins identified as the TMZ-regulated interactors of PIMREG in U87MG retrieved 10 ontological annotations whereas the proteins identified in PIMREG interactome and TMZ-regulated interactome of T98G, retrieved 16 and 50, respectively. These ontological annotations were classified according to their related biological processes (BP) (Fig. 2H, 2I and 2J; Supplementary Tables 8, 9 and 10). The BP terms obtained for TMZ-regulated PIMREG interacting proteins in U87MG cells were related to RNA splicing/processing, cell division/proliferation and mitochondrial metabolism (Fig. 2H; Supplementary Table 8). The BP terms obtained for PIMREG interactome in T98G cells were related to insulin pathway, transcription/mRNA processing, immune response, protein transport, and cell signaling (Fig. 2I; Supplementary Table 9). Lastly, the BP terms described for TMZ-regulated PIMREG interacting proteins identified in the T98G cells were related to cell differentiation/development, transcription/translation, macromolecule metabolic process, telomere maintenance, IFN signaling, ribosome biogenesis, apoptosis, DNA replication and cell division. Notably, the BP related to DDR exhibited the highest number of ontological annotations, which included telomere maintenance– (GO:1904507), intrinsic apoptotic– (GO:0008630), mitotic check-point (GO:0044773) signaling pathways in responses to DNA damage as well as regulation of DNA repair (GO:0006282) and DSB repair via homologous recombination (GO:0000724), underscoring the role of PIMREG in the DDR of T98G cells treated with TMZ (Fig. 2J; Supplementary Table 10).

Co-localization analyses were performed to corroborate the TMZ-regulated interaction of PIMREG with two identified DNA repair factors involved in HR-mediated resolution of DSBs, namely YY1 in T98G cells and PRMT5 in U87MG cells. The analysis revealed strong spatial overlap between PIMREG and both YY1 and PRMT5. Consistently, line-scan intensity profiles across the nuclei demonstrated marked overlap of PIMREG with YY1 and PRMT5 signals, supported by high co-localization coefficients (Pearson’s *r* = 0.621 for YY1 and *r* = 0.914 for PRMT5; Fig. 2K and 2L).

### 3.3. PIMREG is a substrate of ATM/ATR in GBM cells treated with TMZ

PIMREG is a phospho-protein [43] containing residues, which lie within the S/TQ motif, a known target site for ATM and ATR phosphorylation [44]. To ask if PIMREG is a substrate of ATR/ATM, we precipitated endogenous PIMREG from TMZ– and DMSO-treated as well as untreated T98G GBM cells and blotted the precipitated protein against the phospho-ATM/ATR substrate antibody (Fig. 3A) used to detect proteins containing the phosphorylated substrate motif of ATM/ATR. Under all conditions we identified the precipitated PIMREG protein (Fig. 3A, lower panel, lanes 3, 6 and 9). In the reciprocal immunoprecipitation experiment, precipitated phospho-ATM/ATR substrates (Fig. 3B, upper panel) were blotted against anti-PIMREG antibody (Fig. 3B, lower panel) and the PIMREG protein could be detected by anti-PIMREG among the precipitated phospho-ATM/ATR substrates in TMZ-treated cells (Fig. 3B, lower panel, lane 9). These results strongly suggest that PIMREG is a direct substrate for ATM/ATR kinases *in vivo*.

**Fig. 3:**
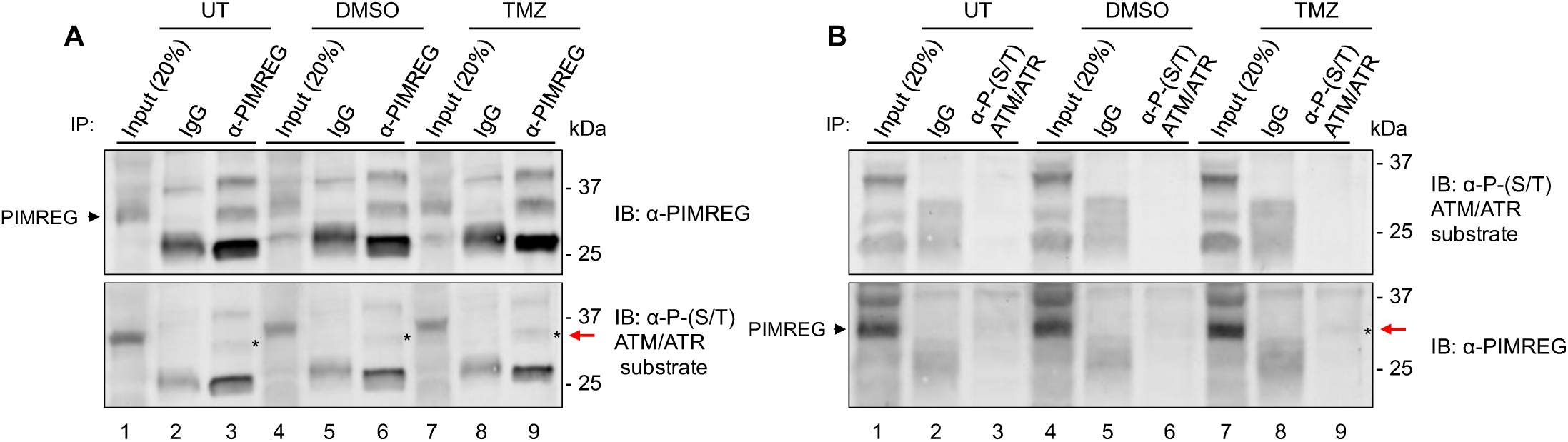
PIMREG is a substrate of ATM/ATR. Lysates of T98G cells untreated (UT) or treated with TMZ and vehicle (DMSO) for 24 h were precipitated with (**A**) anti-PIMREG antibody and isotype control (IgG). Western blot analysis with anti-PIMREG antibody shows the precipitated protein in the upper panel. Precipitated PIMREG is indicated with the arrowhead (lanes 3, 6 and 9). Immunoblotting of the precipitated protein with anti-phospho-ATM/ATR substrate antibody is shown in the lower panel. Asterisks (*) indicate the precipitated PIMREG protein detected with the anti-phospho-ATM/ATR substrate antibody. (**B**) ATM/ATR substrates immunoprecipitated using anti-phospho-ATM/ATR substrate antibody were immunoblotted with the same antibody (upper panel) and anti-PIMREG antibody (lower panel). Asterisk (*) indicates the precipitated phospho-ATM/ATR substrates detected with anti-PIMREG antibody (lane 9). As input control, lanes 1, 4 and 7 were loaded with 20 % of the protein extract used for each precipitation reaction. UT: untreated.

### 3.4. PIMREG deletion interferes with γH2AX and RAD51 accumulation in GBM cells

To investigate the role of PIMREG in the cellular response to DNA damage, we generated PIMREG knockout GBM T98G cell lines (PIMREG KO-1, KO-2, KO-3 and KO-4) and measured γH2AX levels by Western blot to monitor DNA damage and signaling. Quantification of γH2AX band intensity revealed that PIMREG KOs exhibited higher levels of γH2AX than the wild-type (WT) cells (Fig. 4A and 4B; Supplementary Fig. 2A), suggestive of higher basal DNA damage levels and/or genomic instability in the absence of PIMREG when compared to controls.

**Fig. 4:**
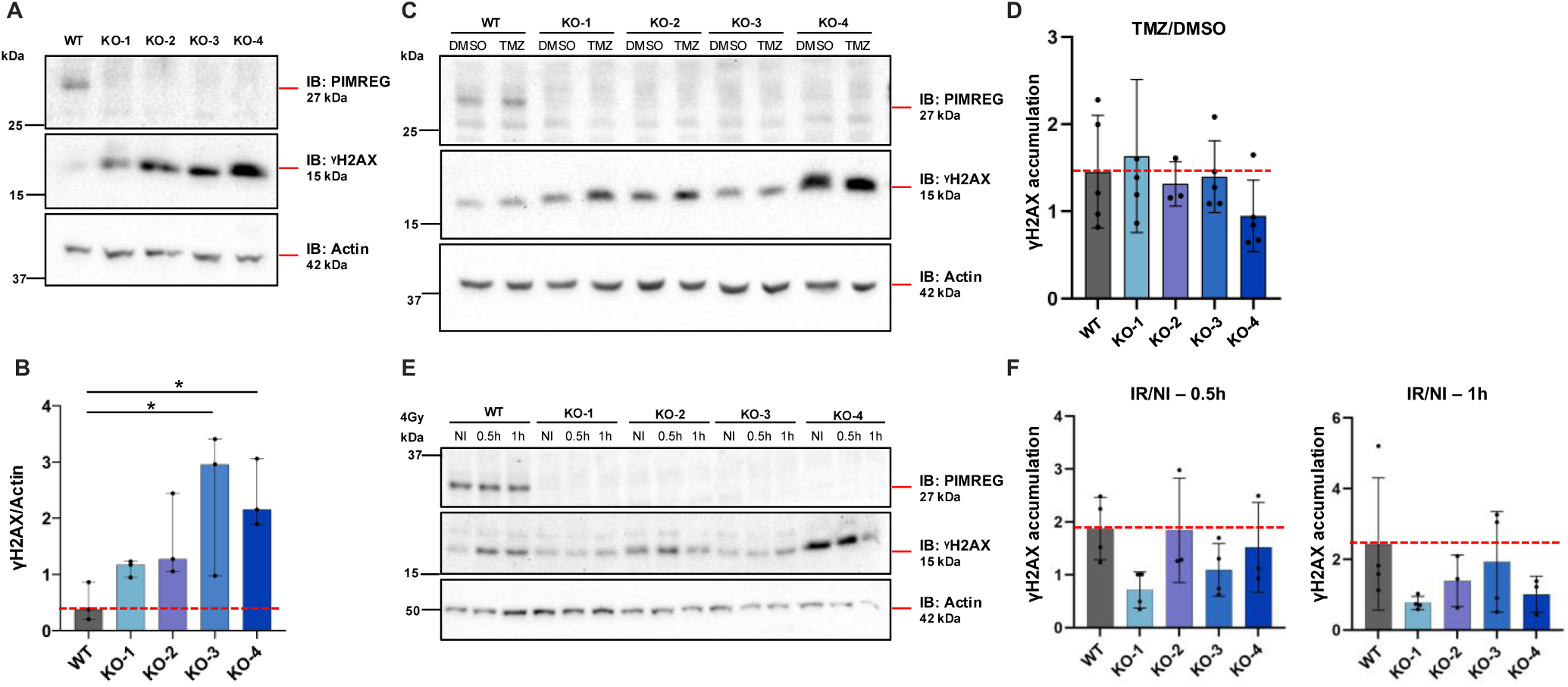
γH2AX accumulation in PIMREG KO cells|. (**A**) Western blot analysis of protein extracts from T98G WT and PIMREG KO (KO-1, KO-2, KO-3 and KO-4) cells. Membranes were blotted with anti-PIMREG, anti-phospho-H2AX (S139) antibodies. Actin was used as a loading control. (**B**) Bar graph showing densitometric quantification of YH2AX bands, normalized to their corresponding endogenous actin levels (YH2AX/actin). Results are shown as mean ± SD (standard deviation) of three independent experiments. Red dashed line indicates γH2AX levels in WT cells for comparison. (**C**) Western blot analysis of protein extracts derived from T98G WT and PIMREG KOs treated with TMZ or vehicle (DMSO) for 24 h. (**D**) Bar graph showing γH2AX accumulation expressed as the ratio of γH2AX levels (YH2AX/actin) in TMZ-treated cells relative to their corresponding DMSO control (TMZ/DMSO). Results are shown as mean ± SD (standard deviation) of five independent experiments. Red dashed line indicates the γH2AX accumulation in WT cells upon TMZ treatment. (**E**) Western blot analysis of protein extracts from T98G WT and PIMREG KO cells following IR exposure. Samples were collected at 0.5 and 1 h post-irradiation. Tubulin was used as a loading control. (**F**) Bar graph showing γH2AX accumulation expressed as the ratio of γH2AX levels (YH2AX/tubulin) in IR-treated cells relative to their corresponding non-irradiated controls (IR/NI). Results are shown as mean ± SD (standard deviation) of four independent experiments. Red dashed line indicates γH2AX accumulation in WT cells following IR exposure. Representative western blots are shown. (*) *p*-value < 0.05, One-way ANOVA test.

We next examined the relative levels of γH2AX in the PIMREG KOs treated with TMZ and IR to control cells. As expected, γH2AX signal increased in WT cells treated with both TMZ and IR compared to DMSO or non-irradiated (NI) control cells (Fig. 4C and 4E, middle panels).

Although PIMREG KOs exhibit higher levels of basal γH2AX than WT cells, when the amount of γH2AX of TMZ– or IR-treated cells was normalized by those of DMSO or NI (TMZ/DMSO or IR/NI), comparable levels of γH2AX accumulation were observed in PIMREG KOs and WT treated with TMZ, with the exception of PIMREG KO-4 which exhibited reduced levels of γH2AX accumulation (Fig. 4D; Supplementary Fig. 2B).

In PIMREG KOs cells treated with IR, reduced levels of γH2AX accumulation were observed across all clones (Fig. 4F; Supplementary Fig. 2C). Taken together, these results suggest that although PIMREG KO cells exhibit high levels of basal γH2AX, its absence compromises the accumulation of γH2AX induced following exposure to genotoxic stress.

To complement these results, immunofluorescence experiments were performed to examine the fluorescence intensity of γH2AX in the nuclei of PIMREG KO and WT GBM cells treated with TMZ or exposed to IR. As shown in Figure 5, TMZ and IR treatments enhanced γH2AX signal in both PIMREG KOs and WT cells, compared to DMSO or non-irradiated (NI) control cells. However, the overall amount of γH2AX fluorescence intensity appeared significantly lower in the PIMREG KOs compared to WT cells (Fig. 5), with the exception of PIMREG KO-3 exposed to IR which exhibited similar levels of γH2AX accumulation compared to WT. Interestingly, in one PIMREG KO cell line (KO-2), where PIMREG protein expression was unexpectedly restored (KO-2-recPIMREG) (Fig. 5C; Supplementary Fig. S3), IR exposure led to the opposite effect, with markedly increased γH2AX accumulation (Fig. 5D). At present we do not know the underlying mechanism of PIMREG recovery. Nevertheless, the data suggest that the presence of PIMREG could rescue the impaired DNA damage response phenotype observed in its absence, which is supported by the fact that the same KO-2 cell line, prior to recovery of PIMREG (Fig. 5A), exhibited significant reduction of γH2AX following TMZ treatment (Fig. 5B).

**Fig. 5.:**
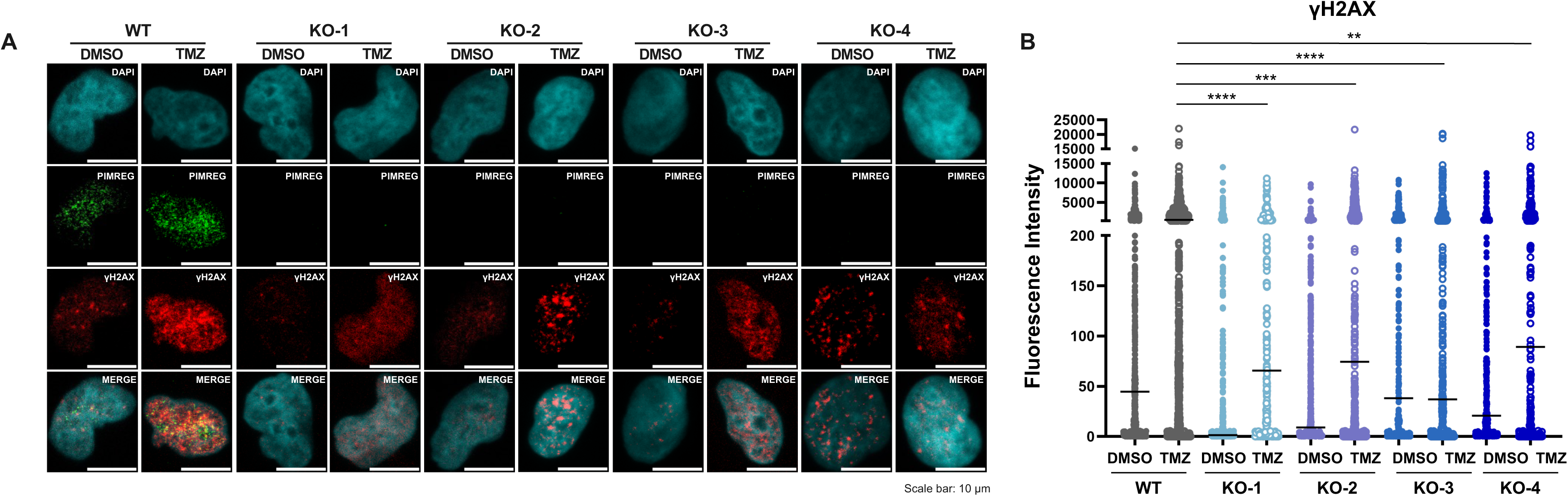
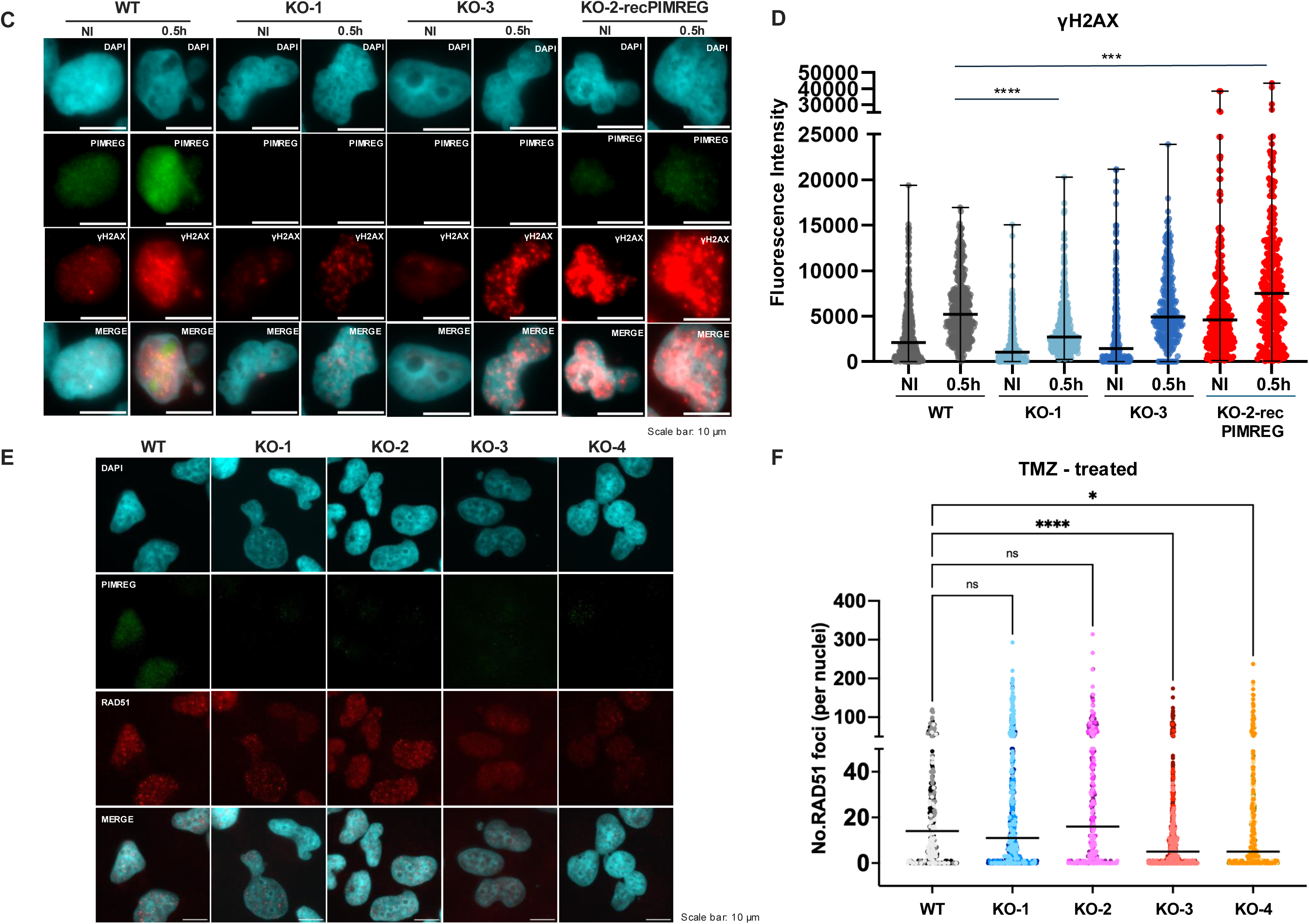
PIMREG deficiency impairs nuclear accumulation of γH2AX and RAD51 foci in GBM cells treated with genotoxic agents. Representative immunofluorescence analysis of T98G WT and PIMREG KO cells treated with (A) TMZ or vehicle (DMSO) for 24 h and (C) exposed to IR (4 Gy) and collected at 0.5 h post-irradiation. PIMREG is shown in green (Alexa Fluor 488), γH2AX in red (Alexa Fluor 555) and nuclei (DAPI) in cyan. Scale bar: 10 µM. (B, D) Fluorescence intensity quantification of nuclear γH2AX, quantified in at least 75 cells per condition. Data are from 3 to 5 independent experiments. The horizontal bars indicate the median intensity. (E) Representative images of T98G WT and PIMREG KO cells treated with TMZ and stainned for RAD51. PIMREG is shown in green (Alexa Fluor 488), RAD51 in red (Alexa Fluor 594) and nuclei (DAPI) in cyan. Scale bar: 10 µM. (F) Number of RAD51 foci was quantified in at least 60 cells per condition. Data are from 3 independent experiments, with with two technical duplicates performed for each KO unless otherwise stated. All replicates are shown. The horizontal bars indicate the median intensity. (*) *p*-value < 0.05, (**) *p*-value < 0.01, (***) *p*-value < 0.001 and (****) *p*-value < 0.0001, One-way ANOVA test followed by the post-test Kruskal-Wallis was applied. WT: wild-type; KO: knockout; KO-2-recPIMREG: KO-2 cells that recovered PIMREG’s expression; NI: non-irradiated.

Next, we evaluate the impact of PIMREG loss upon the formation of RAD51 foci at sites of DSBs in PIMREG KO and WT GBM cells exposed to TMZ. Immunofluorescence analysis revealed RAD51 foci to be readily detectable in WT GBM cells (median of 8 foci per nuclei, data not shown). This number increased in TMZ treated WT GBM cells (median of 14 foci per nuclei) supporting an increase in DSB formation after TMZ exposure. In contrast, deletion of PIMREG in GBM cells lead to a significant reduction in the number of RAD51 foci per nuclei in at least two independent KO cell lines (PIMREG KO-3 and PIMREG KO-4) when compared to WT cells. A similar trend, though not significant, was observed for PIMREG KO-1 cells (Fig. 5E and 5F).

Taken together, our results demonstrate that PIMREG plays a role in DDR signaling, with PIMREG deletion reducing both γH2AX levels and RAD51 foci formation in GBM cells under conditions of genotoxic stress.

### 3.5. PIMREG deletion compromises DNA damage signaling

We next asked whether PIMREG KO affects the activation of additional DNA damage response signaling molecules following treatment with to TMZ or IR (Fig. 6; Supplementary Fig. S4). Besides ATM, we analyzed molecules representing different steps of the DNA damage signaling cascade. These include KAP1, which facilitate DSB repair in both euchromatin and heterochromatin [45]; CHK1, a critical cell-cycle checkpoint regulator; and DNA-PKc, a central component of DSB signaling pathway that orchestrates NHEJ-mediated repair [46]

**Fig. 6:**
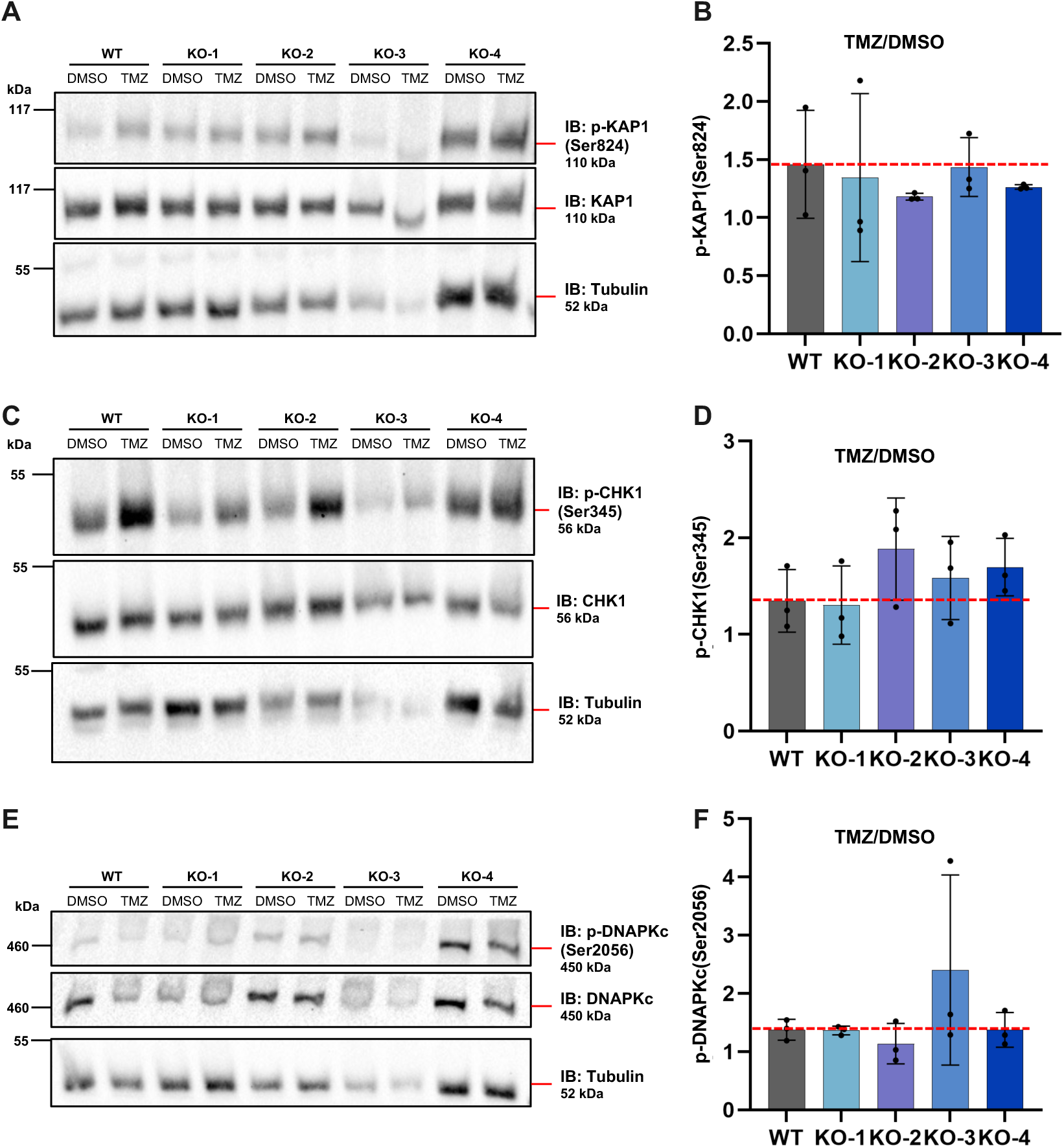
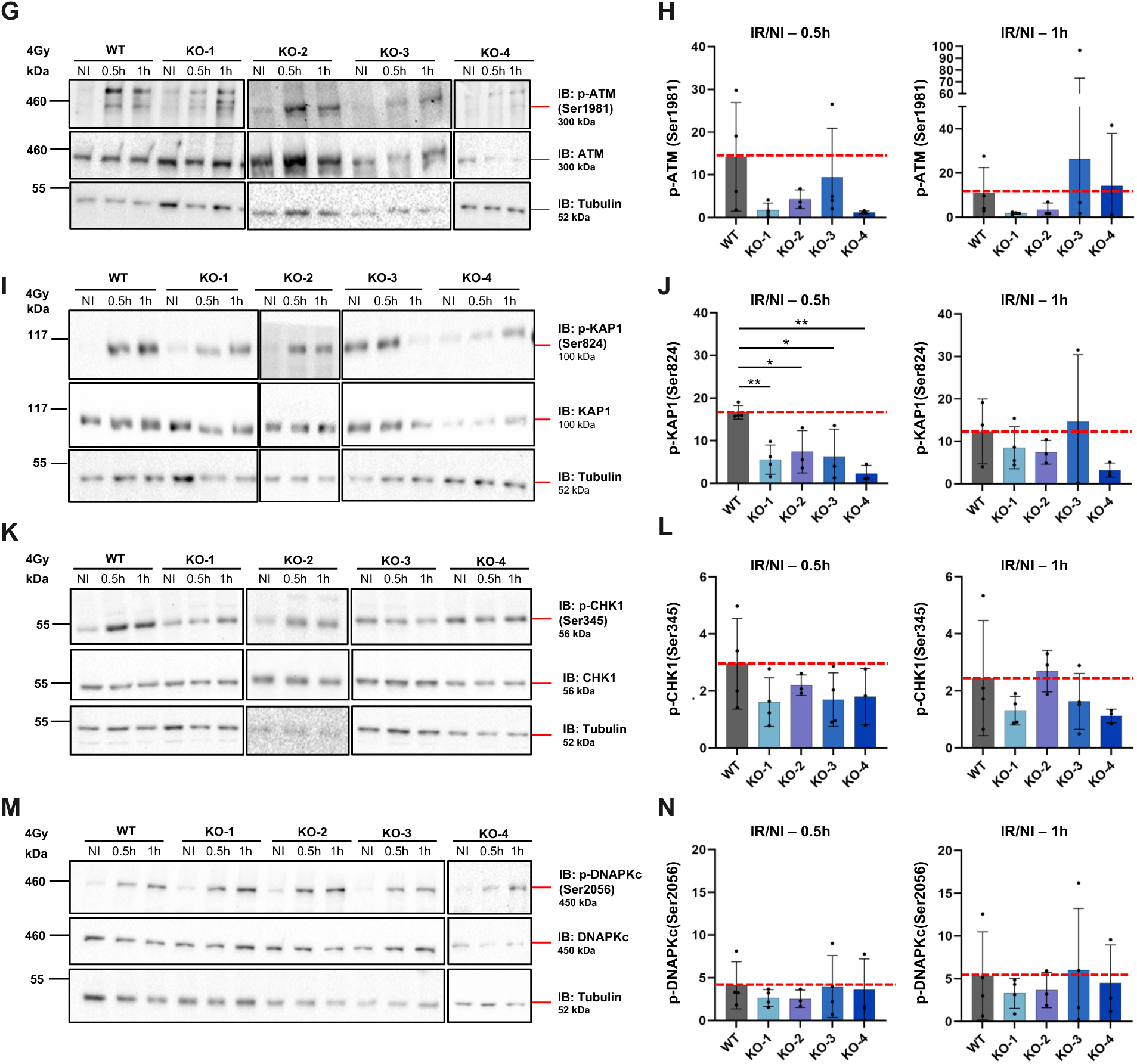
PIMREG KO affects activation of DNA damage response signaling molecule following IR exposure. (**A, C, E**) Western blot analysis of protein extracts derived from T98G WT and PIMREG KOs treated with TMZ or vehicle (DMSO) for 24 h. Membranes were blotted with anti-Phospho-KAP1 (Ser824) and KAP1 (A), anti-Phospho CHK1 (Ser345) and CHK1 (C) and anti-Phospho-DNA-PKc (Ser2056) and DNA-PKc (E) antibodies. Tubulin was used as a loading control. (**B, D, F**) Bar graph showing densitometric quantification of phosphorylated protein band intensities normalized to total protein levels. Protein activation is expressed as the ratio of the phosphorylation levels in TMZ-treated cells relative to their corresponding DMSO control (TMZ/DMSO). Results are shown as mean ± SD (standard deviation) of three independent experiment. Red dashed line indicates the protein activation in WT cells upon TMZ treatment. (**G, I, K, M**) Western blot analysis of protein extracts from T98G WT and PIMREG KO cells (KO-1, KO-2, KO-3, and KO-4) following exposure to IR (4Gy). Samples were collected at 0.5 and 1h post-irradiation. Membranes were blotted with anti-Phospho ATM (Ser1981) and ATM (G), anti-Phospho-KAP1 (Ser824) and KAP1 (I), anti-Phospho CHK1 (Ser345) and CHK1 (K), anti-Phospho-DNA-PKc (Ser2056) and DNA-PKc (M) antibodies. Tubulin was used as a loading control. (**H, J, L, N**) Bar graph showing densitometric quantification of phosphorylated protein band intensities normalized to total protein levels. Protein activation is expressed as the ratio of the phosphorylation levels in IR samples relative to their corresponding non-irradiated controls (IR/NI). Results are shown as mean ± SD (standard deviation) of four independent experiment. Red dashed line indicates the protein activation in WT cells at the indicated time points following IR exposure. (*) *p*-value < 0.05 and (**) *p*-value < 0.01, One-way ANOVA test. WT: wild-type; KO: knockout; NI: non-irradiated.

Western blot analysis and quantification of phosphorylated protein band intensities revealed no differences in the activation of KAP1, CHK1 or DNA-PKc between PIMREG KOs and WT cells after exposure to TMZ (Fig. 6A-6F). In contrast, PIMREG KO cells exposed to IR showed a consistent trend of reduced ATM, KAP1 and CHK1 activation compared to WT cells. This reduction was most evident at 0.5 h post-irradiation (Fig. 6G-6L). Activation of DNA-PKc was unaffected in PIMREG KOs cells relative to WT following IR exposure (Fig. 6M and 6N). Thus, loss of PIMREG has a broader impact on DDR signaling in irradiated GBM cells.

### 3.6. PIMREG deletion attenuates TMZ-induced DNA break accumulation

Alkaline comet assays were performed to assess whether the reduced damage signaling observed in PIMREG KO cells reflected a decrease in DNA break formation following TMZ treatment. The results showed that TMZ-treated PIMREG KO cells accumulated significantly fewer DNA breaks compared with WT cells. This reduction of DNA lesions was consistently observed in the KO-2 and KO-3 cell lines (Fig. 7).

**Fig. 7:**
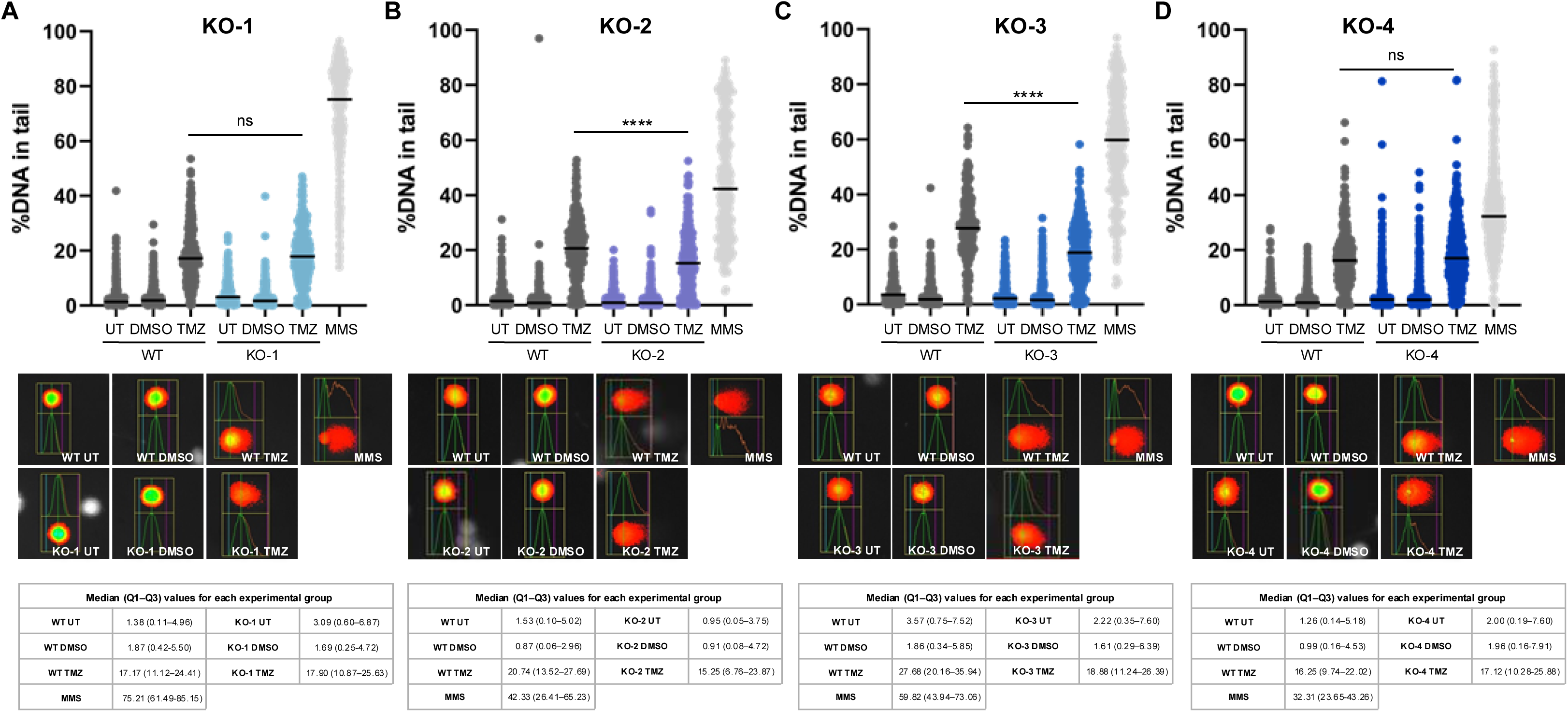
PIMREG deficient cells exhibit reduced DNA breaks after TMZ treatment. (A-D) Alkaline comet assay of T98G WT and PIMREG KO cells treated with TMZ or vehicle (DMSO). DNA damage was quantified as the percentage (%) of DNA in the comet tail relative to the head. One hundred cells were analyzed per condition in three (KO-1, KO-2 and KO-3) and four (KO-4) independent experiments. All replicates are shown. The horizontal bars indicate the median values. Representative images of nuclei with comet tails are shown below each graph. Tables report the median values with interquartile range (Q1-Q3) for each group. (****) *p*-value < 0.0001, unpaired Mann-Whitney test. WT: wild-type; KO: knockout; UT: untreated; MMS: methyl methanesulfonate (positive control).

### 3.7. PIMREG deletion interferes with the repair of DSBs

ATM, γH2AX and RAD51 are key regulators of DSBs signaling cascade to promote repair [46,47]. Given that PIMREG loss affects the signaling of these molecules, we sought to investigate its consequence on DSB repair.

To this end, we performed DNA repair assays using HR (U2OS DR-GFP) and NHEJ (HeLa EJ5-GFP) reporter cell lines [40,41] in which PIMREG was deleted. In the DR-GFP and EJ5-GFP reporter systems, the expression of GFP protein is restored upon HR and NHEJ repair, respectively.

The absence of PIMREG significantly impaired DNA repair by HR in the U2OS DR-GFP PIMREG KOs (KO-1, KO-2, KO-3 and KO-4). Although the reduction of the HR-mediated repair was not statistically significant in the PIMREG KO-2 cell line, this trend was consistently observed across the three independent experiments (Fig. 8A). Additionally, the absence of PIMREG was associated with an increase in DNA repair by the NHEJ pathway in one out of four HeLa EJ5-GFP PIMREG KOs (KO-4), with a similar trend being observed in the PIMREG KO-3 cell line (Fig. 8B).

**Fig. 8:**
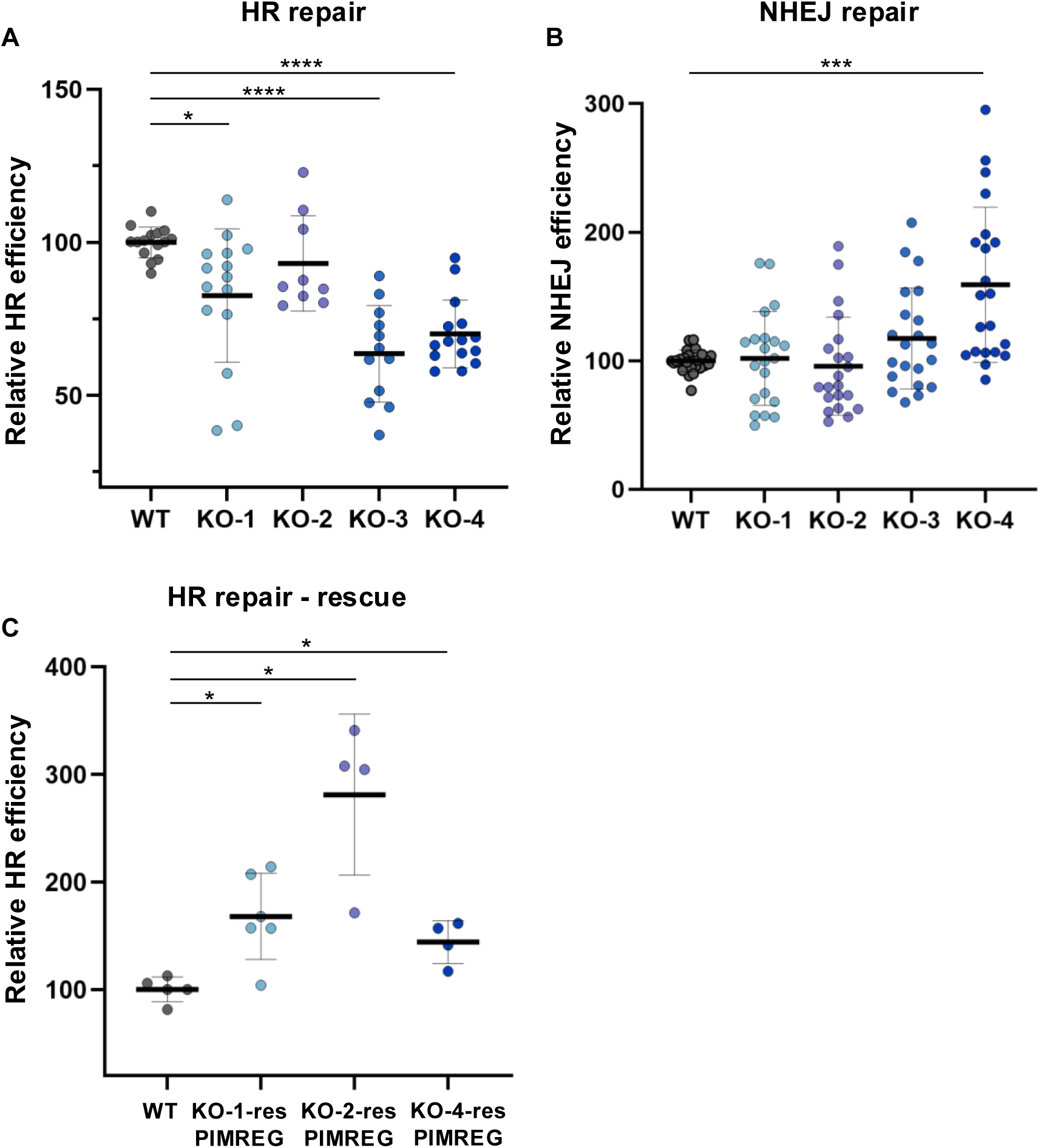
DNA repair reporter assay. The graphs show the percentage of PIMREG WT, PIMREG KO (KO-1, KO-2, KO-3, and KO-4) and rescued (KO-1-resPIMREG, KO-2-resPIMREG, and KO-4-resPIMREG) cells that repaired the DSBs. GFP reports the repair events via (A, C) HR in U2OS DR-GFP and (B) NHEJ in HeLa EJ5-GFP cells. (C) Rescue experiment in HR reporter cells. U2OS DR-GFP – PIMREG-KO-1, –KO2 and – KO-3 cells ectopically expressing PIMREG (KO-1-resPIMREG, KO-2-resPIMREG, and KO-4-resPIMREG) or control vector (WT), all carring the tdTOMATO reporter. GFP-positive cells were specifically analyzed within the tdTOMATO-positive population. The percentage of GFP-positive cells was normalized to the mean percentage of WT control cells within each experiment. Experiments were performed at least two times in triplicates. Fifty thousand events were acquired. Sample sizes for HR repair (A): WT (n = 5), KO-1 (n = 5), KO-2 (n = 3), KO-3 (n = 4), KO-4 (n = 5). Sample sizes for NHEJ repair (B): WT (n = 9), KO-1, KO-2, KO-3 and KO-4 (n = 7). Sample sizes for HR repair – rescue (C): WT (n = 3), KO-1-resPIMREG (n = 3), KO-2-resPIMREG (n = 2), KO-4-resPIMREG (n = 2). (*) *p*-value < 0.05, (***) *p*-value < 0.001 and (****) *p*-value < 0.0001, Brown-Forsythe and Welch ANOVA.

Rescue experiments demonstrated that ectopic expression of PIMREG markedly increased HR repair efficiency in the PIMREG KO cell lines by 67 % (KO-1-resPIMREG), 180 % (KO-2-resPIMREG) and 44 % (KO-4-resPIMREG) relative to WT cells (Fig. 8C), fully restoring the HR repair capacity impaired under KO conditions (Fig. 8A). These findings confirm that the observed phenotype is specifically attributable to PIMREG loss.

Together, these findings suggest that PIMREG plays a role in promoting HR-mediated DSBs repair, while its absence may favor repair by NHEJ pathway.

## 4. Discussion

This study greatly expands our understanding of PIMREG, redefining it from a proliferation marker to an integrated component of the DDR in GBM. Our findings reveal that PIMREG functions as a direct substrate for ATM/ATR kinases, interacts with key DNA repair proteins, and actively facilitates the cellular response to chemotherapy– and radiotherapy-induced DNA damage. By demonstrating its role in signaling and repairing of DSBs, we consolidate PIMREG as a key player in maintaining genomic integrity and possibly, promoting treatment resistance.

Pathway enrichment analysis performed using GBM patient samples revealed putative functions for PIMREG across multiple DNA repair pathways and in RNA splicing and processing, finding supported by our mass spectrometry data. Specifically, our PIMREG interactome uncovered novel associations with key DNA repair factors involved in HR-mediated resolution of DSB, including YY1 in T98G cells, and TRIP13 and PRMT5 in U87MG.

YY1 is a transcription factor, which complex with chromatin-remodeling factors and is essential for HR-based DNA repair [48]. TRIP13 is an ATPase involved in early DNA damage sensing and ATM signal amplification [49] thereby contributing to the promotion of HR [50]. PRMT5, an arginine methyltransferase with described roles in splicing and stemness in GBM [51] is also a key regulator of HR-mediated DSB repair [52]. Notably, such associations between PIMREG and DDR-related proteins were only observed in TMZ-regulated interactomes, suggesting that these interactions or the modulation of binding of these proteins to PIMREG are specific to TMZ exposure, and thus DNA damage, further evidencing roles for PIMREG in DSB repair by HR.

In both TMZ-regulated PIMREG interactome and PIMREG interactome under control conditions, we identified putative interactions between PIMREG and proteins related to RNA splicing and processing. To the best of our knowledge, this is the first report linking PIMREG to RNA metabolism. Such observations provide a strong foundation for future investigation into PIMREG’s role in RNA metabolism in GBM. Moreover, the difference in the splicing network observed between PIMREG interactome under control condition or induced by TMZ treatment evidences a dynamic role in RNA processing that may be influenced in response to the genotoxic stress. It is tempting to speculate that this shift in the splicing network relates to a non-canonical function of PIMREG in DDR-specific splicing. Mounting evidence supports crosstalk between RNA splicing and the DDR [53–58], for example, the splicing factor SF3B2, a putative interactor of PIMREG identified in this study, has been previously linked to HR-mediated repair of DSB [59]. In support, we have recently reported UHMK1, a known PIMREG interacting protein [43], as a splicing regulatory kinase [60]. In addition to phosphorylating PIMREG and several mRNA related proteins, UHMK1 also regulates the phosphorylation of critical DDR proteins, including RAD54L [61], THRAP3, BCLAF1 [62], and TPT1 [63].

Another PIMREG-associated function observed from TMZ-regulated PIMREG interactome was related to NADH dehydrogenase activity and mitochondrial metabolism, suggestive of a potential role for PIMREG in mitochondrial regulation under genotoxic stress. Though metabolic reprogramming is a known mechanism of TMZ resistance in GBM [64], how PIMREG contributes in this context remains to be investigated.

We observed distinct PIMREG protein interacting profiles between the two GBM cell lines. Though both cell lines support TMZ-induced PIMREG interactions with DDR-related proteins, interactions related to RNA splicing and mitochondrial metabolism appeared exclusive to U87MG cells. Such differences likely reflect the distinct genetic background of the two cell lines: T98G cells are MGMT-proficient and carry *TP53* mutation, while U87MG cells are MGMT-deficient and *TP53* wild-type, features that enable distinct response to TMZ treatment [65].

Surprisingly, several proteins previously reported to interact with PIMREG were not identified in our dataset, including PICALM [66], UHMK1 [43], PrPC [67], ROD1 [68], NuRD complex [14], STAT3 [17], NFkB/p65 [20], and BRSK2 [25]. A plausible explanation is that these interactions were originally identified in different cell lines.

In summary, our proteomic findings expand the current understanding of PIMREG’s interaction network in established and novel cellular processes under both physiological and genotoxic stress contexts of GBM cells.

PIMREG has two amino acid residues, serine at position 137 (S137) and threonine at position 114 (T114) that could be sites for ATM/ATR phosphorylation [44]. Our data shows that PIMREG is recognized by the phospho-ATM/ATR substrate antibody, suggestive of PIMREG as a direct substrate of these DDR kinases. This is further supported by the identification of S137 and T114 of PIMREG as phosphosites in proteomic studies enriched for phospho-ATM/ATR substrates and curated into PhosphoSitePlus® database (http://www.phosphosite.org). Notably, phosphorylated PIMREG was detected under DMSO– and TMZ-treated, as well as in untreated conditions. Therefore, whether ATM/ATR-mediated phosphorylation of PIMREG is specifically induced in response to DNA damage or occurs as part of a broader mechanism remains to be determined.

PIMREG deficient cells in the absence of genotoxic stress exhibited elevated levels of γH2AX, suggesting PIMREG importance for genomic stability. Conversely, upon exposure to TMZ or IR, these cells overall exhibited an impaired DSB response. Quantification of γH2AX fluorescence intensity confirmed reduced γH2AX in both TMZ and IR treated KO cells. However, under IR treatment, in which three out of four PIMREG KO cell lines were assessed, reduced levels of γH2AX were observed exclusively in one PIMREG KO cell line.

Unexpectedly, the PIMREG KO-2 cell line restored PIMREG protein expression after extended culturing. Remarkably, upon genotoxic stress, PIMREG KO-2 that regained PIMREG (KO-2-recPIMREG) displayed increased levels of γH2AX, effectively reversing the phenotype observed in early passages and highlighting the heterogeneity among the KO cell lines. The mechanism behind PIMREG re-expression in the KO-2 cell line remains unclear. However, given that the T98G cells employed for genome editing, are highly polyploid cells [69], it is plausible that one or more *PIMREG* alleles would lie within chromatin regions that were inaccessible to Cas9 at the time of editing. Over time, selective pressures during cell culture may have led to chromatin remodeling, enabling reactivation of a previously silenced, unedited allele and restoration of PIMREG expression.

We observed reduced loading of RAD51 in some of our KO cells, these data were in agreement with reduced levels of γH2AX in TMZ-treated cells.

Reduced activation of ATM (Ser1981) in both PIMREG KO cells exposed to IR and in PIMREG knockdown cells treated with TMZ [35] also supports roles for PIMREG in the DSB signaling under damage. Given that γH2AX and RAD51 loading represents key early events in DSB signaling [46], PIMREG may also act during the early stages of the cell response to damage. Consistent with this, we demonstrated that KAP1 activation was significantly impaired in PIMREG KOs cells exposed to IR. KAP1 is a protein involved in chromatin condensation. Following DSB induced by IR, ATM phosphorylates KAP-1 at S824 at the site of damage, where it colocalizes with γH2AX foci and rapidly spreads throughout the chromatin [70]. Phosphorylated KAP1 facilitates DSB repair in both euchromatin and heterochromatin [45].

Noteworthy, the impaired damage signaling observed in PIMREG KO cells subjected to genotoxic stress reflects, at least in part, a reduced extent of DNA break formation demonstrated by comet assays.

Importantly, our data is consistent with a role for PIMREG during HR-direct DSB repair; loss of PIMREG significantly reduced the formation of RAD51 foci in T98G cells treated with TMZ in at least two KO cell lines and, in addition, a direct evaluation of HR activity coincided with reduced HR capacity in the absence of PIMREG and ectopic expression of PIMREG restored the HR capacity on rescue experiments. In contrast, NHEJ repair was only mildly enhanced in two out four KO cell lines.

In line with these observations, DNA-PKc activation, a central orchestrator of NHEJ, was unaffected by PIMREG absence after IR exposure, further supporting specific roles for PIMREG in HR. PIMREG’s influence on HR may be partially explained by its expression pattern throughout the cell cycle. PIMREG levels become elevated during S and G2, sharply declining as cells enter G1 [13]. HR is predominantly active in S and G2 phases, due to the availability of a sister chromatid for repair, thus this temporal expression profile may underlie PIMREG’s preferential role in supporting HR over NHEJ.

Overall, our findings support a role for PIMREG in facilitating DSB signaling and repair following damage, however it is important to note that a degree of phenotypic heterogeneity was observed across treatments (TMZ and IR) and amongst KO clones. The origin of this variability may be a result of off-target effects or the clonal differences arising during the monoclonal selection of the KOs.

We observed a distinct difference in the activation of the DNA signaling molecules in PIMREG KO cells treated with TMZ or exposed to IR. These differences are likely attributable to the nature and timing of DNA damage induced by each agent. IR rapidly induces DSBs whereas TMZ primarily causes base alkylation. The formation of secondary DSBs after TMZ treatment depends on one or more rounds of DNA replication with futile MMR repair [9]. It is conceivable that our cells, treated with TMZ for 24h, harbor significant lower levels of DSBs at the time point analyzed, perhaps explaining why western blot analysis failed to detect substantial changes in DSB signaling in TMZ-treated PIMREG KO cells. In contrast, by immunofluorescence analysis, we detected a reduction in γH2AX intensity in TMZ-treated KO cells. Notably, γH2AX fluorescence intensity was approximately 100-fold higher in IR-treated KO-1 cells compared to those treated with TMZ, highlighting the stark difference in DSB burden between the two treatments. These findings, together with the reduced number of RAD51 foci per nuclei in the TMZ treated cells, strongly implicate PIMREG as a DDR factor, specifically PIMREG may act to support HR-directed repair of DSBs in GBM cells. A key question emerging from this study is whether PIMREG participates in these pathways directly or exerts its effects indirectly through broader cellular processes.

Given such broad functionality for PIMREG, our data is more suggestive of PIMREG as an indirect contributor to the DDR, either through the regulation of upstream or parallel processes essential for an effective response to genotoxic stress. PIMREG’s multifaceted roles are therefore especially important to understand in the context of cancer. Especially given the consistent overexpression of PIMREG in many, if not all, types of cancer, suggesting dysregulation of PIMREG activity may promote tumor progression and treatment resistance, not only in GBM but in other cancer types.

In conclusion, our findings provide new evidence that PIMREG plays a role in DSB signaling and HR repair highlighting PIMREG as a potential contributor to therapy resistance and a promising target for future cancer therapies.

## Credit authorship contribution statement

**Maria Vitória de Rizzo Gasparini:** Investigation, Conceptualization, Visualization, Formal analysis, Writing – Original Draft. **Laís do Carmo:** Investigation, Visualization, Formal analysis, Writing – Original Draft. **Giovanna Gonçalves de Almeida:** Investigation, Visualization, Formal analysis, Writing – Original Draft. **Mariana Siqueira Lacerda Mamede:** Investigation. **Isabela David Cardoso:** Investigation. **Lígia Arantes Sardinha:** Investigation. **Isabela Spido Dias:** Investigation. **Jennifer Ann Black:** Supervision, Writing – Review & Editing. **Vanessa C Arfelli:** Validation, Supervision, Writing – Review & Editing. **Yun-Chien Chang:** Investigation, Formal analysis, Writing – Review & Editing. **Elza Tiemi Sakamoto Hojo:** Resources, Writing – Review & Editing**. Stephanie Heinzlmeir:** Supervision, Writing – Review & Editing. **Bernhard Kuster:** Resources, Writing – Review & Editing. **Philipp A. Greif:** Supervision, Resources, Writing – Review & Editing. **Leticia Fröhlich Archangelo:** Conceptualization, Supervision, Writing – Original Draft, Project administration, Funding acquisition.

## Data availability

The raw mass spectrometry data, MaxQuant search results, and the Swiss-Prot reference database used for analysis have been deposited to the ProteomeXchange Consortium via the MassIVE partner repository with the dataset identifier PXD069394 (MassIVE ID: MSV000099465).

## Supporting information

This article contains supporting information.

## Acknowledgements

We thank Dr. Jeremy A. Squire for the English review and Patrícia Vianna Bonini, Camila Menezes Bonaldo (FACS) and Leandro Federiche Borges (IR) for technical assistance. We are also grateful to Dr. Carlos Frederico Martins Menck, Katlin B. Massirer and Cesar Seigi Fuziwara for kindly providing aliquots of anti-RAD51, anti-PRMT5 and anti-YY1 antibodies, respectively. Marlon Fortes Rocha and Dr. Larissa Dias da Cunha are acknowledged for helping with the FACS analysis. PAG acknowledges support by the Wilhelm Sander-Stiftung (Förderantrag Nr. 2014.162.3). This research was supported by the Fundação de Amparo à Pesquisa do Estado de São Paulo (FAPESP 2019/26035-1 and 2025/02105-1 to LFA). FAPESP supports the LMMC (2004/08868-0).

