## Supplementary material for "Loss of PIMREG impairs double-strand break signaling and repair"

### Supplementary Figure S1

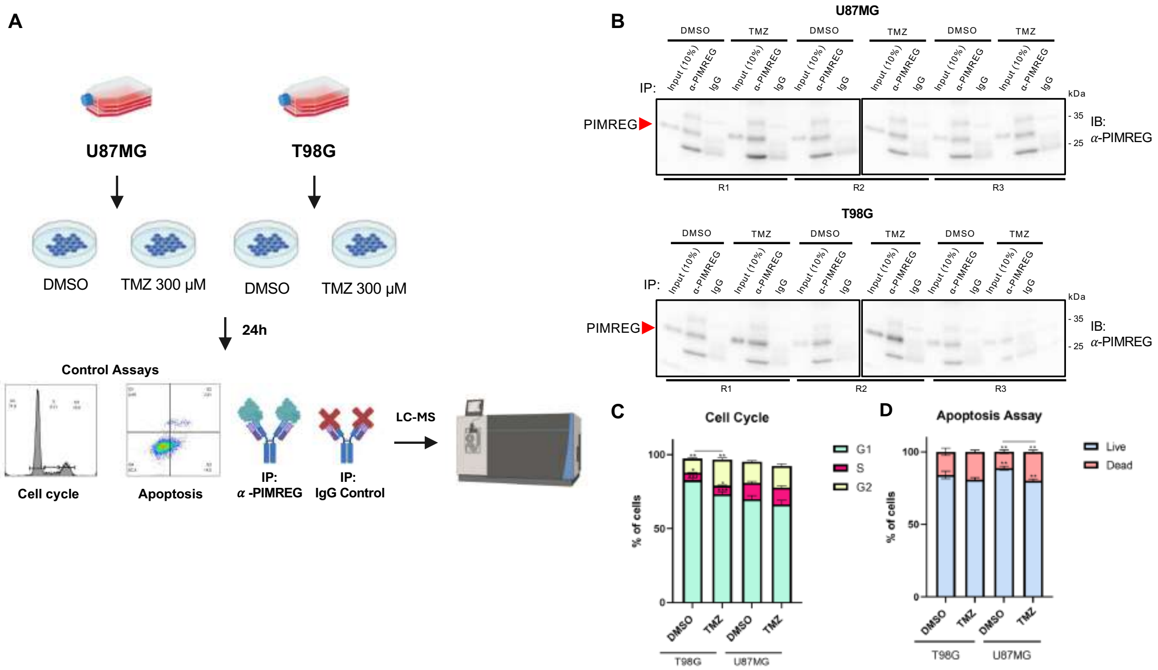


**Fig. S1: Experimental design and sample preparation for liquid chromatography-mass spectrometry (LC-MS) analysis.** (**A**) Experimental workflow. GBM cells (U87MG and T98G) were treated with 300 μM TMZ or vehicle (DMSO) for 24 h. Following treatment, cells were harvested for protein extraction, immunoprecipitation, and LC-MS analysis. Each sample was assessed for cell cycle and apoptosis. (**B**) Immunoblotting analysis of the samples submitted to LC-MS analysis. Precipitated PIMREG is shown in the samples incubated with anti-PIMREG antibody but not with the isotype control (IgG). A total of 10 % of the lysate was used as input. (**C**) Cell cycle and (**D**) apoptosis analysis of the same samples. T98G cells treated with TMZ exhibited slightly increased percentage of cells in the G2 phase of the cell cycle in comparison to DMSO. U87MG cells treated with TMZ exhibited slightly decreased percentage of viable cells in comparison to DMSO. *p*-value < 0.0001, Two-way ANOVA test.

**Supplementary Figure S2:**

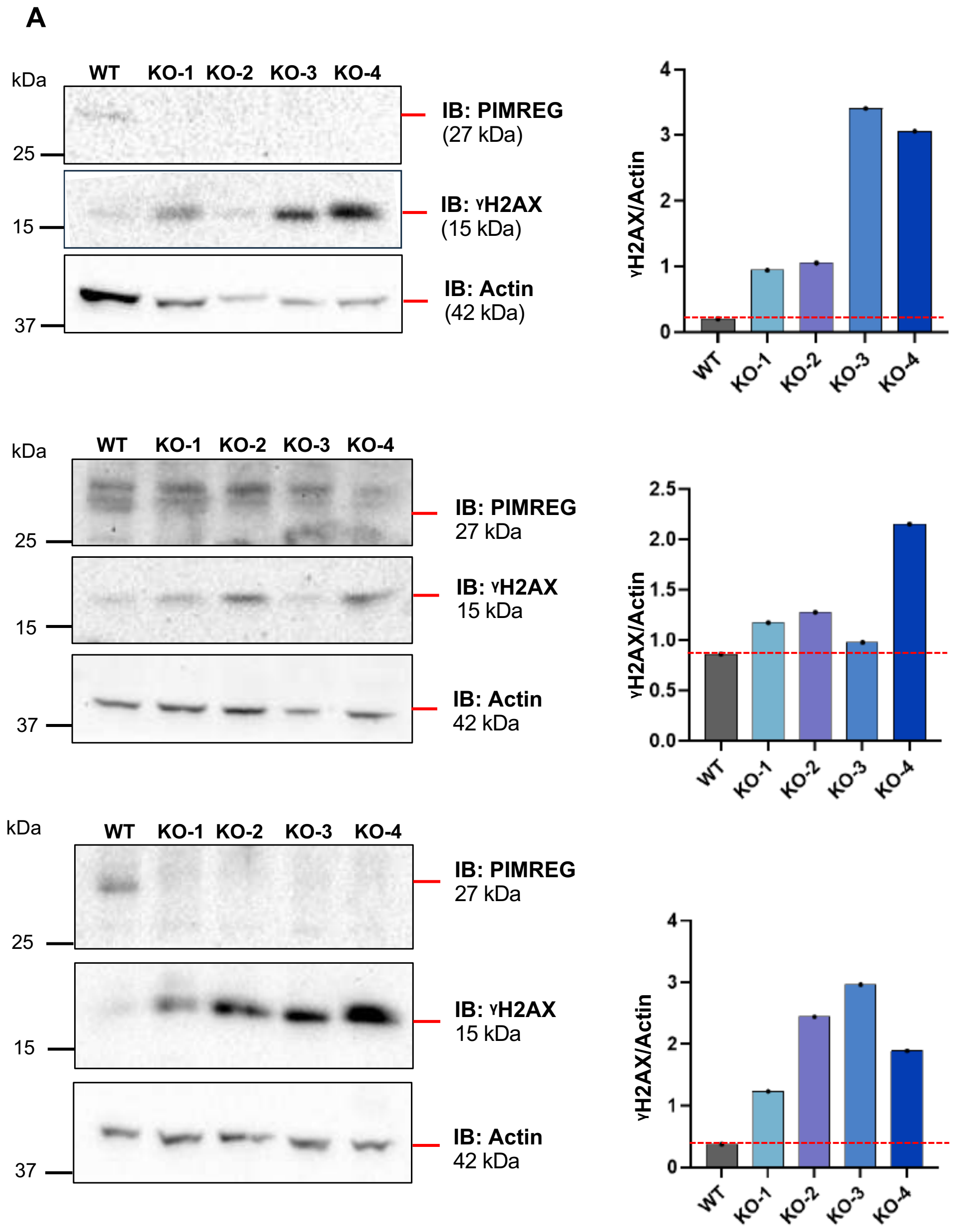


**Fig. S2A: Western blot analysis of γH2AX in T98G WT and PIMREG KO cells.** Membranes were blotted with anti-PIMREG, anti-phospho-H2AX (S139) antibodies. Actin was used as a loading control. On the right, bar graphs showing densitometric quantification of γH2AX bands, normalized to corresponding endogenous actin levels (γH2AX/actin) for each individual blot. Red dashed line indicates γH2AX levels in WT cells for comparison. Three independent experiments are shown.

**Supplementary Figure S2:**

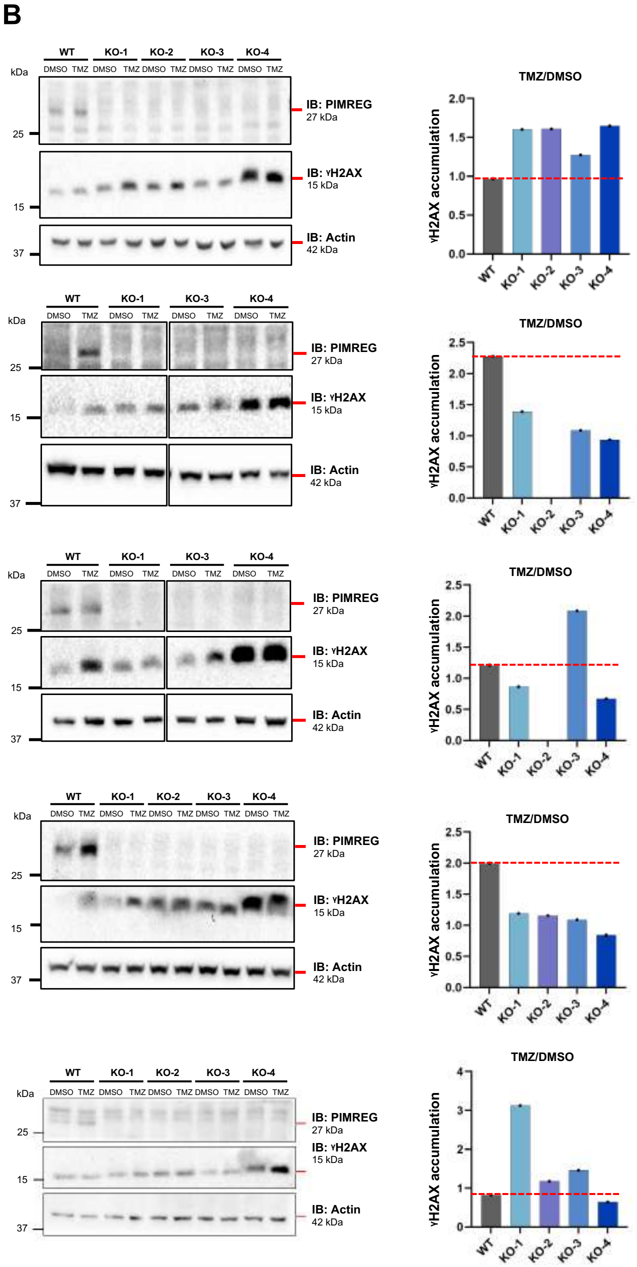


**Fig. S2B: Western blot analysis of γH2AX in T98G WT and PIMREG KO cells treated with TMZ.** Western blot analysis of protein extracts derived from T98G WT and PIMREG KOs treated with 300 µM TMZ or vehicle (DMSO) for 24 h. Membranes were blotted with anti-PIMREG, anti-phospho-H2AX (S139) antibodies. Actin was used as a loading control. On the right, bar graphs showing γH2AX accumulation expressed as the ratio of γH2AX levels in TMZ-treated cells relative to their corresponding DMSO control (TMZ/DMSO) for each individual blot. Red dashed line indicates the γH2AX accumulation in WT cells upon TMZ treatment. Five independent experiments are shown.

### Supplementary Figure S2:

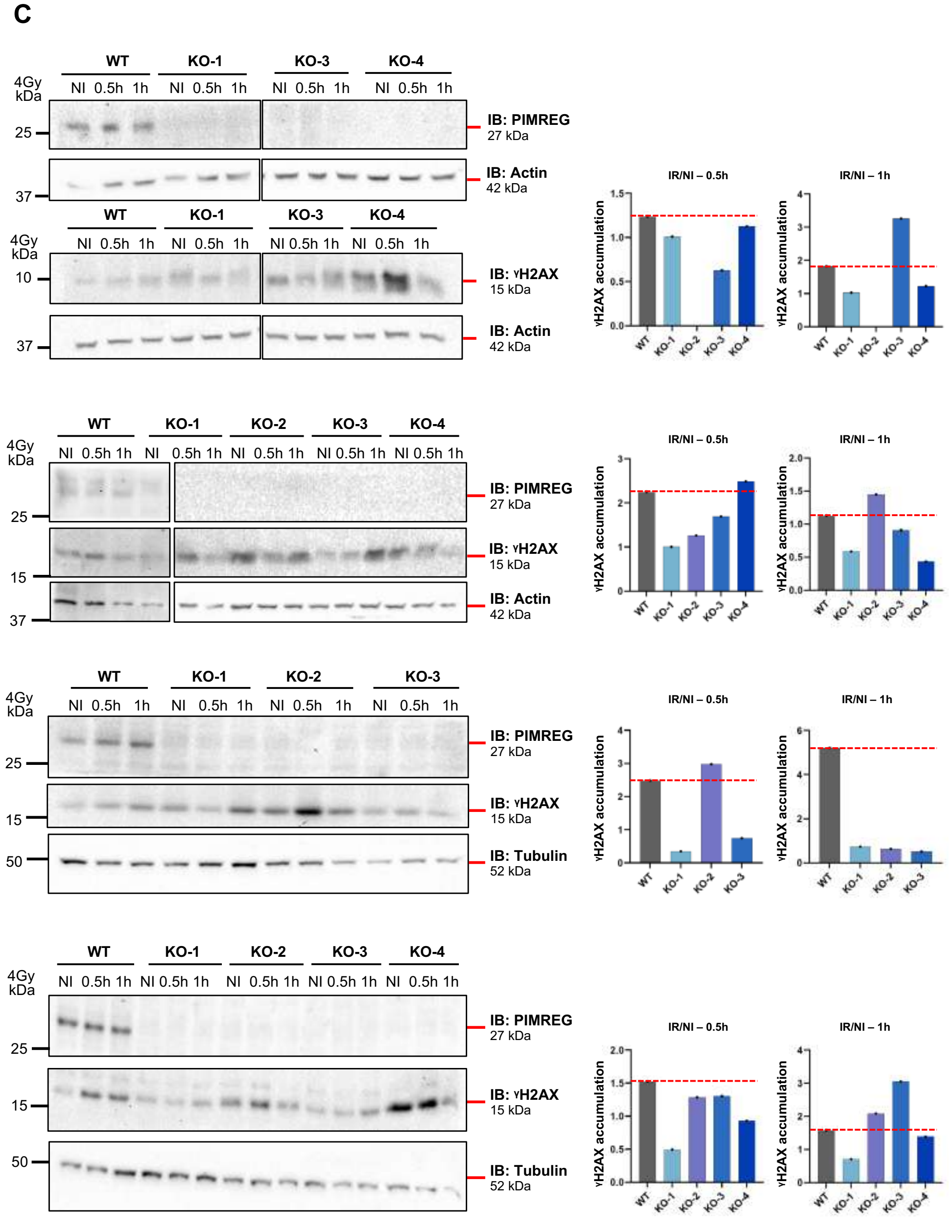

**Fig. S2C: Western blot analysis of γH2AX in T98G WT and PIMREG KO cells treated with IR.** Western blot analysis of protein extracts derived from T98G WT and PIMREG KOs following exposure to IR (4 Gy) and collected 0.5 and 1 h post-irradiation. Membranes were blotted with anti-PIMREG, anti-phospho-H2AX (S139) antibodies. Tubulin was used as a loading control. On the right, bar graphs showing γH2AX accumulation expressed as the ratio of γH2AX levels in IR-treated cells relative to their corresponding non-irradiated controls (IR/NI) for each individual blot at the indicated time points after IR exposure. Red dashed line indicates the γH2AX accumulation in WT cells following IR exposure. Four independent experiments are shown.

**Supplementary Figure S2:**

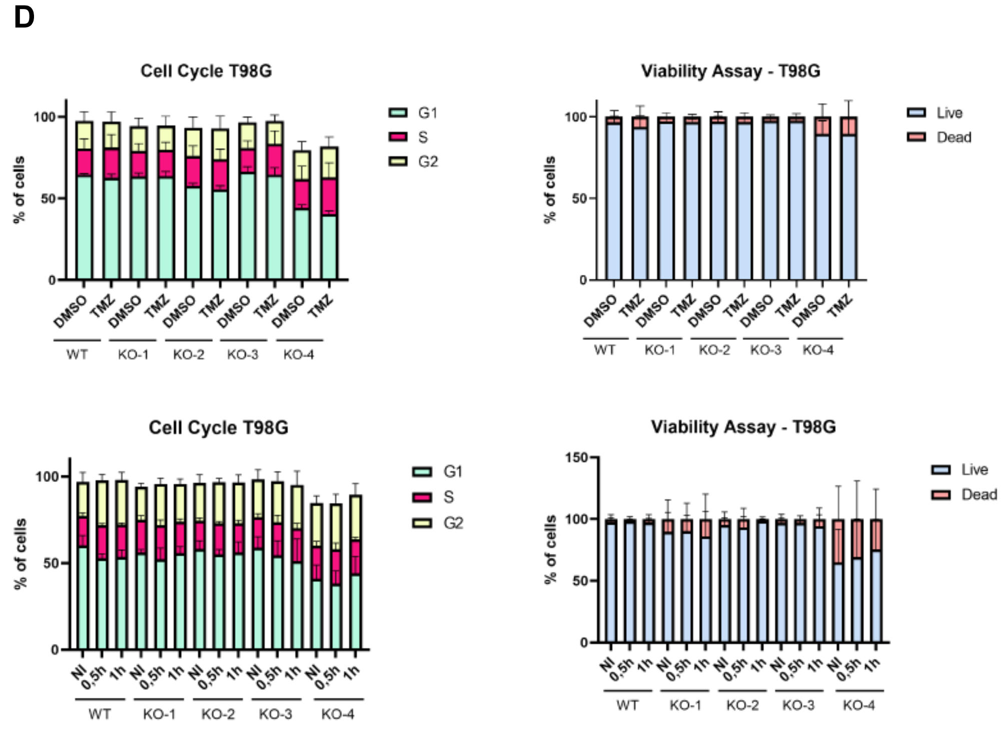


**Fig. S2D:** Cell cycle (left) and viability assays (right) from T98G WT and PIMREG KO (KO-1, KO-2, KO-3 and KO-4) cells treated with 300 µM TMZ or vehicle (DMSO) for 24 h (upper panels) and exposed to 4 Gy IR (lower panels). The same samples of cell extracts in B and C, were analyzed in D. Cell cycle and viability were comparable between all groups of cells in each condition. Results are shown as mean ± SD (standard deviation) of five (TMZ-treated) and four (IR-treated) independent experiments. WT: wild-type; KO: knockout; NI: non-irradiated.

**Supplementary Figure S3:**

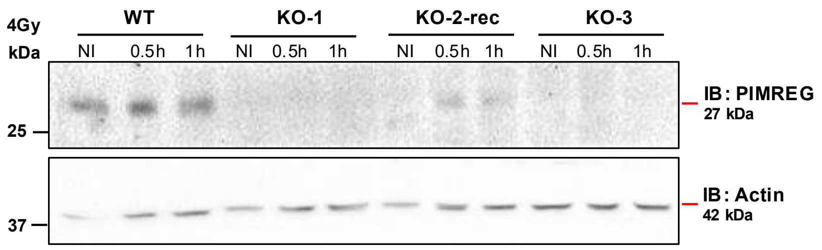


**Fig. S3:** PIMREG expression is restored in PIMREG KO-2 cell line (KO-2-recPIMREG). Western blot analysis of PIMREG in the same samples as Fig. 5C and 5D. Samples are T98G WT, PIMREG KOs (KO-1 and KO-3) and KO-2-recPIMREG. Membranes were blotted with anti-PIMREG and actin was used as a loading control.

**Supplementary Figure S4**

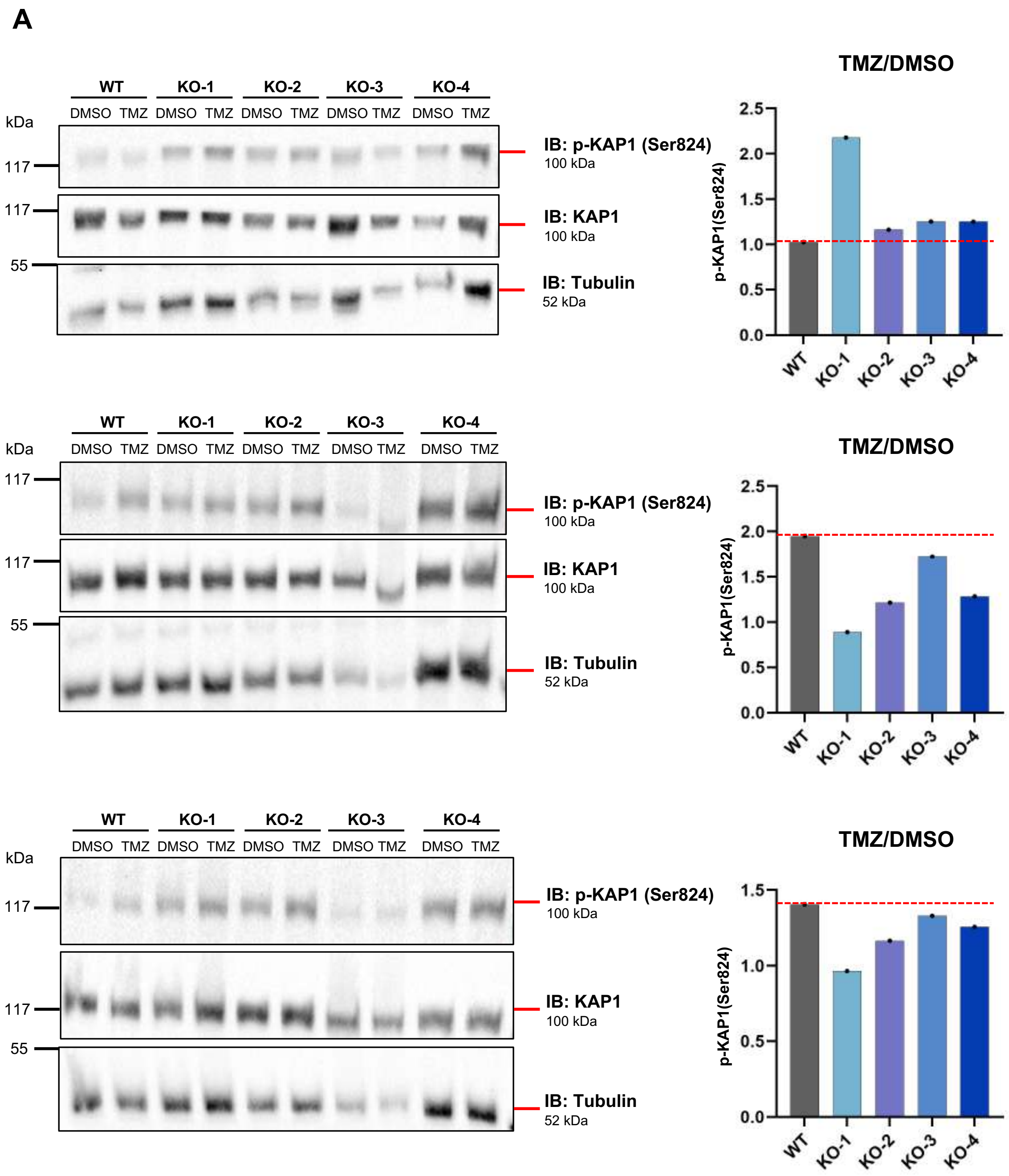

**B**

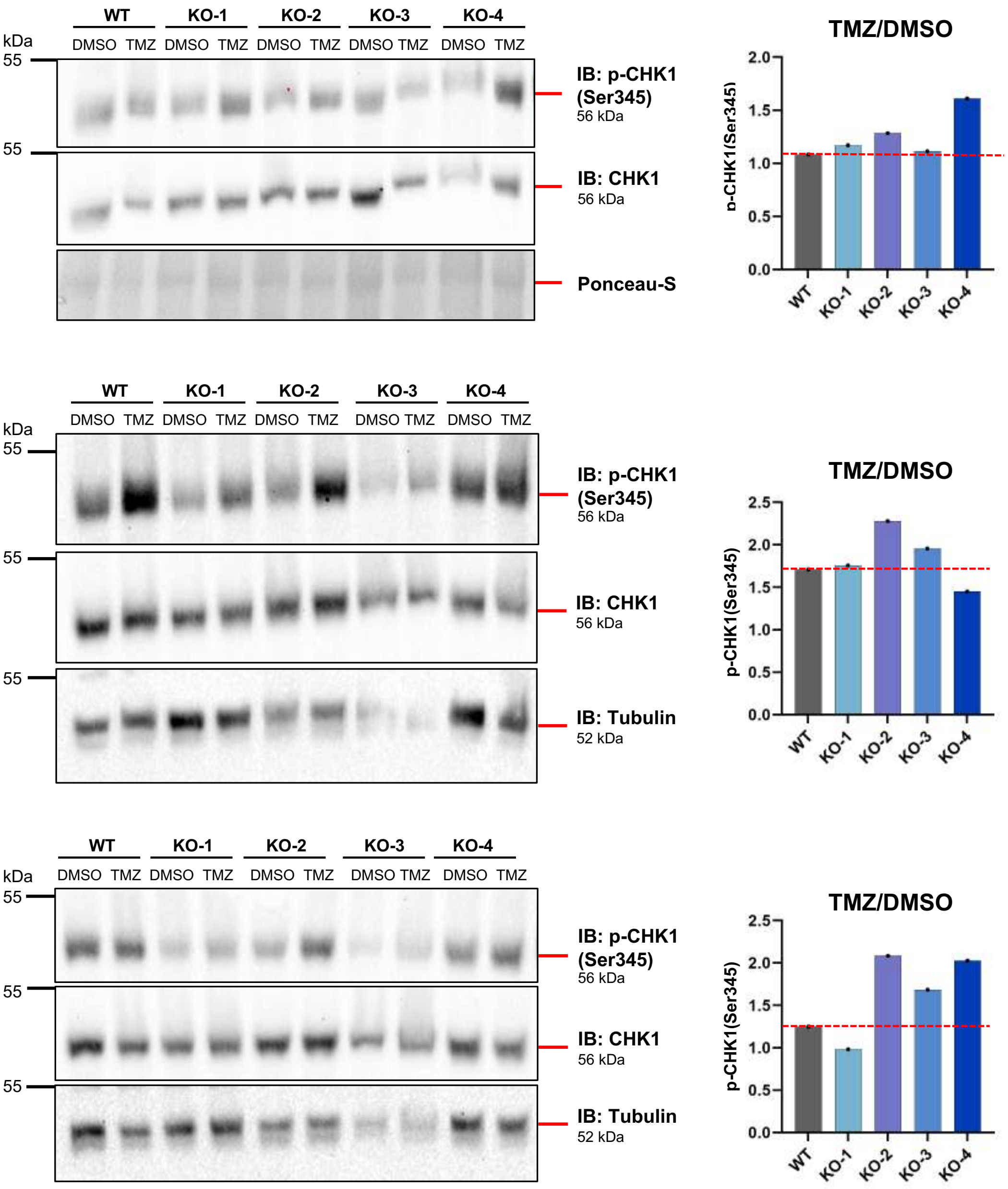

WT
KO-1
KO-2
KO-3
KO-4
DMSO TMZ DMSO TMZ DMSO TMZ DMSO TMZ DMSO TMZ
kDa
55
IB: p-CHK1 (Ser345)
56 kDa
IB: CHK1
56 kDa
Ponceau-S
IB: Tubulin
52 kDa
TMZ/DMSO
p-CHK1(Ser345)
0.0
0.5
1.0
1.5
2.0
2.5

C

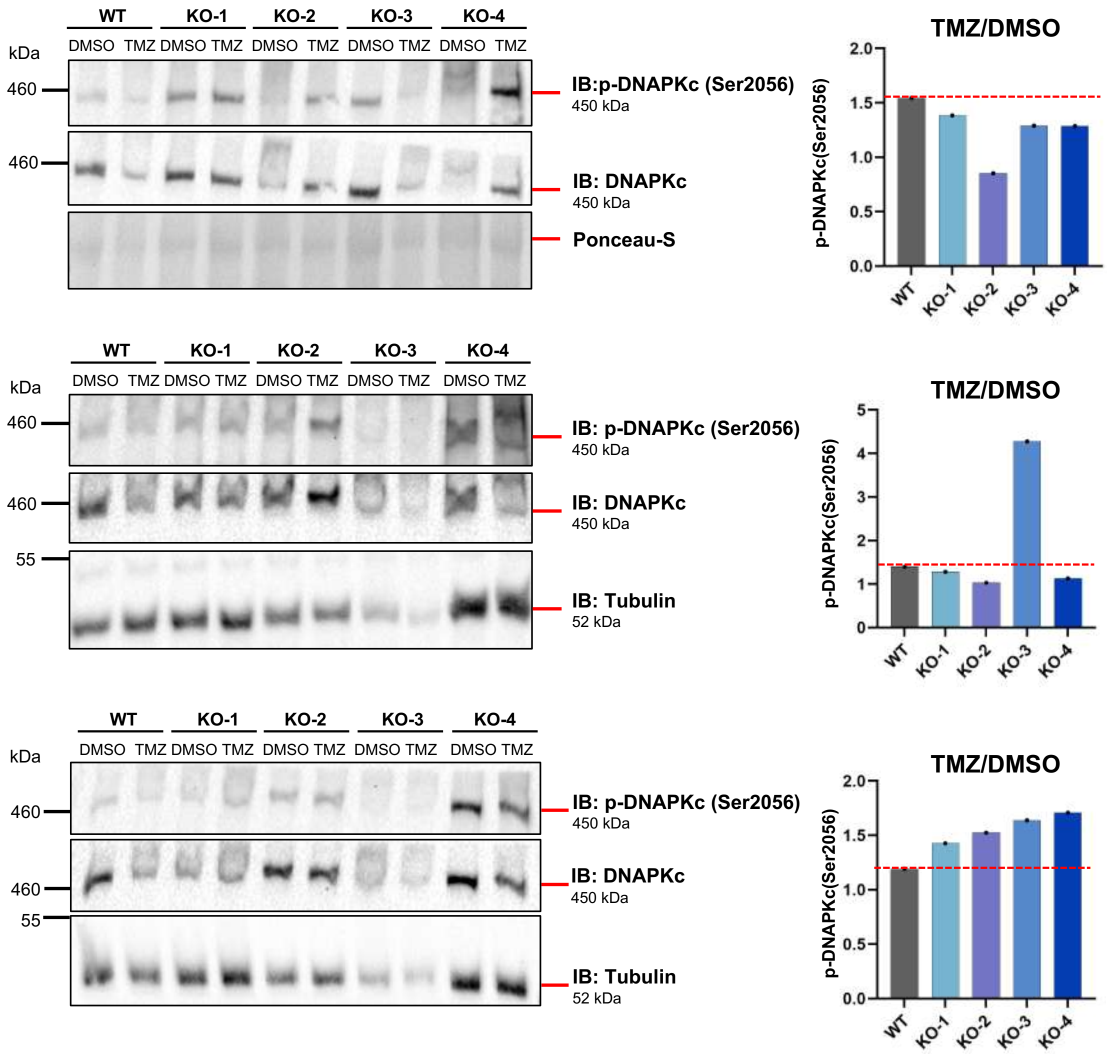


**Fig. S4A-C: Western blot analysis of DNA damage response signaling molecules in T98G WT and PIMREG KO cells treated with TMZ.** Western blot analysis of protein extracts derived from T98G WT and PIMREG KOs treated with 300 µM TMZ or vehicle (DMSO) for 24 h. Membranes were blotted with (A)anti-Phospho-KAP1 (Ser824) and KAP1, (B) anti-Phospho CHK1 (Ser345) and CHK1 and (C) anti-Phospho-DNA-PKc (Ser2056) and DNA-PKc antibodies. Tubulin was used as a loading control. On the right, bar graph showing densitometric quantification of phosphorylated protein band intensities normalized to total protein levels. Protein activation is expressed as the ratio of the phosphorylation levels in TMZ-treated cells relative to their corresponding DMSO control (TMZ/DMSO) for each individual blot. Red dashed line indicates the protein activation in WT cells upon TMZ treatment. Three independent experiments are shown.

**D**

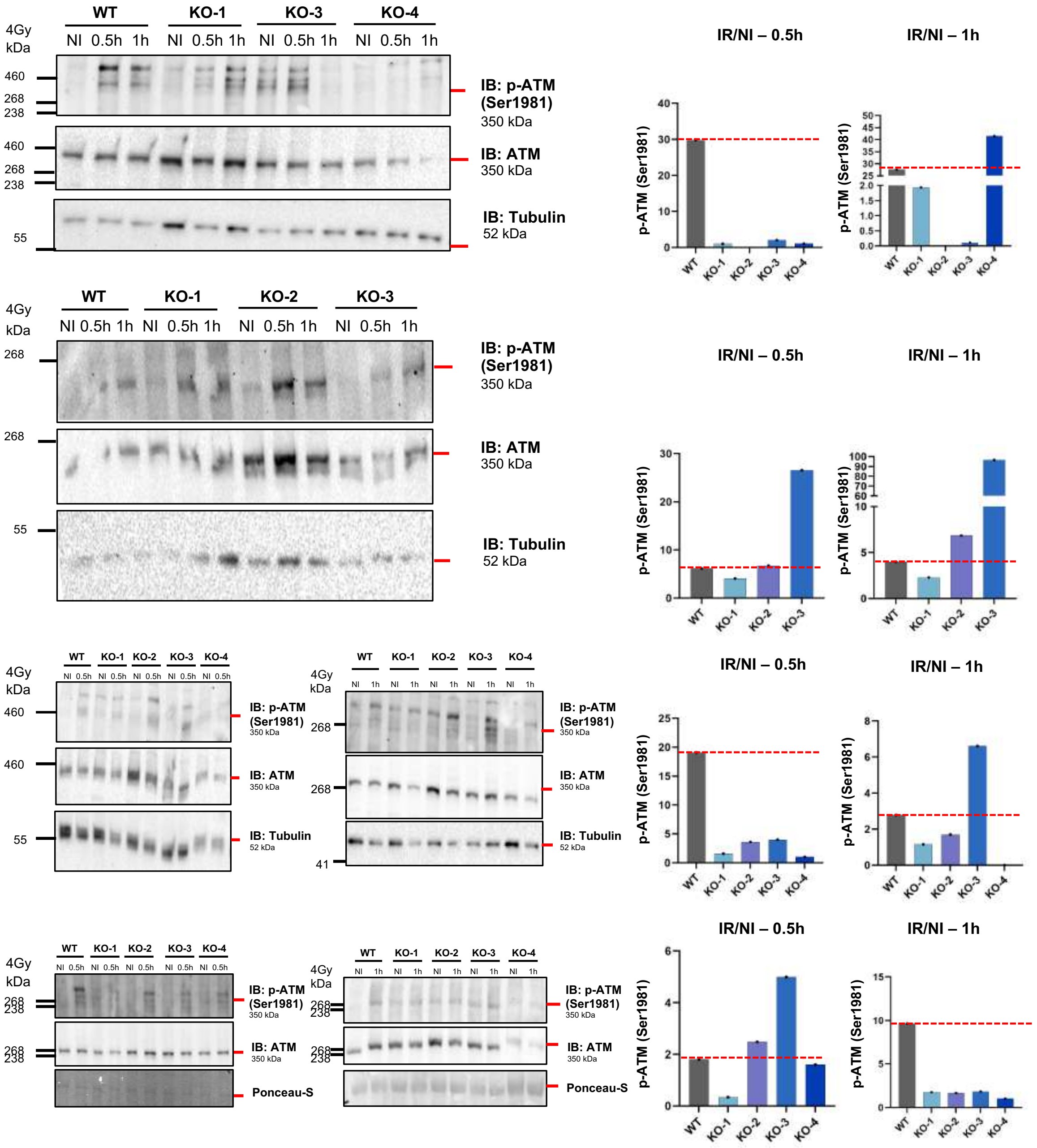

WT
KO-1
KO-2
KO-3
KO-4
4Gy
kDa
NI 0.5h 1h
460
268
238
55
41
IB: p-ATM (Ser1981)
350 kDa
IB: ATM
350 kDa
IB: Tubulin
52 kDa
Ponceau-S
IR/NI – 0.5h
IR/NI – 1h
p-ATM (Ser1981)

**E**

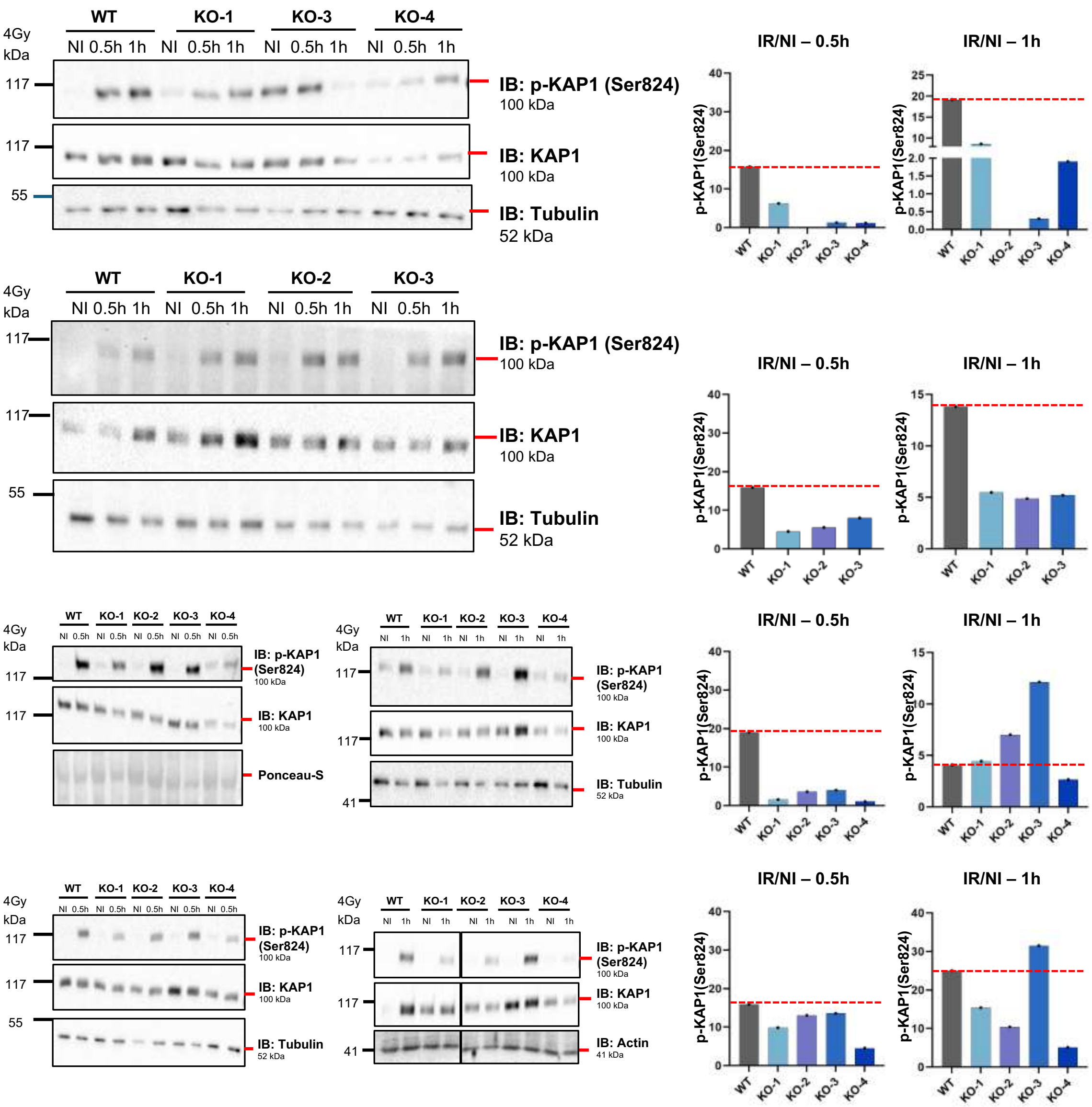

4Gy
kDa
WT
KO-1
KO-2
KO-3
KO-4
NI
0.5h
1h
117
55
41
IB: p-KAP1 (Ser824)
100 kDa
IB: KAP1
100 kDa
IB: Tubulin
52 kDa
Ponceau-S
IB: Actin
41 kDa
IR/NI – 0.5h
IR/NI – 1h
p-KAP1(Ser824)

F

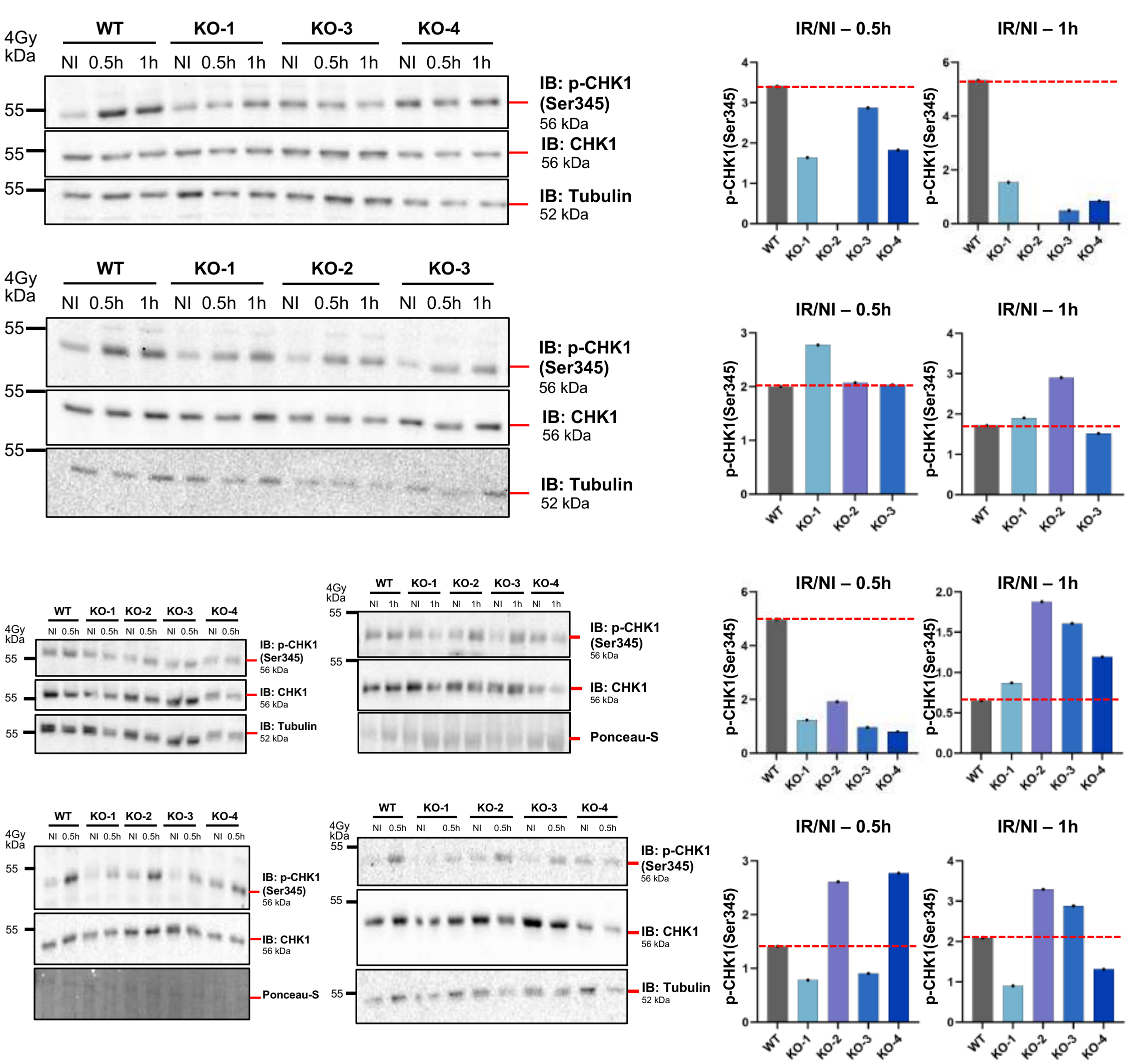
WT
KO-1
KO-3
KO-4
4Gy
kDa
NI 0.5h 1h NI 0.5h 1h NI 0.5h 1h NI 0.5h 1h
55
IB: p-CHK1 (Ser345) 56 kDa
IB: CHK1 56 kDa
IB: Tubulin 52 kDa
KO-2
Ponceau-S
IR/NI – 0.5h
IR/NI – 1h
p-CHK1(Ser345)

**G**

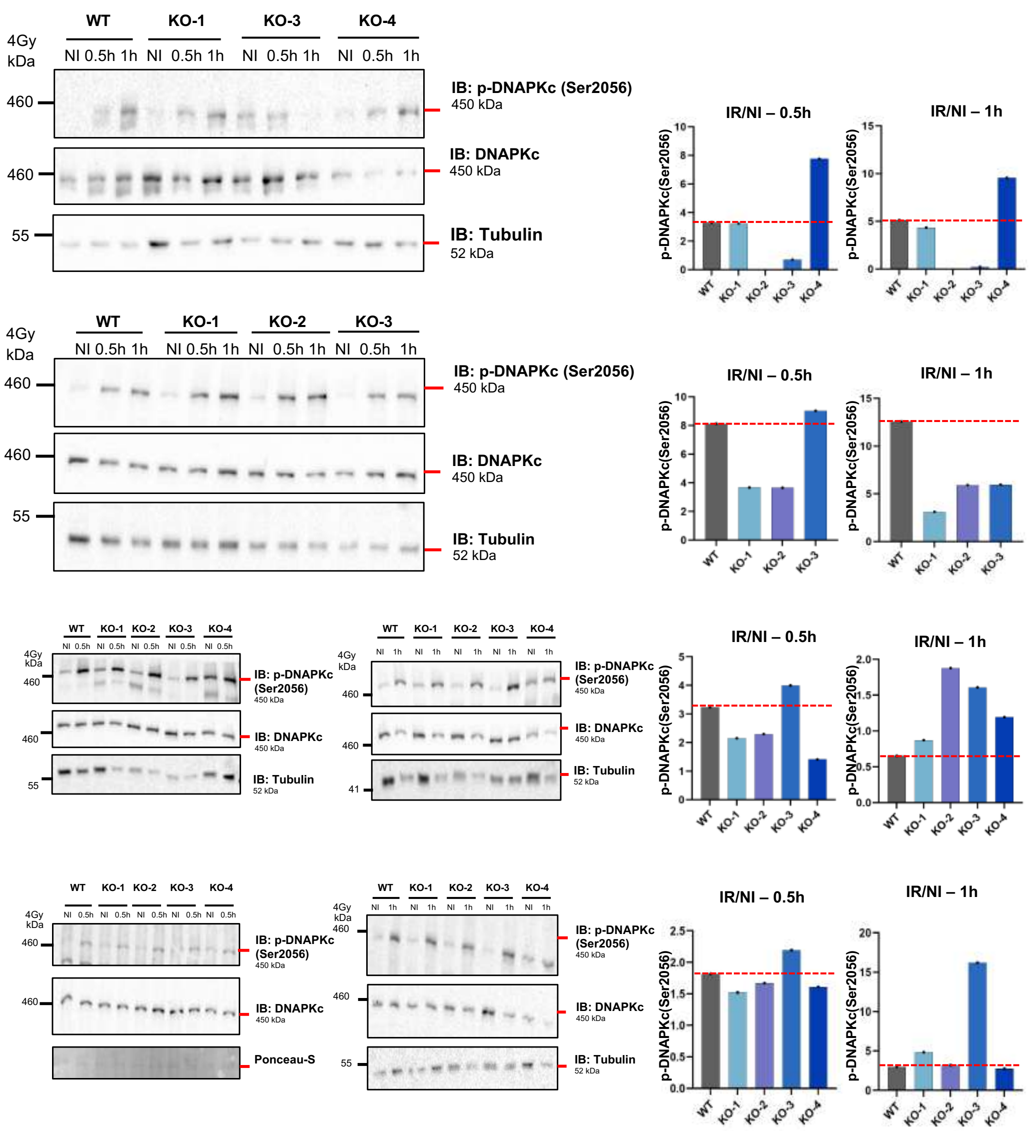


**Fig. S4D-G: Western blot analysis of DNA damage response signaling molecules T98G WT and PIMREG KO cells exposed to IR.** Western blot analysis of protein extracts from T98G WT and PIMREG KO cells (KO-1, KO-2, KO-3, and KO-4) following exposure to IR (4 Gy). Samples were collected at 0.5 and 1h post-irradiation. Membranes were blotted with (**D**) anti-Phospho-ATM (Ser1981) and ATM, (**E**) anti-Phospho-KAP1 (Ser824) and KAP1, (**F**) anti-Phospho CHK1 (Ser345) and CHK1, (**G**) anti-Phospho-DNA-PKc (Ser2056) and DNA-PKc antibodies. Tubulin was used as a loading control. On the right, bar graph showing densitometric quantification of phosphorylated protein band intensities normalized to total protein levels. Protein activation is expressed as the ratio of the phosphorylation levels in IR samples relative to their corresponding non-irradiated controls (IR/NI) for each individual blot. Red dashed line indicates the protein activation in WT cells at the indicated time points following IR exposure. Four independent experiments are shown (*) $p$-value < 0.05 and (**) $p$-value < 0.01, One-way ANOVA test. WT: wild-type; KO: knockout; NI: non-irradiated.

**Supplementary Table 1.** GSEA analysis according to *PIMREG* expression in GBM cohort (TCGA-PanCancer).

| Database | Gene Set | NES | NOM *p*-value | FDR q-value |
|---|---|---|---|---|
| HALLMARK | E2F_TARGETS | 2.107 | 0.000 | 0.002 |
| HALLMARK | MYC_TARGETS_V1 | 2.019 | 0.000 | 0.005 |
| HALLMARK | DNA_REPAIR | 2.015 | 0.000 | 0.004 |
| HALLMARK | G2M_CHECKPOINT | 2.007 | 0.002 | 0.003 |
| HALLMARK | MYC_TARGETS_V2 | 1.781 | 0.019 | 0.041 |
| HALLMARK | SPERMATOGENESIS | 1.582 | 0.027 | 0.132 |
| HALLMARK | UV_RESPONSE_DN | -1.883 | 0.000 | 0.122 |
| HALLMARK | ESTROGEN_RESPONSE_EARLY | -1.671 | 0.008 | 0.127 |
| HALLMARK | TGF_BETA_SIGNALING | -1.678 | 0.018 | 0.182 |
| HALLMARK | INFLAMMATORY_RESPONSE | -1.788 | 0.024 | 0.161 |
| HALLMARK | IL2_STAT5_SIGNALING | -1.673 | 0.025 | 0.149 |
| HALLMARK | COMPLEMENT | -1.659 | 0.026 | 0.104 |
| HALLMARK | IL6_JAK_STAT3_SIGNALING | -1.710 | 0.026 | 0.192 |
| HALLMARK | HEME_METABOLISM | -1.500 | 0.027 | 0.136 |
| HALLMARK | MYOGENESIS | -1.529 | 0.032 | 0.125 |
| HALLMARK | TNFA_SIGNALING_VIA_NFKB | -1.642 | 0.048 | 0.100 |
| HALLMARK | ANGIOGENESIS | -1.562 | 0.049 | 0.134 |
| KEGG | CELL CYCLE | 2.140 | 0.000 | 0.016 |
| KEGG | OOCYTE MEIOSIS | 1.856 | 0.002 | 0.072 |
| KEGG | PROGESTERONE-MEDIATED OOCYTE MATURATION | 1.753 | 0.002 | 0.101 |
| KEGG | SPLICEOSOME | 1.915 | 0.002 | 0.136 |
| KEGG | RNA POLYMERASE | 1.875 | 0.002 | 0.071 |
| KEGG | BASE EXCISION REPAIR | 1.849 | 0.004 | 0.067 |
| KEGG | DNA REPLICATION | 1.908 | 0.004 | 0.074 |
| KEGG | FANCONI ANEMIA PATHWAY | 1.821 | 0.005 | 0.076 |
| KEGG | NUCLEOTIDE EXCISION REPAIR | 1.797 | 0.014 | 0.085 |
| KEGG | THERMOGENESIS | 1.910 | 0.014 | 0.095 |
| KEGG | PARKINSON DISEASE | 1.875 | 0.015 | 0.085 |
| KEGG | PYRIMIDINE METABOLISM | 1.664 | 0.021 | 0.130 |
| KEGG | RNA TRANSPORT | 1.687 | 0.028 | 0.126 |
| KEGG | MISMATCH REPAIR | 1.691 | 0.030 | 0.140 |
| KEGG | PROTEASOME | 1.690 | 0.032 | 0.131 |
| KEGG | OXIDATIVE PHOSPHORYLATION | 1.741 | 0.036 | 0.102 |
| KEGG | HUNTINGTON DISEASE | 1.768 | 0.040 | 0.099 |

| | | | | |
|---|---|---|---|---|
| KEGG | RNA DEGRADATION | 1.566 | 0.042 | 0.216 |
| KEGG | RIBOSOME | 1.684 | 0.046 | 0.120 |
| KEGG | BASAL TRANSCRIPTION FACTORS | 1.557 | 0.049 | 0.217 |
| KEGG | RIG-I-LIKE RECEPTOR SIGNALING PATHWAY | -1.802 | 0.000 | 0.174 |
| KEGG | INSULIN RESISTANCE | -1.694 | 0.000 | 0.138 |
| KEGG | AMOEBIASIS | -1.770 | 0.002 | 0.162 |
| KEGG | PROTEOGLYCANS IN CANCER | -1.682 | 0.003 | 0.139 |
| KEGG | C-TYPE LECTIN RECEPTOR SIGNALING PATHWAY | -1.803 | 0.005 | 0.200 |
| KEGG | LEUKOCYTE TRANSENDOTHELIAL MIGRATION | -1.757 | 0.005 | 0.166 |
| KEGG | PHOSPHATIDYLINOSITOL SIGNALING SYSTEM | -1.700 | 0.005 | 0.145 |
| KEGG | NF-KAPPA B SIGNALING PATHWAY | -1.812 | 0.007 | 0.219 |
| KEGG | OSTEOCLAST DIFFERENTIATION | -1.847 | 0.007 | 0.197 |
| KEGG | TNF SIGNALING PATHWAY | -1.786 | 0.007 | 0.157 |
| KEGG | COMPLEMENT AND COAGULATION CASCADES | -1.798 | 0.009 | 0.157 |
| KEGG | TOLL-LIKE RECEPTOR SIGNALING PATHWAY | -1.682 | 0.010 | 0.144 |
| KEGG | AGE-RAGE SIGNALING PATHWAY IN DIABETIC COMPLICATIONS | -1.650 | 0.010 | 0.148 |
| KEGG | REGULATION OF ACTIN CYTOSKELETON | -1.580 | 0.011 | 0.175 |
| KEGG | ADIPOCYTOKINE SIGNALING PATHWAY | -1.655 | 0.013 | 0.159 |
| KEGG | MUCIN TYPE O-GLYCAN BIOSYNTHESIS | -1.753 | 0.014 | 0.157 |
| KEGG | JAK-STAT SIGNALING PATHWAY | -1.655 | 0.015 | 0.153 |
| KEGG | TOXOPLASMOSIS | -1.724 | 0.016 | 0.170 |
| KEGG | PERTUSSIS | -1.708 | 0.016 | 0.169 |
| KEGG | STARCH AND SUCROSE METABOLISM | -1.703 | 0.016 | 0.149 |
| KEGG | CELL ADHESION MOLECULES (CAMS) | -1.696 | 0.016 | 0.143 |
| KEGG | PANCREATIC SECRETION | -1.536 | 0.017 | 0.201 |
| KEGG | MINERAL ABSORPTION | -1.603 | 0.023 | 0.165 |
| KEGG | INFLAMMATORY BOWEL DISEASE (IBD) | -1.745 | 0.023 | 0.153 |
| KEGG | CYTOKINE-CYTOKINE RECEPTOR INTERACTION | -1.677 | 0.024 | 0.139 |
| KEGG | TH17 CELL DIFFERENTIATION | -1.703 | 0.025 | 0.157 |
| KEGG | MAPK SIGNALING PATHWAY | -1.485 | 0.026 | 0.218 |
| KEGG | HEMATOPOIETIC CELL LINEAGE | -1.721 | 0.026 | 0.163 |
| KEGG | PLATELET ACTIVATION | -1.651 | 0.027 | 0.151 |
| KEGG | ADHERENS JUNCTION | -1.606 | 0.027 | 0.166 |
| KEGG | HISTIDINE METABOLISM | -1.643 | 0.028 | 0.144 |
| KEGG | CHAGAS DISEASE (AMERICAN TRYPANOSOMIASIS) | -1.597 | 0.028 | 0.165 |
| KEGG | STAPHYLOCOCCUS AUREUS INFECTION | -1.703 | 0.036 | 0.166 |
| KEGG | OVARIAN STEROIDOGENESIS | -1.560 | 0.038 | 0.191 |

| | | | | |
|---|---|---|---|---|
| KEGG | VITAMIN DIGESTION AND ABSORPTION | -1.563 | 0.038 | 0.192 |
| KEGG | LEISHMANIASIS | -1.646 | 0.038 | 0.147 |
| KEGG | FOCAL ADHESION | -1.485 | 0.039 | 0.221 |
| KEGG | GASTRIC ACID SECRETION | -1.544 | 0.039 | 0.195 |
| KEGG | INFLUENZA A | -1.551 | 0.042 | 0.191 |
| KEGG | TH1 AND TH2 CELL DIFFERENTIATION | -1.618 | 0.043 | 0.163 |
| KEGG | ECM-RECEPTOR INTERACTION | -1.553 | 0.043 | 0.194 |
| KEGG | INTESTINAL IMMUNE NETWORK FOR IGA PRODUCTION | -1.610 | 0.043 | 0.166 |
| KEGG | TUBERCULOSIS | -1.589 | 0.048 | 0.170 |
| KEGG | SALIVARY SECRETION | -1.466 | 0.048 | 0.221 |
| KEGG | KAPOSI SARCOMA-ASSOCIATED HERPESVIRUS INFECTION | -1.478 | 0.049 | 0.222 |
| REACTOME | SEPARATION OF SISTER CHROMATIDS_HOMO SAPIENS_R-HSA-2467813 | 2.212 | 0.000 | 0.007 |
| REACTOME | CELL CYCLE CHECKPOINTS_HOMO SAPIENS_R-HSA-69620 | 2.185 | 0.000 | 0.008 |
| REACTOME | M PHASE_HOMO SAPIENS_R-HSA-68886 | 2.216 | 0.000 | 0.008 |
| REACTOME | MITOTIC ANAPHASE_HOMO SAPIENS_R-HSA-68882 | 2.223 | 0.000 | 0.009 |
| REACTOME | MITOTIC METAPHASE AND ANAPHASE_HOMO SAPIENS_R-HSA-2555396 | 2.225 | 0.000 | 0.011 |
| REACTOME | RESOLUTION OF SISTER CHROMATID COHESION_HOMO SAPIENS_R-HSA-2500257 | 2.129 | 0.000 | 0.012 |
| REACTOME | RHO GTPASES ACTIVATE FORMINS_HOMO SAPIENS_R-HSA-5663220 | 2.112 | 0.000 | 0.013 |
| REACTOME | MITOTIC PROMETAPHASE_HOMO SAPIENS_R-HSA-68877 | 2.114 | 0.000 | 0.013 |
| REACTOME | MITOTIC G1-G1/S PHASES_HOMO SAPIENS_R-HSA-453279 | 2.131 | 0.000 | 0.013 |
| REACTOME | MITOTIC G2-G2/M PHASES_HOMO SAPIENS_R-HSA-453274 | 2.235 | 0.000 | 0.014 |
| REACTOME | RNA POLYMERASE III TRANSCRIPTION INITIATION_HOMO SAPIENS_R-HSA-76046 | 2.092 | 0.000 | 0.014 |
| REACTOME | G2/M CHECKPOINTS_HOMO SAPIENS_R-HSA-69481 | 2.098 | 0.000 | 0.014 |
| REACTOME | E2F MEDIATED REGULATION OF DNA REPLICATION_HOMO SAPIENS_R-HSA-113510 | 2.135 | 0.000 | 0.015 |
| REACTOME | S PHASE_HOMO SAPIENS_R-HSA-69242 | 2.078 | 0.000 | 0.016 |
| REACTOME | DNA REPLICATION_HOMO SAPIENS_R-HSA-69306 | 2.058 | 0.000 | 0.017 |
| REACTOME | G1/S TRANSITION_HOMO SAPIENS_R-HSA-69206 | 2.053 | 0.000 | 0.017 |
| REACTOME | DNA REPAIR_HOMO SAPIENS_R-HSA-73894 | 2.058 | 0.000 | 0.018 |
| REACTOME | CYCLIN D ASSOCIATED EVENTS IN G1_HOMO SAPIENS_R-HSA-69231 | 2.023 | 0.000 | 0.019 |
| REACTOME | CYCLIN A/B1 ASSOCIATED EVENTS DURING G2/M TRANSITION_HOMO SAPIENS_R-HSA-69273 | 2.042 | 0.000 | 0.019 |
| REACTOME | CHROMOSOME MAINTENANCE_HOMO SAPIENS_R-HSA-73886 | 2.019 | 0.002 | 0.019 |
| REACTOME | TRANSCRIPTION-COUPLED NUCLEOTIDE EXCISION REPAIR (TC-NER)_HOMO SAPIENS_R-HSA-6781827 | 2.037 | 0.000 | 0.019 |
| REACTOME | APC-CDC20 MEDIATED DEGRADATION OF NEK2A_HOMO SAPIENS_R-HSA-179409 | 1.961 | 0.000 | 0.019 |
| REACTOME | G1 PHASE_HOMO SAPIENS_R-HSA-69236 | 2.023 | 0.000 | 0.019 |
| REACTOME | PROCESSING OF CAPPED INTRON-CONTAINING PRE-MRNA_HOMO SAPIENS_R-HSA-72203 | 1.961 | 0.002 | 0.020 |
| REACTOME | REGULATION OF TP53 EXPRESSION AND DEGRADATION_HOMO SAPIENS_R-HSA-6806003 | 1.962 | 0.000 | 0.020 |
| REACTOME | G2/M TRANSITION_HOMO SAPIENS_R-HSA-69275 | 2.239 | 0.000 | 0.020 |

| | | | | |
|---|---|---|---|---|
| REACTOME | M/G1 TRANSITION_HOMO SAPIENS_R-HSA-68874 | 1.964 | 0.000 | 0.020 |
| REACTOME | MRNA SPLICING_HOMO SAPIENS_R-HSA-72172 | 2.005 | 0.000 | 0.020 |
| REACTOME | RNA POLYMERASE III TRANSCRIPTION INITIATION FROM TYPE 1 PROMOTER_HOMO SAPIENS_R-HSA-76061 | 2.024 | 0.000 | 0.020 |
| REACTOME | DNA REPLICATION PRE-INITIATION_HOMO SAPIENS_R-HSA-69002 | 1.964 | 0.000 | 0.020 |
| REACTOME | DUAL INCISION IN TC-NER_HOMO SAPIENS_R-HSA-6782135 | 1.965 | 0.002 | 0.020 |
| REACTOME | RNA POLYMERASE III TRANSCRIPTION_HOMO SAPIENS_R-HSA-74158 | 1.966 | 0.000 | 0.020 |
| REACTOME | COOPERATION OF PREFOLDIN AND TRIC/CCT IN ACTIN AND TUBULIN FOLDING_HOMO SAPIENS_R-HSA-389958 | 2.007 | 0.000 | 0.020 |
| REACTOME | REGULATION OF TP53 DEGRADATION_HOMO SAPIENS_R-HSA-6804757 | 2.026 | 0.000 | 0.021 |
| REACTOME | INACTIVATION OF APC/C VIA DIRECT INHIBITION OF THE APC/C COMPLEX_HOMO SAPIENS_R-HSA-141430 | 1.986 | 0.000 | 0.021 |
| REACTOME | GAP-FILLING DNA REPAIR SYNTHESIS AND LIGATION IN TC-NER_HOMO SAPIENS_R-HSA-6782210 | 1.998 | 0.002 | 0.021 |
| REACTOME | RNA POLYMERASE III ABORTIVE AND RETRACTIVE INITIATION_HOMO SAPIENS_R-HSA-749476 | 1.966 | 0.000 | 0.021 |
| REACTOME | REGULATION OF MITOTIC CELL CYCLE_HOMO SAPIENS_R-HSA-453276 | 1.968 | 0.000 | 0.021 |
| REACTOME | MRNA SPLICING - MINOR PATHWAY_HOMO SAPIENS_R-HSA-72165 | 2.008 | 0.000 | 0.021 |
| REACTOME | NUCLEOTIDE EXCISION REPAIR_HOMO SAPIENS_R-HSA-5696398 | 1.975 | 0.004 | 0.021 |
| REACTOME | INHIBITION OF THE PROTEOLYTIC ACTIVITY OF APC/C REQUIRED FOR THE ONSET OF ANAPHASE BY MITOTIC SPINDLE CHECKPOINT COMPONENTS_HOMO SAPIENS_R-HSA-141405 | 1.986 | 0.000 | 0.021 |
| REACTOME | FGFR2 ALTERNATIVE SPLICING_HOMO SAPIENS_R-HSA-6803529 | 1.992 | 0.000 | 0.021 |
| REACTOME | MITOTIC SPINDLE CHECKPOINT_HOMO SAPIENS_R-HSA-69618 | 1.971 | 0.002 | 0.021 |
| REACTOME | RNA POLYMERASE III TRANSCRIPTION INITIATION FROM TYPE 2 PROMOTER_HOMO SAPIENS_R-HSA-76066 | 1.993 | 0.000 | 0.021 |
| REACTOME | PREFOLDIN MEDIATED TRANSFER OF SUBSTRATE TO CCT/TRIC_HOMO SAPIENS_R-HSA-389957 | 1.976 | 0.000 | 0.021 |
| REACTOME | CELL CYCLE, MITOTIC_HOMO SAPIENS_R-HSA-69278 | 2.274 | 0.000 | 0.021 |
| REACTOME | RNA POLYMERASE III TRANSCRIPTION INITIATION FROM TYPE 3 PROMOTER_HOMO SAPIENS_R-HSA-76071 | 1.986 | 0.000 | 0.021 |
| REACTOME | SYNTHESIS OF DNA_HOMO SAPIENS_R-HSA-69239 | 1.989 | 0.000 | 0.021 |
| REACTOME | APC/C-MEDIATED DEGRADATION OF CELL CYCLE PROTEINS_HOMO SAPIENS_R-HSA-174143 | 1.968 | 0.000 | 0.021 |
| REACTOME | MRNA SPLICING - MAJOR PATHWAY_HOMO SAPIENS_R-HSA-72163 | 1.976 | 0.000 | 0.022 |
| REACTOME | PHOSPHORYLATION OF THE APC/C_HOMO SAPIENS_R-HSA-176412 | 1.946 | 0.000 | 0.023 |
| REACTOME | FORMATION OF TUBULIN FOLDING INTERMEDIATES BY CCT/TRIC_HOMO SAPIENS_R-HSA-389960 | 1.936 | 0.000 | 0.023 |
| REACTOME | REGULATION OF APC/C ACTIVATORS BETWEEN G1/S AND EARLY ANAPHASE_HOMO SAPIENS_R-HSA-176408 | 1.936 | 0.002 | 0.024 |
| REACTOME | DNA DOUBLE-STRAND BREAK REPAIR_HOMO SAPIENS_R-HSA-5693532 | 1.938 | 0.005 | 0.024 |
| REACTOME | REGULATION OF PLK1 ACTIVITY AT G2/M TRANSITION_HOMO SAPIENS_R-HSA-2565942 | 1.930 | 0.002 | 0.025 |
| REACTOME | REGULATION OF DNA REPLICATION_HOMO SAPIENS_R-HSA-69304 | 1.922 | 0.002 | 0.027 |
| REACTOME | FORMATION OF HIV-1 ELONGATION COMPLEX CONTAINING HIV-1 TAT_HOMO SAPIENS_R-HSA-167200 | 1.906 | 0.004 | 0.027 |
| REACTOME | RESOLUTION OF ABASIC SITES (AP SITES)_HOMO SAPIENS_R-HSA-73933 | 1.902 | 0.000 | 0.027 |
| REACTOME | TAT-MEDIATED ELONGATION OF THE HIV-1 TRANSCRIPT_HOMO SAPIENS_R-HSA-167246 | 1.906 | 0.004 | 0.028 |
| REACTOME | BASE EXCISION REPAIR_HOMO SAPIENS_R-HSA-73884 | 1.902 | 0.000 | 0.028 |
| REACTOME | APC/C:CDC20 MEDIATED DEGRADATION OF CYCLIN B_HOMO SAPIENS_R-HSA-174048 | 1.910 | 0.000 | 0.028 |

| | | | | |
|---|---|---|---|---|
| REACTOME | CONVERSION FROM APC/C:CDC20 TO APC/C:CDH1 IN LATE ANAPHASE_HOMO SAPIENS_R-HSA-176407 | 1.913 | 0.000 | 0.028 |
| REACTOME | TAT-MEDIATED HIV ELONGATION ARREST AND RECOVERY_HOMO SAPIENS_R-HSA-167243 | 1.915 | 0.002 | 0.028 |
| REACTOME | AURKA ACTIVATION BY TPX2_HOMO SAPIENS_R-HSA-8854518 | 1.909 | 0.002 | 0.028 |
| REACTOME | HOMOLOGY DIRECTED REPAIR_HOMO SAPIENS_R-HSA-5693538 | 1.899 | 0.007 | 0.028 |
| REACTOME | ACTIVATION OF APC/C AND APC/C:CDC20 MEDIATED DEGRADATION OF MITOTIC PROTEINS_HOMO SAPIENS_R-HSA-176814 | 1.911 | 0.004 | 0.028 |
| REACTOME | HIV TRANSCRIPTION ELONGATION_HOMO SAPIENS_R-HSA-167169 | 1.906 | 0.004 | 0.028 |
| REACTOME | APC/C:CDC20 MEDIATED DEGRADATION OF MITOTIC PROTEINS_HOMO SAPIENS_R-HSA-176409 | 1.897 | 0.006 | 0.028 |
| REACTOME | RHO GTPASE EFFECTORS_HOMO SAPIENS_R-HSA-195258 | 1.884 | 0.003 | 0.028 |
| REACTOME | PAUSING AND RECOVERY OF TAT-MEDIATED HIV ELONGATION_HOMO SAPIENS_R-HSA-167238 | 1.915 | 0.002 | 0.028 |
| REACTOME | HDR THROUGH HOMOLOGOUS RECOMBINATION (HR) OR SINGLE STRAND ANNEALING (SSA)_HOMO SAPIENS_R-HSA-5693567 | 1.885 | 0.011 | 0.028 |
| REACTOME | REMOVAL OF LICENSING FACTORS FROM ORIGINS_HOMO SAPIENS_R-HSA-69300 | 1.884 | 0.006 | 0.028 |
| REACTOME | GLOBAL GENOME NUCLEOTIDE EXCISION REPAIR (GG-NER)_HOMO SAPIENS_R-HSA-5696399 | 1.916 | 0.002 | 0.028 |
| REACTOME | ELONGATION ARREST AND RECOVERY_HOMO SAPIENS_R-HSA-112387 | 1.888 | 0.004 | 0.028 |
| REACTOME | FORMATION OF THE HIV-1 EARLY ELONGATION COMPLEX_HOMO SAPIENS_R-HSA-167158 | 1.886 | 0.004 | 0.028 |
| REACTOME | FORMATION OF RNA POL II ELONGATION COMPLEX_HOMO SAPIENS_R-HSA-112382 | 1.878 | 0.004 | 0.029 |
| REACTOME | PAUSING AND RECOVERY OF HIV ELONGATION_HOMO SAPIENS_R-HSA-167290 | 1.888 | 0.004 | 0.029 |
| REACTOME | FORMATION OF THE EARLY ELONGATION COMPLEX_HOMO SAPIENS_R-HSA-113418 | 1.886 | 0.004 | 0.029 |
| REACTOME | FANCONI ANEMIA PATHWAY_HOMO SAPIENS_R-HSA-6783310 | 1.892 | 0.002 | 0.029 |
| REACTOME | NUCLEOSOME ASSEMBLY_HOMO SAPIENS_R-HSA-774815 | 1.890 | 0.008 | 0.029 |
| REACTOME | ABORTIVE ELONGATION OF HIV-1 TRANSCRIPT IN THE ABSENCE OF TAT_HOMO SAPIENS_R-HSA-167242 | 1.880 | 0.002 | 0.029 |
| REACTOME | RNA POLYMERASE II TRANSCRIPTION ELONGATION_HOMO SAPIENS_R-HSA-75955 | 1.878 | 0.004 | 0.029 |
| REACTOME | DEGRADATION OF BETA-CATENIN BY THE DESTRUCTION COMPLEX_HOMO SAPIENS_R-HSA-195253 | 1.893 | 0.004 | 0.029 |
| REACTOME | HIV ELONGATION ARREST AND RECOVERY_HOMO SAPIENS_R-HSA-167287 | 1.888 | 0.004 | 0.029 |
| REACTOME | DEPOSITION OF NEW CENPA-CONTAINING NUCLEOSOMES AT THE CENTROMERE_HOMO SAPIENS_R-HSA-606279 | 1.890 | 0.008 | 0.029 |
| REACTOME | FORMATION OF HIV ELONGATION COMPLEX IN THE ABSENCE OF HIV TAT_HOMO SAPIENS_R-HSA-167152 | 1.878 | 0.004 | 0.029 |
| REACTOME | ORC1 REMOVAL FROM CHROMATIN_HOMO SAPIENS_R-HSA-68949 | 1.870 | 0.008 | 0.030 |
| REACTOME | LOSS OF PROTEINS REQUIRED FOR INTERPHASE MICROTUBULE ORGANIZATION FROM THE CENTROSOME_HOMO SAPIENS_R-HSA-380284 | 1.868 | 0.004 | 0.030 |
| REACTOME | APC:CDC20 MEDIATED DEGRADATION OF CELL CYCLE PROTEINS PRIOR TO SATISFATION OF THE CELL CYCLE CHECKPOINT_HOMO SAPIENS_R-HSA-179419 | 1.866 | 0.010 | 0.030 |
| REACTOME | SWITCHING OF ORIGINS TO A POST-REPLICATIVE STATE_HOMO SAPIENS_R-HSA-69052 | 1.870 | 0.008 | 0.030 |
| REACTOME | FORMATION OF TC-NER PRE-INCISION COMPLEX_HOMO SAPIENS_R-HSA-6781823 | 1.871 | 0.006 | 0.030 |
| REACTOME | LOSS OF NLP FROM MITOTIC CENTROSOMES_HOMO SAPIENS_R-HSA-380259 | 1.868 | 0.004 | 0.030 |
| REACTOME | MICRORNA (MIRNA) BIOGENESIS_HOMO SAPIENS_R-HSA-203927 | 1.866 | 0.002 | 0.031 |
| REACTOME | PROTEIN FOLDING_HOMO SAPIENS_R-HSA-391251 | 1.863 | 0.000 | 0.031 |
| REACTOME | CHAPERONIN-MEDIATED PROTEIN FOLDING_HOMO SAPIENS_R-HSA-390466 | 1.862 | 0.000 | 0.031 |

| | | | | |
|---|---|---|---|---|
| REACTOME | EXTENSION OF TELOMERES_HOMO SAPIENS_R-HSA-180786 | 1.860 | 0.002 | 0.031 |
| REACTOME | HIV INFECTION_HOMO SAPIENS_R-HSA-162906 | 1.858 | 0.007 | 0.032 |
| REACTOME | TRANSCRIPTIONAL REGULATION BY TP53_HOMO SAPIENS_R-HSA-3700989 | 1.853 | 0.002 | 0.032 |
| REACTOME | APC/C:CDH1 MEDIATED DEGRADATION OF CDC20 AND OTHER APC/C:CDH1 TARGETED PROTEINS IN LATE MITOSIS/EARLY G1_HOMO SAPIENS_R-HSA-174178 | 1.853 | 0.008 | 0.033 |
| REACTOME | HIV LIFE CYCLE_HOMO SAPIENS_R-HSA-162587 | 1.849 | 0.004 | 0.033 |
| REACTOME | CDC20:PHOSPHO-APC/C MEDIATED DEGRADATION OF CYCLIN A_HOMO SAPIENS_R-HSA-174184 | 1.848 | 0.010 | 0.034 |
| REACTOME | DNA DAMAGE RECOGNITION IN GG-NER_HOMO SAPIENS_R-HSA-5696394 | 1.836 | 0.009 | 0.035 |
| REACTOME | POLO-LIKE KINASE MEDIATED EVENTS_HOMO SAPIENS_R-HSA-156711 | 1.836 | 0.004 | 0.036 |
| REACTOME | MRNA CAPPING_HOMO SAPIENS_R-HSA-72086 | 1.836 | 0.006 | 0.036 |
| REACTOME | VIRAL MESSENGER RNA SYNTHESIS_HOMO SAPIENS_R-HSA-168325 | 1.832 | 0.007 | 0.036 |
| REACTOME | TP53 REGULATES TRANSCRIPTION OF DNA REPAIR GENES_HOMO SAPIENS_R-HSA-6796648 | 1.836 | 0.004 | 0.036 |
| REACTOME | G1/S DNA DAMAGE CHECKPOINTS_HOMO SAPIENS_R-HSA-69615 | 1.831 | 0.013 | 0.036 |
| REACTOME | G0 AND EARLY G1_HOMO SAPIENS_R-HSA-1538133 | 1.830 | 0.011 | 0.036 |
| REACTOME | RNA POL II CTD PHOSPHORYLATION AND INTERACTION WITH CE_HOMO SAPIENS_R-HSA-77075 | 1.836 | 0.006 | 0.037 |
| REACTOME | RNA POLYMERASE III TRANSCRIPTION TERMINATION_HOMO SAPIENS_R-HSA-73980 | 1.828 | 0.004 | 0.037 |
| REACTOME | RNA POL II CTD PHOSPHORYLATION AND INTERACTION WITH CE_HOMO SAPIENS_R-HSA-167160 | 1.836 | 0.006 | 0.037 |
| REACTOME | TRANSLESION SYNTHESIS BY Y FAMILY DNA POLYMERASES BYPASSES LESIONS ON DNA TEMPLATE_HOMO SAPIENS_R-HSA-110313 | 1.838 | 0.007 | 0.037 |
| REACTOME | RNA POLYMERASE III CHAIN ELONGATION_HOMO SAPIENS_R-HSA-73780 | 1.837 | 0.000 | 0.037 |
| REACTOME | HDR THROUGH HOMOLOGOUS RECOMBINATION (HRR)_HOMO SAPIENS_R-HSA-5685942 | 1.823 | 0.018 | 0.038 |
| REACTOME | ASSEMBLY OF THE PRE-REPLICATIVE COMPLEX_HOMO SAPIENS_R-HSA-68867 | 1.824 | 0.010 | 0.038 |
| REACTOME | INFECTIOUS DISEASE_HOMO SAPIENS_R-HSA-5663205 | 1.816 | 0.035 | 0.040 |
| REACTOME | PIWI-INTERACTING RNA (PIRNA) BIOGENESIS_HOMO SAPIENS_R-HSA-5601884 | 1.811 | 0.011 | 0.041 |
| REACTOME | HOST INTERACTIONS OF HIV FACTORS_HOMO SAPIENS_R-HSA-162909 | 1.808 | 0.011 | 0.041 |
| REACTOME | COOPERATION OF PDCL (PHLP1) AND TRIC/CCT IN G-PROTEIN BETA FOLDING_HOMO SAPIENS_R-HSA-6814122 | 1.809 | 0.002 | 0.041 |
| REACTOME | RESOLUTION OF AP SITES VIA THE MULTIPLE-NUCLEOTIDE PATCH REPLACEMENT PATHWAY_HOMO SAPIENS_R-HSA-110373 | 1.810 | 0.002 | 0.042 |
| REACTOME | DNA DAMAGE BYPASS_HOMO SAPIENS_R-HSA-73893 | 1.811 | 0.005 | 0.042 |
| REACTOME | SUMOYLATION OF DNA REPLICATION PROTEINS_HOMO SAPIENS_R-HSA-4615885 | 1.805 | 0.013 | 0.042 |
| REACTOME | P53-DEPENDENT G1 DNA DAMAGE RESPONSE_HOMO SAPIENS_R-HSA-69563 | 1.804 | 0.017 | 0.042 |
| REACTOME | P53-DEPENDENT G1/S DNA DAMAGE CHECKPOINT_HOMO SAPIENS_R-HSA-69580 | 1.804 | 0.017 | 0.042 |
| REACTOME | GAP-FILLING DNA REPAIR SYNTHESIS AND LIGATION IN GG-NER_HOMO SAPIENS_R-HSA-5696397 | 1.797 | 0.006 | 0.044 |
| REACTOME | KSRP (KHSRP) BINDS AND DESTABILIZES MRNA_HOMO SAPIENS_R-HSA-450604 | 1.797 | 0.009 | 0.044 |
| REACTOME | ACTIVATION OF ATR IN RESPONSE TO REPLICATION STRESS_HOMO SAPIENS_R-HSA-176187 | 1.793 | 0.009 | 0.045 |
| REACTOME | CYCLIN E ASSOCIATED EVENTS DURING G1/S TRANSITION_HOMO SAPIENS_R-HSA-69202 | 1.791 | 0.017 | 0.045 |
| REACTOME | SCF(SKP2)-MEDIATED DEGRADATION OF P27/P21_HOMO SAPIENS_R-HSA-187577 | 1.792 | 0.013 | 0.045 |
| REACTOME | RECRUITMENT OF MITOTIC CENTROSOME PROTEINS AND COMPLEXES_HOMO SAPIENS_R-HSA-380270 | 1.788 | 0.020 | 0.046 |

| | | | | |
|---|---|---|---|---|
| REACTOME | CENTROSOME MATURATION_HOMO SAPIENS_R-HSA-380287 | 1.788 | 0.020 | 0.046 |
| REACTOME | AUTODEGRADATION OF CDH1 BY CDH1:APC/C_HOMO SAPIENS_R-HSA-174084 | 1.783 | 0.015 | 0.048 |
| REACTOME | DUAL INCISION IN GG-NER_HOMO SAPIENS_R-HSA-5696400 | 1.781 | 0.002 | 0.048 |
| REACTOME | CYCLIN A:CDK2-ASSOCIATED EVENTS AT S PHASE ENTRY_HOMO SAPIENS_R-HSA-69656 | 1.777 | 0.018 | 0.049 |
| REACTOME | THE ROLE OF GTSE1 IN G2/M PROGRESSION AFTER G2 CHECKPOINT_HOMO SAPIENS_R-HSA-8852276 | 1.778 | 0.015 | 0.049 |
| REACTOME | SCF-BETA-TRCP MEDIATED DEGRADATION OF EMI1_HOMO SAPIENS_R-HSA-174113 | 1.772 | 0.017 | 0.049 |
| REACTOME | APC/C:CDC20 MEDIATED DEGRADATION OF SECURIN_HOMO SAPIENS_R-HSA-174154 | 1.774 | 0.015 | 0.050 |
| REACTOME | RNA POLYMERASE II TRANSCRIPTION_HOMO SAPIENS_R-HSA-73857 | 1.771 | 0.015 | 0.050 |
| REACTOME | G1/S-SPECIFIC TRANSCRIPTION_HOMO SAPIENS_R-HSA-69205 | 1.773 | 0.004 | 0.050 |
| REACTOME | TP53 REGULATES TRANSCRIPTION OF CELL CYCLE GENES_HOMO SAPIENS_R-HSA-6791312 | 1.770 | 0.019 | 0.050 |
| REACTOME | REGULATION OF TP53 ACTIVITY THROUGH PHOSPHORYLATION_HOMO SAPIENS_R-HSA-6804756 | 1.774 | 0.024 | 0.050 |
| REACTOME | TELOMERE C-STRAND (LAGGING STRAND) SYNTHESIS_HOMO SAPIENS_R-HSA-174417 | 1.765 | 0.011 | 0.051 |
| REACTOME | PROCESSING OF DNA DOUBLE-STRAND BREAK ENDS_HOMO SAPIENS_R-HSA-5693607 | 1.765 | 0.027 | 0.051 |
| REACTOME | DEADENYLATION-DEPENDENT MRNA DECAY_HOMO SAPIENS_R-HSA-429914 | 1.758 | 0.009 | 0.053 |
| REACTOME | LATE PHASE OF HIV LIFE CYCLE_HOMO SAPIENS_R-HSA-162599 | 1.759 | 0.009 | 0.053 |
| REACTOME | POST-CHAPERONIN TUBULIN FOLDING PATHWAY_HOMO SAPIENS_R-HSA-389977 | 1.757 | 0.009 | 0.053 |
| REACTOME | INTERACTIONS OF VPR WITH HOST CELLULAR PROTEINS_HOMO SAPIENS_R-HSA-176033 | 1.754 | 0.015 | 0.055 |
| REACTOME | TELOMERE MAINTENANCE_HOMO SAPIENS_R-HSA-157579 | 1.749 | 0.035 | 0.056 |
| REACTOME | TP53 REGULATES TRANSCRIPTION OF GENES INVOLVED IN G2 CELL CYCLE ARREST_HOMO SAPIENS_R-HSA-6804114 | 1.749 | 0.011 | 0.056 |
| REACTOME | PCNA-DEPENDENT LONG PATCH BASE EXCISION REPAIR_HOMO SAPIENS_R-HSA-5651801 | 1.750 | 0.005 | 0.056 |
| REACTOME | DEGRADATION OF DVL_HOMO SAPIENS_R-HSA-4641258 | 1.746 | 0.019 | 0.056 |
| REACTOME | MITOCHONDRIAL PROTEIN IMPORT_HOMO SAPIENS_R-HSA-1268020 | 1.741 | 0.026 | 0.058 |
| REACTOME | LAGGING STRAND SYNTHESIS_HOMO SAPIENS_R-HSA-69186 | 1.743 | 0.011 | 0.058 |
| REACTOME | G2/M DNA DAMAGE CHECKPOINT_HOMO SAPIENS_R-HSA-69473 | 1.741 | 0.037 | 0.058 |
| REACTOME | METABOLISM OF POLYAMINES_HOMO SAPIENS_R-HSA-351202 | 1.737 | 0.035 | 0.060 |
| REACTOME | DNA STRAND ELONGATION_HOMO SAPIENS_R-HSA-69190 | 1.734 | 0.013 | 0.060 |
| REACTOME | TRANSLESION SYNTHESIS BY POLI_HOMO SAPIENS_R-HSA-5656121 | 1.733 | 0.004 | 0.061 |
| REACTOME | HIV TRANSCRIPTION INITIATION_HOMO SAPIENS_R-HSA-167161 | 1.723 | 0.019 | 0.061 |
| REACTOME | ASYMMETRIC LOCALIZATION OF PCP PROTEINS_HOMO SAPIENS_R-HSA-4608870 | 1.722 | 0.029 | 0.061 |
| REACTOME | ACTIVATION OF HOX GENES DURING DIFFERENTIATION_HOMO SAPIENS_R-HSA-5619507 | 1.720 | 0.042 | 0.061 |
| REACTOME | GENE SILENCING BY RNA_HOMO SAPIENS_R-HSA-211000 | 1.727 | 0.036 | 0.061 |
| REACTOME | MITOTIC PROPHASE_HOMO SAPIENS_R-HSA-68875 | 1.731 | 0.034 | 0.061 |
| REACTOME | TRNA PROCESSING IN THE NUCLEUS_HOMO SAPIENS_R-HSA-6784531 | 1.717 | 0.026 | 0.061 |
| REACTOME | RNA POLYMERASE II TRANSCRIPTION PRE-INITIATION AND PROMOTER OPENING_HOMO SAPIENS_R-HSA-73779 | 1.723 | 0.019 | 0.061 |
| REACTOME | ACTIVATION OF ANTERIOR HOX GENES IN HINDBRAIN DEVELOPMENT DURING EARLY EMBRYOGENESIS_HOMO SAPIENS_R-HSA-5617472 | 1.720 | 0.042 | 0.061 |

| | | | | |
|---|---|---|---|---|
| REACTOME | COPI-DEPENDENT GOLGI-TO-ER RETROGRADE TRAFFIC_HOMO SAPIENS_R-HSA-6811434 | 1.727 | 0.007 | 0.061 |
| REACTOME | REGULATION OF MRNA STABILITY BY PROTEINS THAT BIND AU-RICH ELEMENTS_HOMO SAPIENS_R-HSA-450531 | 1.718 | 0.036 | 0.061 |
| REACTOME | ORGANELLE BIOGENESIS AND MAINTENANCE_HOMO SAPIENS_R-HSA-1852241 | 1.731 | 0.018 | 0.061 |
| REACTOME | REGULATION OF TP53 ACTIVITY_HOMO SAPIENS_R-HSA-5633007 | 1.720 | 0.015 | 0.062 |
| REACTOME | RNA POLYMERASE II TRANSCRIPTION INITIATION_HOMO SAPIENS_R-HSA-75953 | 1.723 | 0.019 | 0.062 |
| REACTOME | MITOCHONDRIAL TRANSLATION TERMINATION_HOMO SAPIENS_R-HSA-5419276 | 1.728 | 0.020 | 0.062 |
| REACTOME | MITOCHONDRIAL TRANSLATION ELONGATION_HOMO SAPIENS_R-HSA-5389840 | 1.729 | 0.025 | 0.062 |
| REACTOME | RMTS METHYLATE HISTONE ARGININES_HOMO SAPIENS_R-HSA-3214858 | 1.728 | 0.049 | 0.062 |
| REACTOME | REGULATION OF RAS BY GAPS_HOMO SAPIENS_R-HSA-5658442 | 1.715 | 0.035 | 0.062 |
| REACTOME | RNA POLYMERASE II TRANSCRIPTION INITIATION AND PROMOTER CLEARANCE_HOMO SAPIENS_R-HSA-76042 | 1.723 | 0.019 | 0.062 |
| REACTOME | MITOCHONDRIAL TRANSLATION_HOMO SAPIENS_R-HSA-5368287 | 1.703 | 0.033 | 0.062 |
| REACTOME | RECOGNITION OF DNA DAMAGE BY PCNA-CONTAINING REPLICATION COMPLEX_HOMO SAPIENS_R-HSA-110314 | 1.704 | 0.016 | 0.062 |
| REACTOME | RNA POLYMERASE II PROMOTER ESCAPE_HOMO SAPIENS_R-HSA-73776 | 1.723 | 0.019 | 0.062 |
| REACTOME | PROSTACYCLIN SIGNALLING THROUGH PROSTACYCLIN RECEPTOR_HOMO SAPIENS_R-HSA-392851 | 1.704 | 0.017 | 0.062 |
| REACTOME | TRANSLESION SYNTHESIS BY REV1_HOMO SAPIENS_R-HSA-110312 | 1.699 | 0.011 | 0.063 |
| REACTOME | CDT1 ASSOCIATION WITH THE CDC6:ORC:ORIGIN COMPLEX_HOMO SAPIENS_R-HSA-68827 | 1.706 | 0.025 | 0.063 |
| REACTOME | TRNA PROCESSING_HOMO SAPIENS_R-HSA-72306 | 1.698 | 0.026 | 0.063 |
| REACTOME | MITOCHONDRIAL TRANSLATION INITIATION_HOMO SAPIENS_R-HSA-5368286 | 1.704 | 0.028 | 0.063 |
| REACTOME | DEGRADATION OF AXIN_HOMO SAPIENS_R-HSA-4641257 | 1.701 | 0.032 | 0.063 |
| REACTOME | UBIQUITIN-DEPENDENT DEGRADATION OF CYCLIN D_HOMO SAPIENS_R-HSA-75815 | 1.699 | 0.025 | 0.063 |
| REACTOME | RNA POLYMERASE II HIV PROMOTER ESCAPE_HOMO SAPIENS_R-HSA-167162 | 1.723 | 0.019 | 0.063 |
| REACTOME | SUMOYLATION OF RNA BINDING PROTEINS_HOMO SAPIENS_R-HSA-4570464 | 1.698 | 0.026 | 0.063 |
| REACTOME | ACTIVATION OF THE PRE-REPLICATIVE COMPLEX_HOMO SAPIENS_R-HSA-68962 | 1.711 | 0.009 | 0.063 |
| REACTOME | PCP/CE PATHWAY_HOMO SAPIENS_R-HSA-4086400 | 1.697 | 0.026 | 0.063 |
| REACTOME | DEGRADATION OF GLI2 BY THE PROTEASOME_HOMO SAPIENS_R-HSA-5610783 | 1.706 | 0.026 | 0.063 |
| REACTOME | TRANSCRIPTION OF THE HIV GENOME_HOMO SAPIENS_R-HSA-167172 | 1.710 | 0.013 | 0.063 |
| REACTOME | UBIQUITIN MEDIATED DEGRADATION OF PHOSPHORYLATED CDC25A_HOMO SAPIENS_R-HSA-69601 | 1.711 | 0.028 | 0.063 |
| REACTOME | UBIQUITIN-DEPENDENT DEGRADATION OF CYCLIN D1_HOMO SAPIENS_R-HSA-69229 | 1.699 | 0.025 | 0.063 |
| REACTOME | RNA POLYMERASE II PRE-TRANSCRIPTION EVENTS_HOMO SAPIENS_R-HSA-674695 | 1.706 | 0.020 | 0.063 |
| REACTOME | ANCHORING OF THE BASAL BODY TO THE PLASMA MEMBRANE_HOMO SAPIENS_R-HSA-5620912 | 1.707 | 0.023 | 0.063 |
| REACTOME | P53-INDEPENDENT G1/S DNA DAMAGE CHECKPOINT_HOMO SAPIENS_R-HSA-69613 | 1.711 | 0.028 | 0.063 |
| REACTOME | P53-INDEPENDENT DNA DAMAGE RESPONSE_HOMO SAPIENS_R-HSA-69610 | 1.711 | 0.028 | 0.064 |
| REACTOME | RNA POLYMERASE I, RNA POLYMERASE III, AND MITOCHONDRIAL TRANSCRIPTION_HOMO SAPIENS_R-HSA-504046 | 1.694 | 0.034 | 0.064 |
| REACTOME | TRANSLESION SYNTHESIS BY POLK_HOMO SAPIENS_R-HSA-5655862 | 1.692 | 0.009 | 0.065 |
| REACTOME | STABILIZATION OF P53_HOMO SAPIENS_R-HSA-69541 | 1.690 | 0.033 | 0.066 |

| | | | | |
|---|---|---|---|---|
| REACTOME | TERMINATION OF TRANSLESION DNA SYNTHESIS_HOMO SAPIENS_R-HSA-5656169 | 1.687 | 0.018 | 0.066 |
| REACTOME | GLI3 IS PROCESSED TO GLI3R BY THE PROTEASOME_HOMO SAPIENS_R-HSA-5610785 | 1.685 | 0.036 | 0.067 |
| REACTOME | SIGNALING BY FGFR2 IN DISEASE_HOMO SAPIENS_R-HSA-5655253 | 1.681 | 0.000 | 0.069 |
| REACTOME | REGULATION OF APOPTOSIS_HOMO SAPIENS_R-HSA-169911 | 1.678 | 0.031 | 0.069 |
| REACTOME | RNA POLYMERASE II TRANSCRIBES SNRNA GENES_HOMO SAPIENS_R-HSA-6807505 | 1.678 | 0.014 | 0.069 |
| REACTOME | CDK-MEDIATED PHOSPHORYLATION AND REMOVAL OF CDC6_HOMO SAPIENS_R-HSA-69017 | 1.676 | 0.040 | 0.070 |
| REACTOME | TRANSLESION SYNTHESIS BY POLH_HOMO SAPIENS_R-HSA-110320 | 1.678 | 0.022 | 0.070 |
| REACTOME | SUMOYLATION_HOMO SAPIENS_R-HSA-2990846 | 1.675 | 0.036 | 0.070 |
| REACTOME | REGULATION OF ACTIVATED PAK-2P34 BY PROTEASOME MEDIATED DEGRADATION_HOMO SAPIENS_R-HSA-211733 | 1.669 | 0.036 | 0.072 |
| REACTOME | REGULATION OF ORNITHINE DECARBOXYLASE (ODC)_HOMO SAPIENS_R-HSA-350562 | 1.669 | 0.044 | 0.072 |
| REACTOME | AUF1 (HNRNP D0) BINDS AND DESTABILIZES MRNA_HOMO SAPIENS_R-HSA-450408 | 1.666 | 0.042 | 0.073 |
| REACTOME | SUMO E3 LIGASES SUMOYLATE TARGET PROTEINS_HOMO SAPIENS_R-HSA-3108232 | 1.657 | 0.045 | 0.075 |
| REACTOME | RESPIRATORY ELECTRON TRANSPORT, ATP SYNTHESIS BY CHEMIOSMOTIC COUPLING, AND HEAT PRODUCTION BY UNCOUPLING PROTEINS._HOMO SAPIENS_R-HSA-163200 | 1.659 | 0.045 | 0.075 |
| REACTOME | SNRNP ASSEMBLY_HOMO SAPIENS_R-HSA-191859 | 1.658 | 0.037 | 0.075 |
| REACTOME | METABOLISM OF NON-CODING RNA_HOMO SAPIENS_R-HSA-194441 | 1.658 | 0.037 | 0.076 |
| REACTOME | KINESINS_HOMO SAPIENS_R-HSA-983189 | 1.651 | 0.041 | 0.078 |
| REACTOME | NUCLEAR ENVELOPE BREAKDOWN_HOMO SAPIENS_R-HSA-2980766 | 1.648 | 0.040 | 0.080 |
| REACTOME | FGFR2 MUTANT RECEPTOR ACTIVATION_HOMO SAPIENS_R-HSA-1839126 | 1.641 | 0.019 | 0.082 |
| REACTOME | COMPLEX I BIOGENESIS_HOMO SAPIENS_R-HSA-6799198 | 1.639 | 0.048 | 0.083 |
| REACTOME | POST-ELONGATION PROCESSING OF THE TRANSCRIPT_HOMO SAPIENS_R-HSA-76044 | 1.632 | 0.049 | 0.085 |
| REACTOME | RNA POLYMERASE II TRANSCRIPTION TERMINATION_HOMO SAPIENS_R-HSA-73856 | 1.632 | 0.049 | 0.086 |
| REACTOME | CLEAVAGE OF GROWING TRANSCRIPT IN THE TERMINATION REGION_HOMO SAPIENS_R-HSA-109688 | 1.632 | 0.049 | 0.086 |
| REACTOME | DEGRADATION OF GLI1 BY THE PROTEASOME_HOMO SAPIENS_R-HSA-5610780 | 1.627 | 0.045 | 0.087 |
| REACTOME | RAF ACTIVATION_HOMO SAPIENS_R-HSA-5673000 | 1.626 | 0.020 | 0.087 |
| REACTOME | MAPK6/MAPK4 SIGNALING_HOMO SAPIENS_R-HSA-5687128 | 1.617 | 0.046 | 0.091 |
| REACTOME | TCF DEPENDENT SIGNALING IN RESPONSE TO WNT_HOMO SAPIENS_R-HSA-201681 | 1.618 | 0.016 | 0.091 |
| REACTOME | VPR-MEDIATED NUCLEAR IMPORT OF PICS_HOMO SAPIENS_R-HSA-180910 | 1.609 | 0.044 | 0.095 |
| REACTOME | RNA POLYMERASE I TRANSCRIPTION INITIATION_HOMO SAPIENS_R-HSA-73762 | 1.594 | 0.034 | 0.102 |
| REACTOME | CONSTITUTIVE SIGNALING BY AKT1 E17K IN CANCER_HOMO SAPIENS_R-HSA-5674400 | 1.590 | 0.034 | 0.103 |
| REACTOME | SIGNALING BY RHO GTPASES_HOMO SAPIENS_R-HSA-194315 | 1.589 | 0.024 | 0.104 |
| REACTOME | REGULATION OF TP53 ACTIVITY THROUGH METHYLATION_HOMO SAPIENS_R-HSA-6804760 | 1.587 | 0.030 | 0.104 |
| REACTOME | PROCESSING OF CAPPED INTRONLESS PRE-MRNA_HOMO SAPIENS_R-HSA-75067 | 1.577 | 0.042 | 0.109 |
| REACTOME | POST-ELONGATION PROCESSING OF INTRONLESS PRE-MRNA_HOMO SAPIENS_R-HSA-112297 | 1.577 | 0.042 | 0.109 |
| REACTOME | PYRIMIDINE METABOLISM_HOMO SAPIENS_R-HSA-73848 | 1.571 | 0.019 | 0.111 |
| REACTOME | CELLULAR RESPONSES TO STRESS_HOMO SAPIENS_R-HSA-2262752 | 1.566 | 0.015 | 0.113 |
| REACTOME | NEGATIVE REGULATION OF MAPK PATHWAY_HOMO SAPIENS_R-HSA-5675221 | 1.563 | 0.050 | 0.114 |

| | | | | |
|---|---|---|---|---|
| REACTOME | ANTIGEN PROCESSING: UBIQUITINATION & PROTEASOME DEGRADATION_HOMO SAPIENS_R-HSA-983168 | 1.559 | 0.039 | 0.115 |
| REACTOME | GOLGI-TO-ER RETROGRADE TRANSPORT_HOMO SAPIENS_R-HSA-8856688 | 1.550 | 0.031 | 0.118 |
| REACTOME | BETA-CATENIN INDEPENDENT WNT SIGNALING_HOMO SAPIENS_R-HSA-3858494 | 1.545 | 0.039 | 0.119 |
| REACTOME | SIGNALING BY WNT_HOMO SAPIENS_R-HSA-195721 | 1.516 | 0.023 | 0.130 |
| REACTOME | DOWNSTREAM SIGNALING EVENTS OF B CELL RECEPTOR (BCR)_HOMO SAPIENS_R-HSA-1168372 | 1.412 | 0.047 | 0.188 |
| REACTOME | DISEASES OF SIGNAL TRANSDUCTION_HOMO SAPIENS_R-HSA-5663202 | 1.341 | 0.045 | 0.240 |
| REACTOME | FCERI MEDIATED CA+2 MOBILIZATION_HOMO SAPIENS_R-HSA-2871809 | -2.092 | 0.000 | 0.036 |
| REACTOME | SYNTHESIS OF IP3 AND IP4 IN THE CYTOSOL_HOMO SAPIENS_R-HSA-1855204 | -2.020 | 0.000 | 0.058 |
| REACTOME | SYNTHESIS OF PIPS AT THE PLASMA MEMBRANE_HOMO SAPIENS_R-HSA-1660499 | -1.942 | 0.000 | 0.144 |
| REACTOME | TRANSPORT OF GLUCOSE AND OTHER SUGARS, BILE SALTS AND ORGANIC ACIDS, METAL IONS AND AMINE COMPOUNDS_HOMO SAPIENS_R-HSA-425366 | -1.751 | 0.000 | 0.204 |
| REACTOME | DISEASES ASSOCIATED WITH THE TLR SIGNALING CASCADE_HOMO SAPIENS_R-HSA-5602358 | -1.873 | 0.002 | 0.186 |
| REACTOME | DISEASES OF IMMUNE SYSTEM_HOMO SAPIENS_R-HSA-5260271 | -1.873 | 0.002 | 0.159 |
| REACTOME | INTERLEUKIN-6 FAMILY SIGNALING_HOMO SAPIENS_R-HSA-6783589 | -1.808 | 0.002 | 0.141 |
| REACTOME | ANTIGEN ACTIVATES B CELL RECEPTOR (BCR) LEADING TO GENERATION OF SECOND MESSENGERS_HOMO SAPIENS_R-HSA-983695 | -1.921 | 0.002 | 0.152 |
| REACTOME | NUCLEAR SIGNALING BY ERBB4_HOMO SAPIENS_R-HSA-1251985 | -1.759 | 0.002 | 0.199 |
| REACTOME | METAL ION SLC TRANSPORTERS_HOMO SAPIENS_R-HSA-425410 | -1.842 | 0.002 | 0.180 |
| REACTOME | GROWTH HORMONE RECEPTOR SIGNALING_HOMO SAPIENS_R-HSA-982772 | -1.825 | 0.004 | 0.151 |
| REACTOME | ROLE OF PHOSPHOLIPIDS IN PHAGOCYTOSIS_HOMO SAPIENS_R-HSA-2029485 | -1.809 | 0.004 | 0.149 |
| REACTOME | IL-6-TYPE CYTOKINE RECEPTOR LIGAND INTERACTIONS_HOMO SAPIENS_R-HSA-6788467 | -1.866 | 0.004 | 0.151 |
| REACTOME | SPHINGOLIPID METABOLISM_HOMO SAPIENS_R-HSA-428157 | -1.815 | 0.005 | 0.157 |
| REACTOME | PLATELET AGGREGATION (PLUG FORMATION)_HOMO SAPIENS_R-HSA-76009 | -1.837 | 0.005 | 0.173 |
| REACTOME | NUCLEOTIDE-BINDING DOMAIN, LEUCINE RICH REPEAT CONTAINING RECEPTOR (NLR) SIGNALING PATHWAYS_HOMO SAPIENS_R-HSA-168643 | -1.830 | 0.007 | 0.155 |
| REACTOME | PHOSPHOLIPID METABOLISM_HOMO SAPIENS_R-HSA-1483257 | -1.582 | 0.008 | 0.238 |
| REACTOME | CELL SURFACE INTERACTIONS AT THE VASCULAR WALL_HOMO SAPIENS_R-HSA-202733 | -1.832 | 0.009 | 0.167 |
| REACTOME | TOLL-LIKE RECEPTORS CASCADES_HOMO SAPIENS_R-HSA-168898 | -1.725 | 0.009 | 0.201 |
| REACTOME | CASPASE ACTIVATION VIA EXTRINSIC APOPTOTIC SIGNALIG PATHWAY_HOMO SAPIENS_R-HSA-5357769 | -1.887 | 0.011 | 0.189 |
| REACTOME | INOSITOL PHOSPHATE METABOLISM_HOMO SAPIENS_R-HSA-1483249 | -1.687 | 0.011 | 0.245 |
| REACTOME | NOD1/2 SIGNALING PATHWAY_HOMO SAPIENS_R-HSA-168638 | -1.811 | 0.011 | 0.155 |
| REACTOME | ACTIVATED TLR4 SIGNALLING_HOMO SAPIENS_R-HSA-166054 | -1.648 | 0.012 | 0.205 |
| REACTOME | LIGAND-DEPENDENT CASPASE ACTIVATION_HOMO SAPIENS_R-HSA-140534 | -1.735 | 0.012 | 0.198 |
| REACTOME | MOLECULES ASSOCIATED WITH ELASTIC FIBRES_HOMO SAPIENS_R-HSA-2129379 | -1.744 | 0.014 | 0.208 |
| REACTOME | MYD88-INDEPENDENT TLR3/TLR4 CASCADE_HOMO SAPIENS_R-HSA-166166 | -1.673 | 0.014 | 0.232 |
| REACTOME | TOLL LIKE RECEPTOR 3 (TLR3) CASCADE_HOMO SAPIENS_R-HSA-168164 | -1.673 | 0.014 | 0.226 |
| REACTOME | TRIF-MEDIATED TLR3/TLR4 SIGNALING_HOMO SAPIENS_R-HSA-937061 | -1.673 | 0.014 | 0.220 |
| REACTOME | PI METABOLISM_HOMO SAPIENS_R-HSA-1483255 | -1.682 | 0.015 | 0.231 |

| | | | | |
|---|---|---|---|---|
| REACTOME | O-LINKED GLYCOSYLATION OF MUCINS_HOMO SAPIENS_R-HSA-913709 | -1.629 | 0.015 | 0.216 |
| REACTOME | GPVI-MEDIATED ACTIVATION CASCADE_HOMO SAPIENS_R-HSA-114604 | -1.723 | 0.015 | 0.198 |
| REACTOME | INTEGRIN ALPHAIIB BETA3 SIGNALING_HOMO SAPIENS_R-HSA-354192 | -1.765 | 0.016 | 0.199 |
| REACTOME | COMPLEMENT CASCADE_HOMO SAPIENS_R-HSA-166658 | -1.783 | 0.016 | 0.174 |
| REACTOME | TOLL LIKE RECEPTOR 4 (TLR4) CASCADE_HOMO SAPIENS_R-HSA-166016 | -1.669 | 0.016 | 0.205 |
| REACTOME | SPHINGOLIPID DE NOVO BIOSYNTHESIS_HOMO SAPIENS_R-HSA-1660661 | -1.675 | 0.017 | 0.235 |
| REACTOME | PLATELET ACTIVATION, SIGNALING AND AGGREGATION_HOMO SAPIENS_R-HSA-76002 | -1.727 | 0.017 | 0.204 |
| REACTOME | ZBP1(DAI) MEDIATED INDUCTION OF TYPE I IFNS_HOMO SAPIENS_R-HSA-1606322 | -1.738 | 0.018 | 0.201 |
| REACTOME | DEATH RECEPTOR SIGNALLING_HOMO SAPIENS_R-HSA-73887 | -1.635 | 0.019 | 0.223 |
| REACTOME | EGFR INTERACTS WITH PHOSPHOLIPASE C-GAMMA_HOMO SAPIENS_R-HSA-212718 | -1.659 | 0.019 | 0.202 |
| REACTOME | TRANS-GOLGI NETWORK VESICLE BUDDING_HOMO SAPIENS_R-HSA-199992 | -1.579 | 0.021 | 0.240 |
| REACTOME | CLATHRIN DERIVED VESICLE BUDDING_HOMO SAPIENS_R-HSA-421837 | -1.579 | 0.021 | 0.236 |
| REACTOME | TNFR1-INDUCED NFKAPPAB SIGNALING PATHWAY_HOMO SAPIENS_R-HSA-5357956 | -1.670 | 0.021 | 0.214 |
| REACTOME | PHOSPHOLIPASE C-MEDIATED CASCADE; FGFR4_HOMO SAPIENS_R-HSA-5654228 | -1.664 | 0.022 | 0.199 |
| REACTOME | ABC-FAMILY PROTEINS MEDIATED TRANSPORT_HOMO SAPIENS_R-HSA-382556 | -1.572 | 0.022 | 0.244 |
| REACTOME | GLYCOSPHINGOLIPID METABOLISM_HOMO SAPIENS_R-HSA-1660662 | -1.670 | 0.023 | 0.209 |
| REACTOME | PHOSPHOLIPASE C-MEDIATED CASCADE; FGFR3_HOMO SAPIENS_R-HSA-5654227 | -1.631 | 0.024 | 0.217 |
| REACTOME | PLC BETA MEDIATED EVENTS_HOMO SAPIENS_R-HSA-112043 | -1.613 | 0.024 | 0.222 |
| REACTOME | BMAL1:CLOCK,NPAS2 ACTIVATES CIRCADIAN GENE EXPRESSION_HOMO SAPIENS_R-HSA-1368108 | -1.684 | 0.024 | 0.234 |
| REACTOME | FGFR1 MUTANT RECEPTOR ACTIVATION_HOMO SAPIENS_R-HSA-1839124 | -1.591 | 0.025 | 0.243 |
| REACTOME | DAG AND IP3 SIGNALING_HOMO SAPIENS_R-HSA-1489509 | -1.627 | 0.025 | 0.214 |
| REACTOME | G-PROTEIN MEDIATED EVENTS_HOMO SAPIENS_R-HSA-112040 | -1.615 | 0.026 | 0.230 |
| REACTOME | REGULATION OF COMPLEMENT CASCADE_HOMO SAPIENS_R-HSA-977606 | -1.715 | 0.027 | 0.202 |
| REACTOME | ACTIVATED TAK1 MEDIATES P38 MAPK ACTIVATION_HOMO SAPIENS_R-HSA-450302 | -1.587 | 0.028 | 0.242 |
| REACTOME | PLC-GAMMA1 SIGNALLING_HOMO SAPIENS_R-HSA-167021 | -1.614 | 0.029 | 0.227 |
| REACTOME | PHOSPHOLIPASE C-MEDIATED CASCADE: FGFR1_HOMO SAPIENS_R-HSA-5654219 | -1.666 | 0.030 | 0.200 |
| REACTOME | SYNTHESIS OF LEUKOTRIENES (LT) AND EOXINS (EX)_HOMO SAPIENS_R-HSA-2142691 | -1.667 | 0.031 | 0.202 |
| REACTOME | INTERFERON GAMMA SIGNALING_HOMO SAPIENS_R-HSA-877300 | -1.743 | 0.032 | 0.201 |
| REACTOME | TNFS BIND THEIR PHYSIOLOGICAL RECEPTORS_HOMO SAPIENS_R-HSA-5669034 | -1.685 | 0.035 | 0.240 |
| REACTOME | REGULATED NECROSIS_HOMO SAPIENS_R-HSA-5218859 | -1.654 | 0.037 | 0.205 |
| REACTOME | RIPK1-MEDIATED REGULATED NECROSIS_HOMO SAPIENS_R-HSA-5213460 | -1.654 | 0.037 | 0.201 |
| REACTOME | INITIAL TRIGGERING OF COMPLEMENT_HOMO SAPIENS_R-HSA-166663 | -1.635 | 0.037 | 0.219 |
| REACTOME | TNF RECEPTOR SUPERFAMILY (TNFSF) MEMBERS MEDIATING NON-CANONICAL NF-KB PATHWAY_HOMO SAPIENS_R-HSA-5676594 | -1.593 | 0.038 | 0.246 |
| REACTOME | BILE ACID AND BILE SALT METABOLISM_HOMO SAPIENS_R-HSA-194068 | -1.585 | 0.038 | 0.238 |
| REACTOME | REGULATION OF KIT SIGNALING_HOMO SAPIENS_R-HSA-1433559 | -1.585 | 0.040 | 0.241 |
| REACTOME | SYNDECAN INTERACTIONS_HOMO SAPIENS_R-HSA-3000170 | -1.671 | 0.041 | 0.218 |
| REACTOME | FCGAMMA RECEPTOR (FCGR) DEPENDENT PHAGOCYTOSIS_HOMO SAPIENS_R-HSA-2029480 | -1.633 | 0.041 | 0.216 |

| | | | | |
|---|---|---|---|---|
| REACTOME | CHEMOKINE RECEPTORS BIND CHEMOKINES_HOMO SAPIENS_R-HSA-380108 | -1.613 | 0.044 | 0.225 |
| REACTOME | PHOSPHOLIPASE C-MEDIATED CASCADE; FGFR2_HOMO SAPIENS_R-HSA-5654221 | -1.587 | 0.046 | 0.245 |

Abbreviations: NES, normalized enrichment score; NOM, nominal; FDR, false discovery rate.

**Supplementary Table 2.** *PIMREG* enriched gene sets related to DDR in GBM cohort (TCGA-PanCancer).

| Database | Gene Set | NES | NOM *p*-value | FDR q-value |
|---|---|---|---|---|
| HALLMARK | DNA_REPAIR | 2.015 | 0.000 | 0.004 |
| HALLMARK | UV_RESPONSE_DN | -1.883 | 0.000 | 0.122 |
| KEGG | BASE EXCISION REPAIR | 1.849 | 0.004 | 0.067 |
| KEGG | DNA REPLICATION | 1.908 | 0.004 | 0.074 |
| KEGG | FANCONI ANEMIA PATHWAY | 1.821 | 0.005 | 0.076 |
| KEGG | NUCLEOTIDE EXCISION REPAIR | 1.797 | 0.014 | 0.085 |
| KEGG | MISMATCH REPAIR | 1.691 | 0.030 | 0.140 |
| REACTOME | DNA REPLICATION_HOMO SAPIENS_R-HSA-69306 | 2.058 | 0.000 | 0.017 |
| REACTOME | TRANSCRIPTION-COUPLED NUCLEOTIDE EXCISION REPAIR (TC-NER)_HOMO SAPIENS_R-HSA-6781827 | 2.037 | 0.000 | 0.019 |
| REACTOME | DUAL INCISION IN TC-NER_HOMO SAPIENS_R-HSA-6782135 | 1.965 | 0.002 | 0.020 |
| REACTOME | GAP-FILLING DNA REPAIR SYNTHESIS AND LIGATION IN TC-NER_HOMO SAPIENS_R-HSA-6782210 | 1.998 | 0.002 | 0.021 |
| REACTOME | NUCLEOTIDE EXCISION REPAIR_HOMO SAPIENS_R-HSA-5696398 | 1.975 | 0.004 | 0.021 |
| REACTOME | DNA DOUBLE-STRAND BREAK REPAIR_HOMO SAPIENS_R-HSA-5693532 | 1.938 | 0.005 | 0.024 |
| REACTOME | RESOLUTION OF ABASIC SITES (AP SITES)_HOMO SAPIENS_R-HSA-73933 | 1.902 | 0.000 | 0.027 |
| REACTOME | BASE EXCISION REPAIR_HOMO SAPIENS_R-HSA-73884 | 1.902 | 0.000 | 0.028 |
| REACTOME | HOMOLOGY DIRECTED REPAIR_HOMO SAPIENS_R-HSA-5693538 | 1.899 | 0.007 | 0.028 |
| REACTOME | HDR THROUGH HOMOLOGOUS RECOMBINATION (HR) OR SINGLE STRAND ANNEALING (SSA)_HOMO SAPIENS_R-HSA-5693567 | 1.885 | 0.011 | 0.028 |
| REACTOME | GLOBAL GENOME NUCLEOTIDE EXCISION REPAIR (GG-NER)_HOMO SAPIENS_R-HSA-5696399 | 1.916 | 0.002 | 0.028 |
| REACTOME | FANCONI ANEMIA PATHWAY_HOMO SAPIENS_R-HSA-6783310 | 1.892 | 0.002 | 0.029 |
| REACTOME | FORMATION OF TC-NER PRE-INCISION COMPLEX_HOMO SAPIENS_R-HSA-6781823 | 1.871 | 0.006 | 0.030 |
| REACTOME | DNA DAMAGE RECOGNITION IN GG-NER_HOMO SAPIENS_R-HSA-5696394 | 1.836 | 0.009 | 0.035 |
| REACTOME | HDR THROUGH HOMOLOGOUS RECOMBINATION (HRR)_HOMO SAPIENS_R-HSA-5685942 | 1.823 | 0.018 | 0.038 |
| REACTOME | RESOLUTION OF AP SITES VIA THE MULTIPLE-NUCLEOTIDE PATCH REPLACEMENT PATHWAY_HOMO SAPIENS_R-HSA-110373 | 1.810 | 0.002 | 0.042 |
| REACTOME | DNA DAMAGE BYPASS_HOMO SAPIENS_R-HSA-73893 | 1.811 | 0.005 | 0.042 |
| REACTOME | P53-DEPENDENT G1 DNA DAMAGE RESPONSE_HOMO SAPIENS_R-HSA-69563 | 1.804 | 0.017 | 0.042 |
| REACTOME | P53-DEPENDENT G1/S DNA DAMAGE CHECKPOINT_HOMO SAPIENS_R-HSA-69580 | 1.804 | 0.017 | 0.042 |
| REACTOME | GAP-FILLING DNA REPAIR SYNTHESIS AND LIGATION IN GG-NER_HOMO SAPIENS_R-HSA-5696397 | 1.797 | 0.006 | 0.044 |
| REACTOME | ACTIVATION OF ATR IN RESPONSE TO REPLICATION STRESS_HOMO SAPIENS_R-HSA-176187 | 1.793 | 0.009 | 0.045 |
| REACTOME | DUAL INCISION IN GG-NER_HOMO SAPIENS_R-HSA-5696400 | 1.781 | 0.002 | 0.048 |
| REACTOME | PROCESSING OF DNA DOUBLE-STRAND BREAK ENDS_HOMO SAPIENS_R-HSA-5693607 | 1.765 | 0.027 | 0.051 |
| REACTOME | PCNA-DEPENDENT LONG PATCH BASE EXCISION REPAIR_HOMO SAPIENS_R-HSA-5651801 | 1.750 | 0.005 | 0.056 |
| REACTOME | G2/M DNA DAMAGE CHECKPOINT_HOMO SAPIENS_R-HSA-69473 | 1.741 | 0.037 | 0.058 |
| REACTOME | TRANSLESION SYNTHESIS BY POLI_HOMO SAPIENS_R-HSA-5656121 | 1.733 | 0.004 | 0.061 |

Abbreviations: NES, normalized enrichment score; NOM, nominal; FDR, false discovery rate.

**Supplementary Table 3.** *PIMREG* enriched gene sets related to RNA metabolism in GBM cohort (TCGA-PanCancer).

| Database | Gene Set | NES | NOM *p*-value | FDR q-value |
|---|---|---|---|---|
| KEGG | SPLICEOSOME | 1.915 | 0.002 | 0.136 |
| KEGG | RNA POLYMERASE | 1.875 | 0.002 | 0.071 |
| KEGG | RNA TRANSPORT | 1.687 | 0.028 | 0.126 |
| KEGG | RNA DEGRADATION | 1.566 | 0.042 | 0.216 |
| KEGG | RIBOSOME | 1.684 | 0.046 | 0.120 |
| KEGG | BASAL TRANSCRIPTION FACTORS | 1.557 | 0.049 | 0.217 |
| REACTOME | RNA POLYMERASE III TRANSCRIPTION INITIATION_HOMO SAPIENS_R-HSA-76046 | 2.092 | 0.000 | 0.014 |
| REACTOME | PROCESSING OF CAPPED INTRON-CONTAINING PRE-MRNA_HOMO SAPIENS_R-HSA-72203 | 1.961 | 0.002 | 0.020 |
| REACTOME | MRNA SPLICING_HOMO SAPIENS_R-HSA-72172 | 2.005 | 0.000 | 0.020 |
| REACTOME | RNA POLYMERASE III TRANSCRIPTION INITIATION FROM TYPE 1 PROMOTER_HOMO SAPIENS_R-HSA-76061 | 2.024 | 0.000 | 0.020 |
| REACTOME | RNA POLYMERASE III TRANSCRIPTION_HOMO SAPIENS_R-HSA-74158 | 1.966 | 0.000 | 0.020 |
| REACTOME | RNA POLYMERASE III ABORTIVE AND RETRACTIVE INITIATION_HOMO SAPIENS_R-HSA-749476 | 1.966 | 0.000 | 0.021 |
| REACTOME | MRNA SPLICING - MINOR PATHWAY_HOMO SAPIENS_R-HSA-72165 | 2.008 | 0.000 | 0.021 |
| REACTOME | FGFR2 ALTERNATIVE SPLICING_HOMO SAPIENS_R-HSA-6803529 | 1.992 | 0.000 | 0.021 |
| REACTOME | RNA POLYMERASE III TRANSCRIPTION INITIATION FROM TYPE 2 PROMOTER_HOMO SAPIENS_R-HSA-76066 | 1.993 | 0.000 | 0.021 |
| REACTOME | RNA POLYMERASE III TRANSCRIPTION INITIATION FROM TYPE 3 PROMOTER_HOMO SAPIENS_R-HSA-76071 | 1.986 | 0.000 | 0.021 |
| REACTOME | MRNA SPLICING - MAJOR PATHWAY_HOMO SAPIENS_R-HSA-72163 | 1.976 | 0.000 | 0.022 |
| REACTOME | FORMATION OF RNA POL II ELONGATION COMPLEX_HOMO SAPIENS_R-HSA-112382 | 1.878 | 0.004 | 0.029 |
| REACTOME | RNA POLYMERASE II TRANSCRIPTION ELONGATION_HOMO SAPIENS_R-HSA-75955 | 1.878 | 0.004 | 0.029 |
| REACTOME | MICRORNA (MIRNA) BIOGENESIS_HOMO SAPIENS_R-HSA-203927 | 1.866 | 0.002 | 0.031 |
| REACTOME | MRNA CAPPING_HOMO SAPIENS_R-HSA-72086 | 1.836 | 0.006 | 0.036 |
| REACTOME | RNA POL II CTD PHOSPHORYLATION AND INTERACTION WITH CE_HOMO SAPIENS_R-HSA-77075 | 1.836 | 0.006 | 0.037 |
| REACTOME | RNA POLYMERASE III TRANSCRIPTION TERMINATION_HOMO SAPIENS_R-HSA-73980 | 1.828 | 0.004 | 0.037 |
| REACTOME | RNA POL II CTD PHOSPHORYLATION AND INTERACTION WITH CE_HOMO SAPIENS_R-HSA-167160 | 1.836 | 0.006 | 0.037 |
| REACTOME | RNA POLYMERASE III CHAIN ELONGATION_HOMO SAPIENS_R-HSA-73780 | 1.837 | 0.000 | 0.037 |
| REACTOME | PIWI-INTERACTING RNA (PIRNA) BIOGENESIS_HOMO SAPIENS_R-HSA-5601884 | 1.811 | 0.011 | 0.041 |
| REACTOME | RNA POLYMERASE II TRANSCRIPTION_HOMO SAPIENS_R-HSA-73857 | 1.771 | 0.015 | 0.050 |
| REACTOME | DEADENYLATION-DEPENDENT MRNA DECAY_HOMO SAPIENS_R-HSA-429914 | 1.758 | 0.009 | 0.053 |
| REACTOME | GENE SILENCING BY RNA_HOMO SAPIENS_R-HSA-211000 | 1.727 | 0.036 | 0.061 |
| REACTOME | MITOTIC PROPHASE_HOMO SAPIENS_R-HSA-68875 | 1.731 | 0.034 | 0.061 |
| REACTOME | TRNA PROCESSING IN THE NUCLEUS_HOMO SAPIENS_R-HSA-6784531 | 1.717 | 0.026 | 0.061 |
| REACTOME | RNA POLYMERASE II TRANSCRIPTION PRE-INITIATION AND PROMOTER OPENING_HOMO SAPIENS_R-HSA-73779 | 1.723 | 0.019 | 0.061 |
| REACTOME | REGULATION OF MRNA STABILITY BY PROTEINS THAT BIND AU-RICH ELEMENTS_HOMO SAPIENS_R-HSA-450531 | 1.718 | 0.036 | 0.061 |

| | | | | |
|---|---|---|---|---|
| REACTOME | RNA POLYMERASE II TRANSCRIPTION INITIATION_HOMO SAPIENS_R-HSA-75953 | 1.723 | 0.019 | 0.062 |
| REACTOME | RNA POLYMERASE II TRANSCRIPTION INITIATION AND PROMOTER CLEARANCE_HOMO SAPIENS_R-HSA-76042 | 1.723 | 0.019 | 0.062 |
| REACTOME | RNA POLYMERASE II PROMOTER ESCAPE_HOMO SAPIENS_R-HSA-73776 | 1.723 | 0.019 | 0.062 |
| REACTOME | TRNA PROCESSING_HOMO SAPIENS_R-HSA-72306 | 1.698 | 0.026 | 0.063 |
| REACTOME | RNA POLYMERASE II PRE-TRANSCRIPTION EVENTS_HOMO SAPIENS_R-HSA-674695 | 1.706 | 0.020 | 0.063 |
| REACTOME | RNA POLYMERASE I, RNA POLYMERASE III, AND MITOCHONDRIAL TRANSCRIPTION_HOMO SAPIENS_R-HSA-504046 | 1.694 | 0.034 | 0.064 |
| REACTOME | RNA POLYMERASE II TRANSCRIBES SNRNA GENES_HOMO SAPIENS_R-HSA-6807505 | 1.678 | 0.014 | 0.069 |
| REACTOME | AUF1 (HNRNP D0) BINDS AND DESTABILIZES MRNA_HOMO SAPIENS_R-HSA-450408 | 1.666 | 0.042 | 0.073 |
| REACTOME | SNRNP ASSEMBLY_HOMO SAPIENS_R-HSA-191859 | 1.658 | 0.037 | 0.075 |
| REACTOME | METABOLISM OF NON-CODING RNA_HOMO SAPIENS_R-HSA-194441 | 1.658 | 0.037 | 0.076 |
| REACTOME | POST-ELONGATION PROCESSING OF THE TRANSCRIPT_HOMO SAPIENS_R-HSA-76044 | 1.632 | 0.049 | 0.085 |
| REACTOME | RNA POLYMERASE II TRANSCRIPTION TERMINATION_HOMO SAPIENS_R-HSA-73856 | 1.632 | 0.049 | 0.086 |
| REACTOME | CLEAVAGE OF GROWING TRANSCRIPT IN THE TERMINATION REGION_HOMO SAPIENS_R-HSA-109688 | 1.632 | 0.049 | 0.086 |
| REACTOME | RNA POLYMERASE I TRANSCRIPTION INITIATION_HOMO SAPIENS_R-HSA-73762 | 1.594 | 0.034 | 0.102 |
| REACTOME | PROCESSING OF CAPPED INTRONLESS PRE-MRNA_HOMO SAPIENS_R-HSA-75067 | 1.577 | 0.042 | 0.109 |
| REACTOME | POST-ELONGATION PROCESSING OF INTRONLESS PRE-MRNA_HOMO SAPIENS_R-HSA-112297 | 1.577 | 0.042 | 0.109 |

Abbreviations: NES, normalized enrichment score; NOM, nominal; FDR, false discovery rate.

**Supplementary Table 4:** Proteins identified as PIMREG interactors in U87MG cells (PIMREG IP vs. IgG under vehicle-DMSO conditions)

| Gene name | log2 FC | Adj. *p*-value | -log10 *p*-value |
|---|---:|---:|---:|
| MYOF | 11.315854 | 0.000002 | 5.779101 |
| PPP1R12A | 6.802054 | 0.000008 | 5.102963 |
| SEP2 | 9.929833 | 0.000009 | 5.043518 |
| SRI | 12.958570 | 0.000030 | 4.522347 |
| IGLL5 | 15.723802 | 0.000034 | 4.473435 |
| RBMS1 | 13.013868 | 0.000034 | 4.465976 |
| SMNDC1 | 13.185869 | 0.000056 | 4.250398 |
| SNRPC | 15.639436 | 0.000062 | 4.205699 |
| EIF5A | 11.956166 | 0.000082 | 4.086294 |
| FKBP15 | 13.818202 | 0.000091 | 4.040672 |
| ZNF316 | 9.510583 | 0.000117 | 3.929979 |
| FAM32A | 13.184695 | 0.000118 | 3.929597 |
| TLE3 | 9.687383 | 0.000118 | 3.926491 |
| RIOK1 | 8.252126 | 0.000140 | 3.852464 |
| LTV1 | 10.407317 | 0.000191 | 3.719399 |
| WTAP | 9.574577 | 0.000205 | 3.688115 |
| AVEN | 11.020655 | 0.000270 | 3.568966 |
| CKAP4 | 11.561487 | 0.000554 | 3.256356 |
| SF1 | 15.504755 | 0.000627 | 3.202943 |
| CDC37 | 11.891359 | 0.000947 | 3.023551 |
| IGKVA18 | 3.367507 | 0.001571 | 2.803865 |
| PHF2 | 10.275753 | 0.002007 | 2.697540 |
| ADNP | 9.520758 | 0.002007 | 2.697540 |
| PPIA | 13.325497 | 0.002050 | 2.688351 |
| TSR1 | 10.370016 | 0.002050 | 2.688351 |
| SF3B2 | 11.478864 | 0.002071 | 2.683867 |
| CCAR1 | 8.868769 | 0.002168 | 2.663938 |
| RAB14 | 9.161145 | 0.002170 | 2.663541 |
| USP15 | 8.687307 | 0.003100 | 2.508587 |
| YME1L1 | 7.440926 | 0.003100 | 2.508587 |
| TCOF1 | 9.583739 | 0.003207 | 2.493854 |
| UACA | 6.064521 | 0.003900 | 2.408981 |
| TPI1 | 10.492472 | 0.005197 | 2.284238 |
| SNRNP70 | 1.191269 | 0.027388 | 1.562444 |
| EIF4G1 | 1.602117 | 0.030416 | 1.516903 |
| CHTF8 | 2.680105 | 0.040034 | 1.397572 |

**Supplementary Table 5:** Proteins identified as TMZ-regulated PIMREG interactors in U87MG cells (TMZ vs. DMSO - PIMREG IPs)

| Gene name | log2 FC | Adj. *p*-value | -log10 *p*-value |
|---|---|---|---|
| NAA40 | 9.3825 | 0.0002 | 3.6328 |
| ZW10 | 7.4520 | 0.0004 | 3.3487 |
| NDUFS6 | 11.7024 | 0.0004 | 3.3487 |
| FAM32A | -13.1847 | 0.0005 | 3.3295 |
| MTDH | -12.8130 | 0.0005 | 3.3295 |
| TLE3 | -9.6874 | 0.0005 | 3.3295 |
| PRMT5 | 9.5973 | 0.0005 | 3.3295 |
| NDUFA8 | 10.8102 | 0.0005 | 3.3295 |
| TIA1 | 12.1996 | 0.0005 | 3.3295 |
| STOM | 12.3241 | 0.0005 | 3.3295 |
| MIF | 13.5607 | 0.0005 | 3.3295 |
| NEDD4L | -9.3180 | 0.0005 | 3.2629 |
| MRPL15 | 11.3214 | 0.0010 | 3.0198 |
| LSM14B | 12.5895 | 0.0011 | 2.9736 |
| TDRD3 | -9.1960 | 0.0015 | 2.8258 |
| NCBP1 | 9.5267 | 0.0020 | 2.6947 |
| PRPF38B | -12.4638 | 0.0021 | 2.6761 |
| DARS2 | 8.3479 | 0.0022 | 2.6593 |
| MRPL44 | 9.7024 | 0.0022 | 2.6593 |
| UFM1 | 12.3424 | 0.0027 | 2.5716 |
| AIP | 10.7708 | 0.0027 | 2.5712 |
| RBM22 | 11.6678 | 0.0029 | 2.5678 |
| ELP3 | 9.9805 | 0.0083 | 2.5358 |
| VPS4B | 10.6899 | 0.0121 | 2.0807 |
| PAFAH1B3 | 10.0534 | 0.0124 | 1.9161 |
| SLC39A10 | -9.4655 | 0.0238 | 1.9059 |
| TRIP13 | 2.6948 | 0.0406 | 1.6229 |
| ACAA1 | 8.2234 | 0.0406 | 1.3912 |
| GNA13 | 9.1202 | 0.0027 | 1.3912 |

**Supplementary Table 6:** Proteins identified as PIMREG interactors in T98G cells (PIMREG IP vs. IgG under vehicle-DMSO conditions)

| **Gene name** | **log2 FC** | **Adj. *p*-value** | **-log10 *p*-value** |
|---|---:|---:|---:|
| IGLL5 | 14.7159 | 0.000303 | 3.518833145 |
| FKBP15 | 12.3106 | 0.002409 | 2.618132569 |
| PIMREG | 11.4388 | 0.007102 | 2.148628406 |
| CIR1 | 8.9144 | 0.001074 | 2.969148248 |
| IRS1 | 6.5144 | 0.009124667 | 2.039782997 |
| BBX | 8.3856 | 4.99E-05 | 4.30147912 |
| PCF11 | 8.3101 | 0.011852 | 1.926196973 |
| RGPD3 | 7.7254 | 0.00798 | 2.098015503 |

**Supplementary Table 7:** Proteins identified as TMZ-regulated PIMREG interactors in T98G cells (TMZ vs. DMSO - PIMREG IPs)

| **Gene name** | **log2 FC** | **Adj. *p*-value** | **-log10 *p*-value** |
|---|---:|---:|---:|
| YY1; ZFP42; YY2 | 10.14025 | 0.037689 | 1.423784134 |
| RPS27L | 13.64826 | 0.037689 | 1.423784134 |

**Supplementary Table 8:** Gene ontology annotations of TMZ-regulated PIMREG interacting proteins in U87MG cells (TMZ vs. DMSO - PIMREG IPs).

| Ontological term | Adj. *p*-value | Genes | Classification (BP) |
|---|---|---|---|
| Mitochondrial Electron Transport. NADH To Ubiquinone (GO:0006120) | 0.048259 | NDUFA8; NDUFS6 | Metabolism |
| Spindle Assembly Checkpoint Signaling (GO:0071173) | 0.038592 | ZW10; TRIP13 | Cell Division/ proliferation |
| Mitotic Spindle Assembly Checkpoint Signaling (GO:0007094) | 0.038592 | ZW10; TRIP13 | Cell Division/ proliferation |
| Mitotic Spindle Checkpoint Signaling (GO:0071174) | 0.038592 | ZW10; TRIP13 | Cell Division/ proliferation |
| Negative Regulation Of Mitotic Metaphase/Anaphase Transition (GO:0045841) | 0.038592 | ZW10; TRIP13 | Cell Division/ proliferation |
| Positive Regulation Of RNA Splicing (GO:0033120) | 0.038592 | PRMT5; NCBP1; RBM22 | mRNA splicing/ processing |
| Regulation Of mRNA Splicing. Via Spliceosome (GO:0048024) | 0.038592 | PRMT5; TIA1; NCBP1 | RNA splicing |
| Positive Regulation Of mRNA Splicing. Via Spliceosome (GO:0048026) | 0.038592 | PRMT5; NCBP1 | RNA splicing |
| Positive Regulation Of mRNA Processing (GO:0050685) | 0.038592 | PRMT5; NCBP1 | mRNA splicing/ processing |
| Regulation Of mRNA Processing (GO:0050684) | 0.048259 | TIA1; NCBP1 | mRNA splicing/ processing |

**Supplementary Table 9:** Gene ontology annotations of PIMREG interacting proteins in T98G cells (PIMREG IP vs. IgG under vehicle-DMSO conditions)

| **Ontological term** | **Adj. *p*-value** | **Genes** | **Classification (BP)** |
|---|---|---|---|
| Co-Transcriptional mRNA 3'-End Processing, Cleavage and Polyadenylation Pathway (GO:0180010) | 0.02392 | PCF11 | Transcription/mRNA processing |
| Positive Regulation of Fatty Acid Beta-Oxidation (GO:0032000) | 0.02392 | IRS1 | Metabolism related to insulin pathway |
| Co-Transcriptional RNA 3'-End Processing, Cleavage and Polyadenylation Pathway (GO:0180012) | 0.02392 | PCF11 | Transcription/mRNA processing |
| Insulin-Like Growth Factor Receptor Signaling Pathway (GO:0048009) | 0.02392 | IRS1 | Metabolism related to insulin pathway |
| Positive Regulation of Fatty Acid Oxidation (GO:0046321) | 0.02392 | IRS1 | Metabolism related to insulin pathway |
| Positive Regulation of Insulin Receptor Signaling Pathway (GO:0046628) | 0.02392 | IRS1 | Metabolism related to insulin pathway |
| Termination of RNA Polymerase II Transcription (GO:0006369) | 0.02392 | PCF11 | Transcription/mRNA processing |
| Positive Regulation of Glycogen Biosynthetic Process (GO:0045725) | 0.02392 | IRS1 | Metabolism related to insulin pathway |
| Positive Regulation of Cellular Response to Insulin Stimulus (GO:1900078) | 0.02392 | IRS1 | Metabolism related to insulin pathway |
| Regulation of Fatty Acid Beta-Oxidation (GO:0031998) | 0.02392 | IRS1 | Metabolism related to insulin pathway |
| Positive Regulation of Glycogen Metabolic Process (GO:0070875) | 0.02392 | IRS1 | Metabolism related to insulin pathway |
| Positive Regulation of Glucose Metabolic Process (GO:0010907) | 0.02392 | IRS1 | Metabolism related to insulin pathway |
| NLS-bearing Protein Import Into Nucleus (GO:0006607) | 0.02392 | RGPD3 | Protein transport |
| Negative Regulation of Insulin Secretion (GO:0046676) | 0.02392 | IRS1 | Metabolism related to insulin pathway |
| Immunoglobulin Mediated Immune Response (GO:0016064) | 0.02392 | IGLL5 | Immune response |
| Positive Regulation of Small Molecule Metabolic Process (GO:0062013) | 0.02392 | IRS1 | Metabolism related to insulin pathway |
| Negative Regulation of Peptide Hormone Secretion (GO:0090278) | 0.02392 | IRS1 | Metabolism related to insulin pathway |
| Response to Fatty Acid (GO:0070542) | 0.02392 | IRS1 | Metabolism related to insulin pathway |
| Cellular Response to Fatty Acid (GO:0071398) | 0.02392 | IRS1 | Metabolism related to insulin pathway |
| B Cell Mediated Immunity (GO:0019724) | 0.02392 | IGLL5 | Immune response |
| DNA-templated Transcription Termination (GO:0006353) | 0.02392 | PCF11 | Transcription/mRNA processing |
| Positive Regulation of Lipid Catabolic Process (GO:0050996) | 0.02397 | IRS1 | Metabolism related to insulin pathway |

| | | | |
|---|---|---|---|
| Positive Regulation of D-glucose Import (GO:0046326) | 0.02701 | IRS1 | Metabolism related to insulin pathway |
| Regulation of Glycogen Biosynthetic Process (GO:0005979) | 0.02701 | IRS1 | Metabolism related to insulin pathway |
| mRNA 3'-End Processing (GO:0031124) | 0.02701 | PCF11 | Transcription/mRNA processing |
| Regulation of Glucose Metabolic Process (GO:0010906) | 0.02701 | IRS1 | Metabolism related to insulin pathway |
| Negative Regulation of Insulin Receptor Signaling Pathway (GO:0046627) | 0.027656 | IRS1 | Metabolism related to insulin pathway |
| Positive Regulation of D-glucose Transmembrane Transport (GO:0010828) | 0.027656 | IRS1 | Metabolism related to insulin pathway |
| Negative Regulation of Cellular Response to Insulin Stimulus (GO:1900077) | 0.027656 | IRS1 | Metabolism related to insulin pathway |
| Positive Regulation of Carbohydrate Metabolic Process (GO:0045913) | 0.028395 | IRS1 | Metabolism related to insulin pathway |
| Regulation of D-glucose Import (GO:0046324) | 0.029086 | IRS1 | Metabolism related to insulin pathway |
| Phosphatidylinositol 3-Kinase/Protein Kinase B Signal Transduction (GO:0043491) | 0.030338 | IRS1 | Cell signaling |
| Negative Regulation of Protein Secretion (GO:0050709) | 0.030338 | IRS1 | Protein transport |
| Regulation of Insulin Receptor Signaling Pathway (GO:0046626) | 0.036014 | IRS1 | Metabolism related to insulin pathway |
| Insulin Receptor Signaling Pathway (GO:0008286) | 0.0364 | IRS1 | Metabolism related to insulin pathway |
| Response to Peptide Hormone (GO:0043434) | 0.038824 | IRS1 | Metabolism related to insulin pathway |

**Supplementary Table 10:** Gene ontology annotations of TMZ-regulated PIMREG interacting proteins in T98G cells (TMZ vs. DMSO - PIMREG IPs).

| Ontological term | Adj. *p*-value | Genes | Classification (BP) |
|---|---|---|---|
| Positive Regulation Of Telomere Maintenance (GO:0032206) | 0.0349807484272702 | YY1 | Telomere maintenance |
| Telomere Organization (GO:0032200) | 0.035307783998499 | YY1 | Telomere maintenance |
| Telomere Maintenance (GO:0000723) | 0.035307783998499 | YY1 | Telomere maintenance |
| Regulation Of DNA Strand Elongation (GO:0060382) | 0.0266160850218898 | YY1 | DNA Replication |
| Regulation Of DNA Replication (GO:0006275) | 0.0372746743314896 | YY1 | DNA Replication |
| Ribosomal Small Subunit Assembly (GO:0000028) | 0.0266160850218898 | RPS27L | Ribosome biogenesis |
| Ribosome Assembly (GO:0042255) | 0.0349807484272702 | RPS27L | Ribosome biogenesis |
| Ribosomal Small Subunit Biogenesis (GO:0042274) | 0.0372813634696548 | RPS27L | Ribosome biogenesis |
| Response To UV-C (GO:0010225) | 0.0246202632327964 | YY1 | DNA Damage Response |
| DNA Damage Response. Signal Transduction By P53 Class Mediator Resulting In Transcription Of P21 Class Mediator (GO:0006978) | 0.0246202632327964 | RPS27L | DNA Damage Response |
| DNA Damage Response. Signal Transduction Resulting In Transcription (GO:0042772) | 0.0246202632327964 | RPS27L | DNA Damage Response |
| Positive Regulation Of Telomere Maintenance In Response To DNA Damage (GO:1904507) | 0.0266160850218898 | YY1 | DNA Damage Response |
| Regulation Of Telomere Maintenance In Response To DNA Damage (GO:1904505) | 0.0266160850218898 | YY1 | DNA Damage Response |
| Intrinsic Apoptotic Signaling Pathway In Response To DNA Damage By P53 Class Mediator (GO:0042771) | 0.0272630017540475 | RPS27L | DNA Damage Response |
| Mitotic G1 DNA Damage Checkpoint Signaling (GO:0031571) | 0.0344134281935479 | RPS27L | DNA Damage Response |
| DNA Damage Response. Signal Transduction By P53 Class Mediator (GO:0030330) | 0.0349807484272702 | RPS27L | DNA Damage Response |
| Intrinsic Apoptotic Signaling Pathway In Response To DNA Damage (GO:0008630) | 0.0349807484272702 | RPS27L | DNA Damage Response |
| Mitotic DNA Damage Checkpoint Signaling (GO:0044773) | 0.035307783998499 | RPS27L | DNA Damage Response |
| Cellular Response To UV (GO:0034644) | 0.0372746743314896 | YY1 | DNA Damage Response |
| Positive Regulation Of DNA Repair (GO:0045739) | 0.0380446698343697 | YY1 | DNA Damage Response |
| Recombinational Repair (GO:0000725) | 0.0394539357720813 | YY1 | DNA Damage Response |

| | | | |
|---|---|---|---|
| Response To UV (GO:0009411) | 0.0394539357720813 | YY1 | DNA Damage Response |
| Regulation Of Cellular Response To Stress (GO:0080135) | 0.0394539357720813 | YY1 | DNA Damage Response |
| Double-Strand Break Repair Via Homologous Recombination (GO:0000724) | 0.0394539357720813 | YY1 | DNA Damage Response |
| Regulation Of DNA Repair (GO:0006282) | 0.0430771077386107 | YY1 | DNA Damage Response |
| Female Gonad Development (GO:0008585) | 0.0246202632327964 | ZFP42 | Cell differentiation/ development |
| Development Of Primary Female Sexual Characteristics (GO:0046545) | 0.0246202632327964 | ZFP42 | Cell differentiation/ development |
| Development Of Primary Male Sexual Characteristics (GO:0046546) | 0.0349807484272702 | ZFP42 | Cell differentiation/ development |
| Male Gonad Development (GO:0008584) | 0.0349807484272702 | ZFP42 | Cell differentiation/ development |
| Gonad Development (GO:0008406) | 0.0349807484272702 | ZFP42 | Cell differentiation/ development |
| Regulation Of Multicellular Organismal Development (GO:2000026) | 0.035307783998499 | YY1 | Cell differentiation/ development |
| Regulation Of Embryonic Development (GO:0045995) | 0.035307783998499 | YY1 | Cell differentiation/ development |
| Negative Regulation Of Interferon-Beta Production (GO:0032688) | 0.0246202632327964 | YY1 | INF signaling |
| Negative Regulation Of Type I Interferon Production (GO:0032480) | 0.0349807484272702 | YY1 | INF signaling |
| Regulation Of Interferon-Beta Production (GO:0032648) | 0.0349807484272702 | YY1 | INF signaling |
| Mitotic G1/S Transition Checkpoint Signaling (GO:0044819) | 0.0246202632327964 | RPS27L | Cell division |
| Regulation Of Chromosome Organization (GO:0033044) | 0.0349807484272702 | YY1 | Cell division |
| Intrinsic Apoptotic Signaling Pathway By P53 Class Mediator (GO:0072332) | 0.0349807484272702 | RPS27L | Apoptosis |
| Activation Of Cysteine-Type Endopeptidase Activity Involved In Apoptotic Process (GO:0006919) | 0.035307783998499 | RPS27L | Apoptosis |
| Positive Regulation Of Cysteine-Type Endopeptidase Activity Involved In Apoptotic Process (GO:0043280) | 0.0394539357720813 | RPS27L | Apoptosis |
| Positive Regulation Of Amide Metabolic Process (GO:0034250) | 0.0372813634696548 | RPS27L | Macromolecule Metabolic Process |
| Positive Regulation Of DNA Metabolic Process (GO:0051054) | 0.0394539357720813 | YY1 | Macromolecule Metabolic Process |
| Positive Regulation Of Macromolecule Biosynthetic Process (GO:0010557) | 0.0422567210750957 | RPS27L | Macromolecule Metabolic Process |
| Regulation Of DNA Metabolic Process (GO:0051052) | 0.0422567210750957 | YY1 | Macromolecule Metabolic Process |

| | | | |
|---|---|---|---|
| Regulation Of DNA-templated Transcription (GO:0006355) | 0.0349807484272702 | YY1; YY2; ZFP42 | Transcription/Translation |
| Regulation Of Transcription By RNA Polymerase II (GO:0006357) | 0.0349807484272702 | YY1; YY2; ZFP42 | Transcription/Translation |
| Positive Regulation Of Transcription By RNA Polymerase II (GO:0045944) | 0.0372746743314896 | YY1; YY2 | Transcription/Translation |
| Positive Regulation Of Translation (GO:0045727) | 0.0394539357720813 | RPS27L | Transcription/Translation |
| Positive Regulation Of DNA-templated Transcription (GO:0045893) | 0.0451493356324105 | YY1; YY2 | Transcription/Translation |
| Protein-RNA Complex Assembly (GO:0022618) | 0.0480983855536169 | RPS27L | Transcription/Translation |
